# Arsenic-sensing domain controls ACR3 transporter trafficking and function in *Marchantia polymorpha*

**DOI:** 10.1101/2025.04.13.648538

**Authors:** Katarzyna Mizio, Ignacy Bonter, Kacper Zbieralski, Alicja Dolzblasz, Paulina Tomaszewska, Jacek Staszewski, Donata Wawrzycka, Anna Reymer, Wojciech Bialek, Verena Kriechbaumer, Jim Haseloff, Robert Wysocki, Ewa Maciaszczyk-Dziubinska

## Abstract

Arsenic, a toxic and carcinogenic metalloid, is a pervasive environmental contaminant that threatens human health through contaminated water and food. The efflux of As(III) via ACR3 transporters is an ancient detoxification mechanism conserved across prokaryotes, fungi, and plants, with the notable exception of angiosperms. Despite their evolutionary significance, plant ACR3s remain largely uncharacterized. Here, we demonstrate that MpACR3, the ACR3 orthologue from the liverwort *Marchantia polymorpha*, functions as a metalloid/proton antiporter, conferring resistance to arsenicals and moderate tolerance to antimony. Additionally, we uncover an arsenic-sensing domain within MpACR3 that regulates its intracellular trafficking. Under normal conditions, MpACR3 sorting to the plasma membrane is delayed, resulting in its retention within Golgi bodies. However, As(III) binding to three cysteine residues in the N-terminal cytosolic domain induces a conformational change that facilitates MpACR3 trafficking to the plasma membrane. Furthermore, mutational analysis of a conserved arginine-based motif reveals that the N-terminal domain not only controls MpACR3 accumulation at the plasma membrane but also modulates its transport activity. Importantly, this arsenic-sensing domain is conserved among plant ACR3 transporters, suggesting a plant-specific adaptation to arsenic toxicity.

## Introduction

Arsenic is a toxic element that occurs naturally in the environment and which, as a result of geogenic and anthropogenic activities, can reach dangerously high concentrations in certain areas (Oremland and Stolz 2003; Wysocki et al. 2023). Arsenic contamination of groundwater has been identified in more than 100 countries, posing a potential health risk to up to 220 million people worldwide (Rahaman et al. 2021). The primary sources of human exposure to arsenic are drinking water (Podgorski and Berg 2020) and food products such as cereals, particularly rice and wheat-based products, fruit and fruit products, vegetables, seafood, and animal products (Zhao et al. 2024). Epidemiological studies have consistently shown that chronic exposure to arsenic is associated with various health issues in humans, including an increased risk of coronary heart disease and several cancers (Naujokas et al. 2013; Oberoi et al. 2019). Hence, the International Agency for Research on Cancer (IARC), a member of the World Health Organization (WHO), has classified arsenic as a human carcinogen. Arsenic is also number one on the U.S. Agency for Toxic Substances and Disease Registry (ATSDR) list of priority substances posing the greatest potential risk to human health.

Arsenite [As(OH)_3_ or As(III)] and arsenate [H_2_AsO_4_^-^/HAsO_4_^2-^ or As(V)] are the most prevalent forms of inorganic arsenic in water and soil (Jain and Ali 2000). Due to molecular mimicry, As(III) and As(V) are readily absorbed by cells across all organisms via aquaglyceroporin channels and phosphate uptake transporters, respectively (Maciaszczyk-Dziubinska et al. 2012; Garbinski et al. 2019). As(III) efflux transporters of the ACR3 family, which confer high-level resistance to As(III) and As(V), are widely distributed across bacteria, archaea, and most eukaryotic lineages, highlighting a significant evolutionary adaptation to environmental arsenic exposure (Indriolo et al. 2010; Maciaszczyk-Dziubinska et al. 2012; Garbinski et al. 2019; Chen et al. 2020). In eukaryotes, arsenic detoxification also involves the formation of As(III)-thiol conjugates through complexation with cysteine-containing peptides such as glutathione and phytochelatins. These conjugates are subsequently sequestered in the vacuole or exported from the cell by ATP-binding cassette (ABC) transporters, a crucial detoxification pathway in organisms lacking ACR3 transporters, including flowering plants and animals (Ghosh et al. 1999; Vatamaniuk et al. 2001; Song et al. 2010; Leslie 2012; Song et al. 2014). Additionally, As(III) can diffuse out of the cytosol down its concentration gradient through aquaglyceroporins located in the plasma membrane (PM) or vacuolar membrane (Bienert et al. 2008; Maciaszczyk-Dziubinska et al. 2010a; Pommerrenig et al. 2015; Karle and Kumar 2024). In plants, the OsABCC7 and Lsi2 transporters in rice (*Oryza sativa*) and the aquaglyceroporins NIP3;1 and NIP7;1 in *Arabidopsis thaliana* have been shown to facilitate As(III) root-to-shoot translocation, contributing to arsenic accumulation in shoots (Ma et al. 2008; Xu et al. 2015; Lindsay and Maathuis 2016; Tang et al. 2019).

ACR3 transporters have been well-characterized in bacteria and budding yeast, where they localize to the PM and facilitate As(III) extrusion (Bobrowicz et al. 1997; Wysocki et al. 1997; Sato and Kobayashi 1998; Ghosh et al. 1999; Fu et al. 2009; Maciaszczyk-Dziubinska et al. 2010b; 2011; Villadangos et al. 2012; Mizio et al. 2023). These proteins share key structural and functional similarities, including a topology of ten transmembrane regions, As(III)/H⁺ antiporter activity, and a dependence on highly conserved cysteine and glutamic acid residues for transport function (Maciaszczyk-Dziubinska et al. 2011; Villadangos et al. 2012). The *Saccharomyces cerevisiae* ACR3 (ScAcr3) can also transport the toxic metalloid antimony [Sb(III)], albeit at a significantly slower rate than As(III) (Maciaszczyk-Dziubinska et al. 2010b; 2011). This suggests a broader substrate specificity than that observed in most bacterial ACR3 transporters studied to date (Sato and Kobayashi 1998; Fu et al. 2009; Villadangos et al. 2012). Notably, ACR3 proteins from the cyanobacterium *Synechocystis* sp. (López-Maury et al. 2003), the campylobacterium *Brevundimonas nasdae* (Yang et al. 2022), and the rhizobacterium *Agrobacterium tumefaciens* (Kang et al. 2015) have been shown to confer both As(III) and Sb(III) tolerance. Furthermore, a single amino acid substitution (E353D) converts ScAcr3 into a Sb(III)-specific efflux transporter (Mizio et al. 2023).

Expression of the *ScACR3* gene in *A. thaliana* and *O. sativa*, both naturally lacking *ACR3*, enhanced tolerance to As(III) and As(V) while promoting increased arsenic efflux from roots to the external environment (Ali et al. 2012; Duan et al. 2012). However, while *A. thaliana* plants expressing *ScACR3* did not show reduced arsenic levels in their shoots (Ali et al. 2012), another study reported that *ScACR3* expression significantly lowered arsenic accumulation in rice straw and grains (Duan et al. 2012). Beyond *ScACR3*, the only other eukaryotic orthologues analysed to date are from the fern genus *Pteris* (Indriolo et al. 2010; Chen et al. 2021; Popov et al. 2021; Sun et al. 2023), and, more recently, from the model bryophyte *Marchantia polymorpha* (Dutta et al. 2024; Li et al. 2024). Noteworthy, the model green algae *Chlamydomonas reinhardtii*, which lacks an *ACR3* orthologue, is ten times more sensitive to As(V) than the acidophilic green algae *C. eustigma*, which expresses *ACR3* (Hirooka et al. 2017). This suggests that the *C. eustigma ACR3* gene (*CeACR3*) encodes a functional arsenic transporter.

*Pteris vittata* is a well-characterized arsenic hyperaccumulator that expresses up to five *ACR3* paralogues: *PvACR3*, *PvACR3;1*, *PvACR3;2*, *PvACR3;3*, and *PvACR3;4* (Indriolo et al. 2010; Chen et al. 2017; Chen et al. 2021; Sun et al. 2023). Heterologous overexpression of *P*. *vittata ACR3* genes in an *S*. *cerevisiae acr3*Δ strain (lacking the endogenous *ACR3* gene) and in *A*. *thaliana* or *Nicotiana tabacum* has demonstrated that these transporters enhance tolerance to both As(III) and As(V) (Indriolo et al. 2010; Chen et al. 2013; Chen et al. 2017; Wang et al. 2018; Chen et al. 2021). Moreover, silencing of the *PvACR3* gene, but not the *PvACR3;1* gene in *P. vittata* gametophytes resulted in increased sensitivity to As(III) (Indriolo et al. 2010). Interestingly, *P. vittata* ACR3 proteins exhibit a distinctive subcellular localisation pattern, which likely contributes to the species’ arsenic hyperaccumulation capability. PvACR3 was found to localize to the vacuolar membrane in *P*. *vittata* gametophytes (Indriolo et al. 2010). Similarly, when expressed in *A. thaliana* and *N. tabacum*, PvACR3;1 and PvACR3;3 localized to the vacuolar membrane, facilitating As(III) sequestration in the vacuole and its retention in root cells (Chen et al. 2017; 2021). In contrast, PvACR3 and PvACR3;2 were targeted to the PM in transgenic plants, where they mediated As(III) extrusion from root cells into the external environment while simultaneously promoting As(III) translocation from roots to shoots (Chen et al. 2013; Wang et al. 2018; Chen et al. 2021).

As a liverwort, *M. polymorpha* holds a pivotal evolutionary position among land plants. Bryophytes have been identified as a monophyletic group, sister to extant tracheophytes, making them valuable for uncovering conserved mechanisms inherited from the common ancestor of all land plants (Harris et al. 2022). In *M. polymorpha*, the Mp*ACR3* orthologue plays a key role in arsenic detoxification, as evidenced by loss-of-function mutants that exhibit hypersensitivity to As(III) and As(V), along with elevated arsenic accumulation. In contrast, Mp*ACR3*-overexpressing transgenic lines display enhanced tolerance and lower arsenic levels in plant tissues compared to wild-type plants (Li et al. 2024). Consistent with its function, the MpACR3-Citrine fusion protein was localized to the PM in *M. polymorpha* cells. Interestingly, MpACR3-Citrine signals were also detected in unidentified intracellular bodies, possibly indicating folding or sorting defects associated with the overexpression of the fusion protein (Li et al. 2024). Notably, the ability of MpACR3 and PvACR3 proteins to confer resistance to Sb(III) has yet to be investigated.

In this study, we conducted a comprehensive characterization of the *M. polymorpha* ACR3 transporter. First, we experimentally demonstrate that, similar to yeast and bacterial ACR3 proteins, MpACR3 functions as a plasma membrane As(III)/H⁺ antiporter, conferring tolerance to both trivalent and pentavalent arsenic. Additionally, we pinpoint key amino acid residues essential for MpACR3 transport activity. Second, we show that, like the *S*. *cerevisiae* ACR3, MpACR3 exhibits limited capacity to transport Sb(III) and provides only low-level resistance to this metalloid. Third, heterologous expression of Mp*ACR3 in A. thaliana* suggests its potential application in developing arsenic-resistant crop plants, as its expression leads to reduced arsenic accumulation in plant tissues. Finally, we identify a plant-specific arsenic-sensing domain within the cytosolic N-terminal region of MpACR3. This domain regulates MpACR3 trafficking to the PM and modulates its transport function in response to arsenic exposure. Our findings also suggest that arsenic binding triggers a conformational change in the sensor, allowing MpACR3 to be released from intracellular retention and enhancing its transport activity. This discovery underscores the evolutionary adaptation of plant ACR3 transporters and provides new insights into the cellular mechanisms underlying arsenic sensing and detoxification in plants.

## Results

### Plant ACR3 transporters and the predicted structure of MpACR3

The phylogenetic analysis of plant ACR3 transporters reveals two distinct clades, corresponding to algal and land plant ACR3 proteins, respectively, which aligns with the current understanding of evolutionary history of plants. Strikingly, an exception is the ACR3 from the lycophyte *Selaginella moellendorffii* (SmACR3), that is positioned at the base of the land plant clade (Supplementary Fig. S1; Supplementary Table S1). ACR3 orthologues are widely distributed across chlorophytes, bryophytes, lycophytes, ferns, and gymnosperms (Supplementary Fig. S1; Supplementary Table S1). In addition to the fern *P. vittata,* which expresses five ACR3 paralogues, two or three copies of the *ACR3* gene are encoded in the genomes of several other species, including the fern *Marsilea vestita*, the lycophytes *Diphasiastrum complanatum* and *Isoetes taiwanensis*, the charophyte *Klebsormidium nitens*, and the chlorophytes *Coccomyxa elongate* and *Coccomyxa viridis* (Supplementary Table S1). Conversely, *ACR3* orthologues are absent in the prasinodermophyte *Prasinoderma coloniale*, several species of chlorophyte green algae, including the model algae *Chlamydomonas reinhardtii* and *Volvox carteri*, as well as the charophyte green algae *Chara braunii*, *Chlorokybus atmophyticus*, and *Mesostigma viride*. Additionally, *ACR3* genes are notably absent in angiosperms (Indriolo et al. 2010).

The *M. polymorpha* genome encodes a single *ACR3* gene (Mp*ACR3*, Mp5g06690), which translates into a 457-amino-acid protein with a predicted ten-transmembrane-helix (TM1-10) topology, featuring both N- and C-termini in the cytoplasm (Supplementary Figs. S2 and S3). The MpACR3 protein shares 44.3%, 40.5%, 66.3% sequence identity with previously characterized ACR3 transporters from the bacterium *Corynebacterium glutamicum* (CgAcr3), the yeast *S*. *cerevisiae* (ScACR3), and the fern *P*. *vittata* (PvACR3), respectively (Supplementary Fig. S2). Consistent with other ACR3 family members (Lv et al. 2022; Mizio et al. 2023), the AlphaFold-predicted structure of MpACR3 (Entry ID A0A176WR38) (Jumper et al. 2021) comprises two distinct domains: a panel domain (formed by α-helices TM1-2 and TM6-7) and a transport domain (α-helices TM3-5 and TM8-10), which is characterized by a crossover region formed by discontinuous α-helices TM4 and TM9 (Supplementary Fig. S3). A notable feature of MpACR3 is its unusually long N-terminal tail of 123 amino acids (Supplementary Fig. S2), predicted to form a cytosolic domain with an antiparallel β-barrel motif (Supplementary Fig. S3). Interestingly, such N-terminal extensions (104-163 residues) are a common characteristic of plant ACR3 orthologues (Supplementary Fig. S4). Only a few ACR3 sequences, primarily from ferns and certain algae, contain N-terminal tails shorter than 100 residues (Supplementary Fig. S5).

### MpACR3 enhances yeast tolerance to arsenic and antimony

To investigate the function of MpACR3, we employed heterologous expression in *S*. *cerevisiae acr3*Δ mutant cells, which are sensitive to arsenic and antimony due to the absence of an endogenous *ACR3* gene (Wysocki et al. 1997; Maciaszczyk-Dziubinska et al. 2010; 2011). The Mp*ACR3* cDNA, isolated from the *M*. *polymorpha* Cam-1 gametophyte, was cloned under the control of the strong *MET17* promoter and fused to a C-terminal GFP reporter. The *acr3*Δ strain was then transformed with pMpACR3, expressing the MpACR3-GFP fusion protein, pScACR3, a positive control expressing the *S*. *cerevisiae ACR3-GFP* fusion gene under the same *MET17* promoter, or pUG35 (empty vector) as a negative control. Western blot analysis confirmed that both MpACR3-GFP and ScACR3-GFP were highly expressed in the transformed yeast strains (Fig. 1A). Notably, metalloid exposure did not affect the protein levels of MpACR3-GFP or ScACR3-GFP. Interestingly, treatment with As(III) and Sb(III) induced the appearance of a slowly migrating form of MpACR3-GFP, suggesting a potential post-translational modification or conformational change (Fig. 1A).

**Figure 1.**
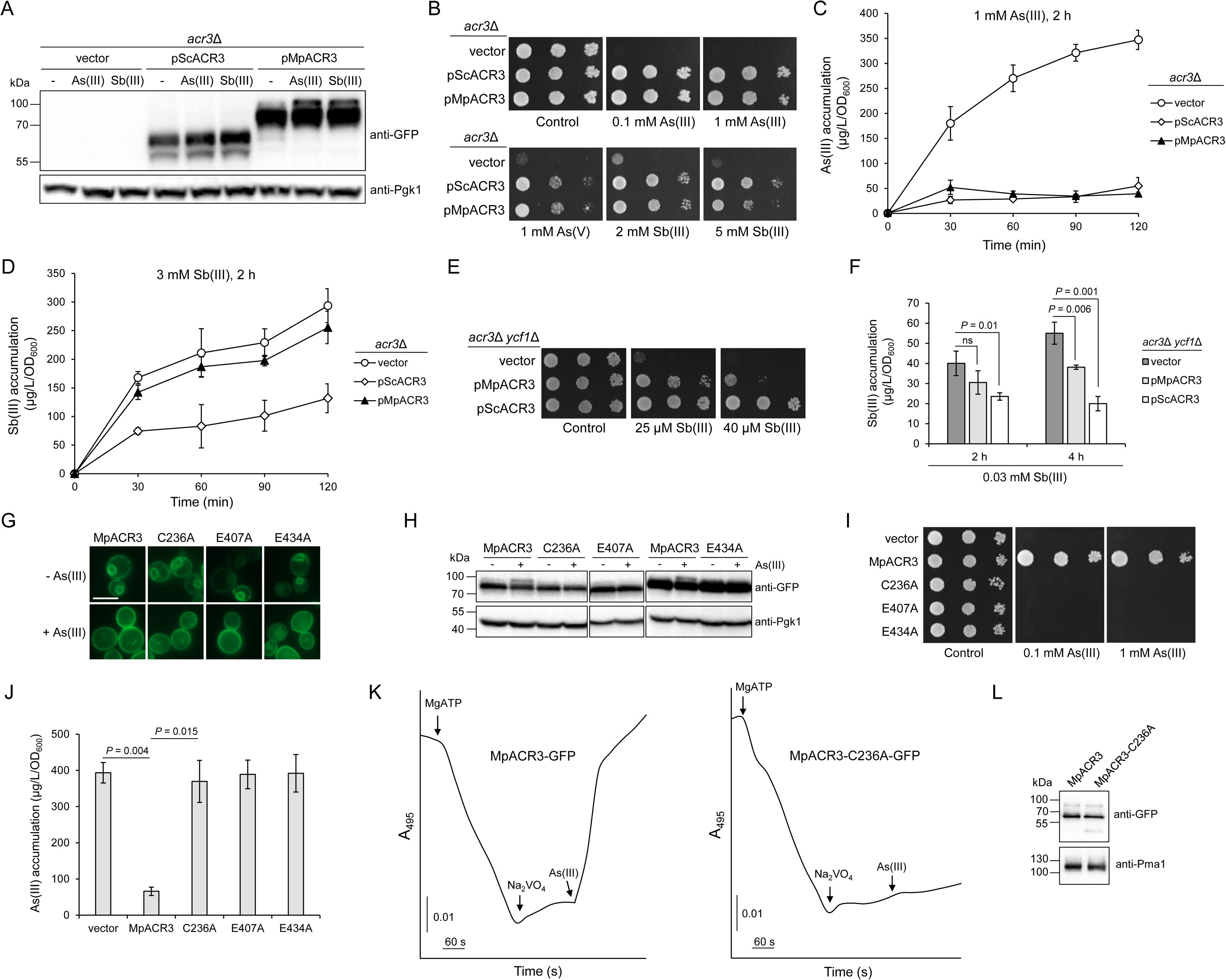
Functional analysis of Mp*ACR3* in *S*. *cerevisiae*. **A** to **D)** Heterologous expression of Mp*ACR3* restores arsenic and antimony resistance in yeast cells lacking *ScACR3*. The *S*. *cerevisiae acr3*Δ strain was transformed with the following plasmids: pUG35 (empty vector), pScACR3, or pMpACR3, expressing *ScACR3-GFP* or Mp*ACR3-GFP* genes, respectively. **A)** Western blot analysis of ScAcr3-GFP and MpACR3-GFP fusion proteins. Cultures were exposed to 0.1 mM As(III), 1 mM Sb(III), or left untreated for 2 h before protein extraction. Pgk1 (3-phosphoglycerate kinase) served as a loading control. **B)** Growth assays showing that Mp*ACR3-GFP* expression rescues the sensitivity of *acr3*Δ cells to metalloids. **C** and **D)** Metalloid accumulation assays demonstrating that Mp*ACR3-GFP* expression significantly reduces As(III) accumulation in *acr3*Δ cells, whereas Sb(III) accumulation remains largely unchanged except at the 30-min time point. Error bars represent mean ± standard deviation (*n* = 3). *P*-values were calculated using one-way ANOVA: **C)** pScACR3 vs. vector, *P* = 0.002; pMpACR3 vs. vector, *P* = 0.005; pScACR3 vs. pMpACR3, not significant. **D)** pScACR3 vs. vector, *P* = 0.008; pMpACR3 vs. vector, not significant (except at 30-min, *P* = 0.02); pScACR3 vs. pMpACR3, *P* = 0.01. **E)** Mp*ACR3-GFP* partially complemented the hypersensitivity of *acr3*Δ *ycf1*Δ cells to Sb(III). **F)** Mp*ACR3-GFP* reduces Sb(III) accumulation in *acr3*Δ *ycf1*Δ cells exposed to low Sb(III) concentrations. Error bars represent mean ± standard deviation of mean (*n* = 3). *P*-values were calculated using one-way ANOVA; ns, not significant. **G** to **L)** MpACR3 is an As(III)/H^+^ antiporter that requires the conserved C236, E407, and E434 residues for transport activity. **G)** The localisation of MpACR3-GFP variants in *acr3*Δ cells was monitored under normal conditions or after exposure to 0.1 mM As(III) for 2 h by fluorescence microscopy. Scale bar = 5 μm. **H)** Western blot analysis of MpACR3-GFP variants expressed in *acr3*Δ cells. Yeast cultures were exposed to 0.1 mM As(III) or left untreated for 2 h before protein extraction. **I)** Expression of the indicated MpACR3 variants did not complement the sensitivity of *acr3*Δ cells to As(III). **J)** Expression of the indicated MpACR3 variants did not prevent As(III) accumulation in *acr3*Δ cells when exposed to 1 mM As(III) for 2 h. Error bars represent mean value ± standard deviation of mean (*n* = 3). One-way ANOVA was used to calculate the *P*-value. **K)** As(III)-induced H^+^ transport across the membranes of inside-out vesicles prepared from *acr3*Δ cells expressing Mp*ACR3-GFP* or the Mp*ACR3-C236A-GFP* mutant was monitored by measuring changes in the absorbance of acridine orange used as a pH-sensitive probe. Vesicle acidification was initiated by the addition of 2 mM of ATP (first arrow), accompanied by a decrease of absorbance. After 3 min, 0.5 mM of sodium orthovanadate was added to inhibit H^+^-ATPase and maintain a steady-state acidic-inside pH gradient (second arrow). At the indicated time point, 10 mM of As(III) was added to initiate As(III)-dependent H^+^ movement (third arrow), which acidified the environment and recovered the absorbance. **L)** Western blot analysis of MpACR3-GFP and Pma1 levels in the inside-out membrane vesicles prepared from *acr3*Δ cells expressing Mp*ACR3-GFP* or the Mp*ACR3-C236A-GFP* mutant. **K** and **L)** Transformant cultures were pre-treated with 0.1 mM As(III) to induce MpACR3-GFP accumulation in the PM.

Functional assays revealed that, similar to ScACR3, overexpression of Mp*ACR3-GFP* fully complemented the As(III) and As(V) sensitivity of *acr3*Δ cells, restoring growth to wild-type levels (Fig. 1B). However, in the presence of Sb(III), Mp*ACR3-GFP* only partially rescued *acr3*Δ growth, indicating a limited role in Sb(III) transport (Fig. 1B). Consistent with the growth assays, cells overexpressing Mp*ACR3-GFP* or *ScACR3-GFP* maintained low intracellular As(III) levels, supporting efficient As(III) extrusion. In contrast, *acr3*Δ cells harbouring the empty vector accumulated progressively higher levels of As(III) (Fig. 1C). While Sb(III) accumulation was significantly reduced in pScACR3 transformants (Fig. 1D), statistical analysis showed no significant difference in Sb(III) levels between pMpACR3 and empty vector samples, except at the 30-min time point (Fig. 1D). Nonetheless, in each biological replicate, *acr3*Δ cells expressing Mp*ACR3-GFP* exhibited a slight but consistent reduction in Sb(III) accumulation compared to the control vector (Supplementary Fig. S6). To further investigate the substrate specificity of MpACR3, we expressed Mp*ACR3-GFP* in the *acr3*Δ *ycf1*Δ double mutant, which lacks both the *ACR3* and *YCF1* genes. In yeast, Ycf1 plays a crucial role in Sb(III) detoxification by sequestering glutathione-conjugated Sb(III) into the vacuole (Ghosh et al. 1999). Consequently, the expression of Mp*ACR3-GFP* partially rescued the Sb(III) hypersensitivity of *acr3*Δ *ycf1*Δ cells (Fig. 1E) and significantly reduced Sb(III) accumulation when cells were exposed to low Sb(III) concentrations for over 2 h. However, this effect was less pronounced compared to *ScACR3* expression (Fig. 1F). Collectively, these results indicate that MpACR3 mediates an extrusion of both As(III) and Sb(III), although Sb(III) appears to be a much weaker substrate than As(III).

Next, we examined the subcellular localisation of the MpACR3-GFP fusion protein by observing distribution of its GFP signal using fluorescence microscopy. Supporting its role as an As(III) efflux transporter, MpACR3-GFP localized to the cell surface during As(III) treatment (Fig. 1G). Interestingly, under normal conditions, the MpACR3-GFP signal appeared as patches at the cell periphery and as a ring-shaped intracellular structure, likely corresponding to the cortical and perinuclear endoplasmic reticulum (ER), respectively (Fig. 1G) (Young et al. 2013). This suggests that in yeast, MpACR3 trafficking to the PM is regulated by As(III).

### MpACR3 employs a conserved mechanism for As(III) transport

Mutational analysis of bacterial and yeast *ACR3* genes has identified several highly conserved residues essential for arsenic transport activity. Specifically, a cysteine residue in TM4 (C129 in *C. glutamicum* Acr3 (CgAcr3) and C151 in ScAcr3) and a glutamic acid residue in TM9 (E305 in CgAcr3 and E353 in ScAcr3) are critical for function (Supplementary Fig. S2) (Fu et al. 2009; Villadangos et al. 2012; Maciaszczyk-Dziubinska et al. 2014; Markowska et al. 2015). Additionally, mutation of a conserved glutamic acid residue in TM10 (E380 in ScAcr3) abolished arsenic transport activity (Supplementary Fig. S2) (Markowska et al. 2015). However, mutating the corresponding residue in CgAcr3 (E332) had no significant impact on transport function (Villadangos et al. 2012).

Despite these insights, key residues required for the transport function of plant ACR3 proteins remain unexplored. To address this, we generated MpACR3 variants in which the corresponding conserved residues, i.e., Cys236, Glu407, and Glu434, were replaced with alanine. Although these mutants maintained wild-type protein levels (Fig. 1H) and exhibited As(III)-induced accumulation at the PM (Fig. 1G), they failed to complement the As(III) sensitivity of *S. cerevisiae acr3*Δ cells (Fig. 1I). Furthermore, yeast transformants expressing these MpACR3 variants accumulated high levels of As(III) (Fig. 1J), confirming that, as in ScAcr3, C236, E407, and E434 are essential for MpACR3 transport activity.

Both bacterial and yeast ACR3 transporters function as low-affinity As(III)/H^+^ antiporters (Maciaszczyk-Dziubinska et al. 2011; Villadangos et al. 2012). To investigate the energy-dependence of MpACR3-mediated As(III) transport, we prepared inside-out vesicles from *acr3*Δ cells expressing MpACR3-GFP. We then generated a proton gradient across the vesicle membranes, as indicated by a decrease in acridine orange absorbance (Fig. 1K). Upon the addition of 10 mM As(III), a complete recovery of absorbance was observed, reflecting solution acidification and confirming H^+^ efflux from the vesicles (Fig. 1K). In contrast, no As(III)-induced movement of protons was detected in vesicles containing the transport-inactive MpACR3-C236A-GFP variant (Fig. 1K). Importantly, membrane vesicle fractions showed comparable levels of MpACR3-GFP and MpACR3-C236A-GFP proteins (Fig. 1L), ruling out differences in protein expression as a factor. These results demonstrate that MpACR3 catalyses As(III)/ H^+^ exchange across the PM.

ScAcr3 also functions as an Sb(III)/H^+^ antiporter but transports Sb(III) at a much slower rate than As(III) (Maciaszczyk-Dziubinska et al. 2011). We therefore tested whether MpACR3 could mediate Sb(III)-induced H^+^ transport. Using everted vesicles from *acr3*Δ cells expressing MpACR3-GFP, we found that adding 10 mM Sb(III) did not induce H^+^ efflux (Supplementary Fig. S7A). However, at a higher concentration (20 mM), Sb(III) partially dissipated the pH gradient, confirming that Sb(III) is a poor substrate of MpACR3 (Supplementary Fig. S7B).

### Arsenic and antimony trigger MpACR3 relocation from the ER to the PM in yeast

In the fern *P. vittata*, ACR3 paralogues localize to either the PM or the vacuole membrane (Indriolo et al. 2010; Chen et al. 2013; 2017; 2021; Wang et al. 2018). In contrast, recent findings indicate that MpACR3 localizes to the PM and unspecified endomembrane systems in *M*. *polymorpha* cells (Li et al. 2024). Here, we found that in the absence of As(III), MpACR3-GFP in *S. cerevisiae* also exhibits an intracellular localisation pattern consistent with the ER (Fig. 1G).

To further investigate its subcellular localisation, we stained MpACR3-GFP transformant cells with FM4-64, a fluorescent marker that selectively binds to vacuolar membranes in yeast. The MpACR3-GFP signal did not align with the vacuole periphery, which is morphologically distinctive in differential interference contrast (DIC) microscopy, nor did it co-localize with FM4-64-labelled vacuolar membranes (Fig. 2A). To confirm the ER localisation of MpACR3-GFP, we next employed the *acr3*Δ strain expressing mCherry-HDEL, a red fluorescent ER marker containing the HDEL retention signal (Zhu et al. 2019). Fluorescence microscopy revealed that MpACR3-GFP co-localized with mCherry-HDEL, confirming its presence in the ER (Fig. 2B). As observed in Fig. 1G, after 2 h of exposure to 0.1 mM As(III), the ER-associated MpACR3-GFP signal disappeared, while the GFP signal at the cell surface became more uniform (Fig. 2B). This strongly suggests that under normal conditions, MpACR3-GFP is retained in the ER but relocates to the PM in response to As(III) (Fig. 2B).

**Figure 2.**
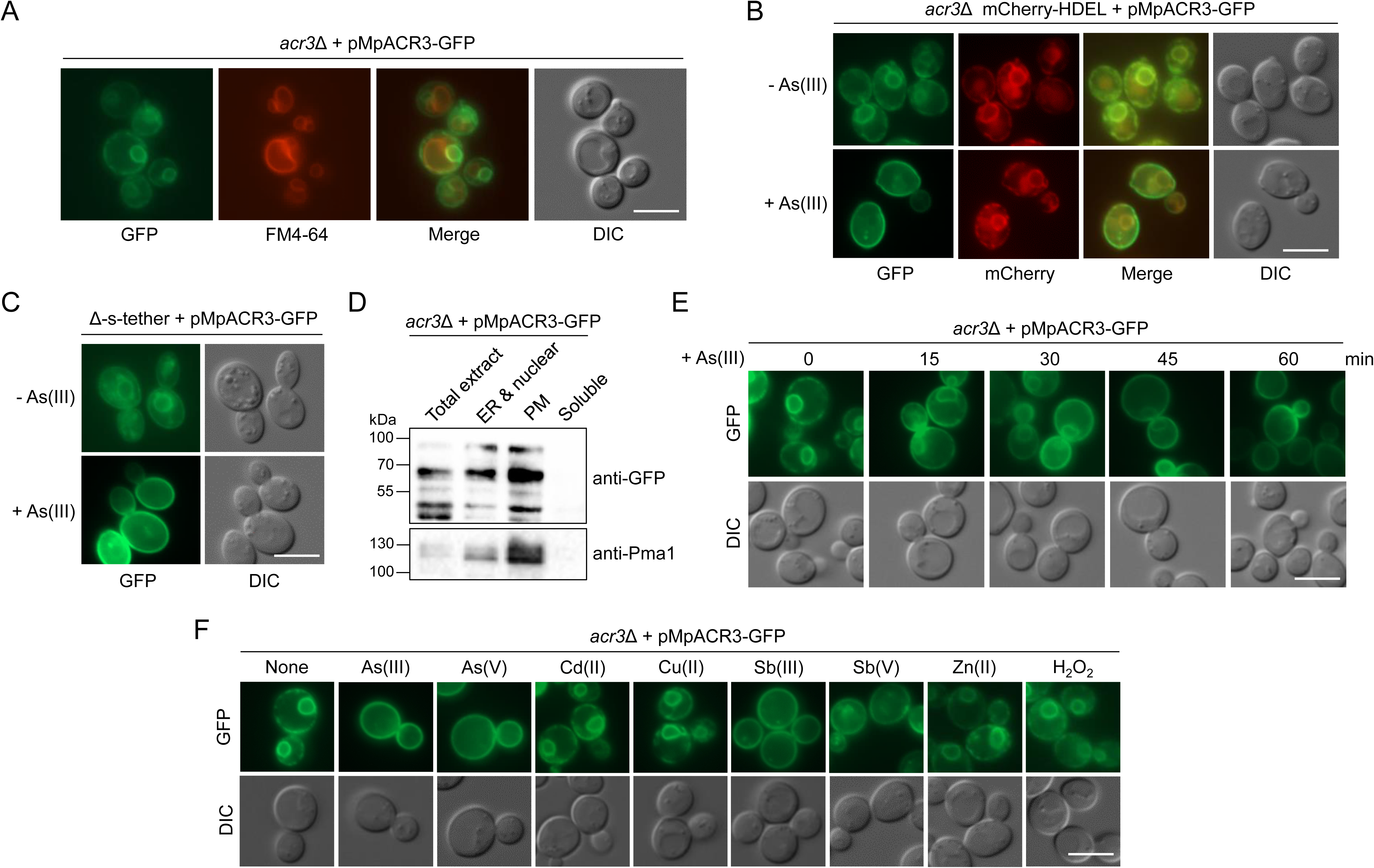
MpACR3 is retained in the ER under non-stress conditions in yeast cells and is relocated to the PM in response to arsenic and antimony. **A)** Under normal conditions, MpACR3-GFP exhibits an intracellular localisation that is distinct from vacuolar membranes. Yeast vacuolar membranes were visualized by staining with the FM4-64 dye. **B)** MpACR3-GFP co-localized with the mCherry-HDEL ER marker and re-localized to the PM in response to As(III). Yeast cells were treated with 0.1 mM As(III) for 2 h or left untreated before microscopic observations. **C** and **D)** In the absence of As(III), MpACR3-GFP exhibited dual localisation in the ER and PM as demonstrated by using a Δ-s-tether yeast mutant characterized by separation of the ER from the PM in **C)** and by cellular fractionation followed by Western blot analysis of the obtained membrane fractions in **D)**. Plasma membrane H^+^-ATPase (Pma1) was used as a marker of membrane fractions. **E)** The rate of MpACR3-GFP re-localisation from the ER to the PM after the addition of 0.1 mM As(III). **F)** The loss of MpACR3-GFP localisation in the ER was only observed after the addition of arsenic and antimony compounds. Yeast cultures were exposed to 0.1 mM As(III) (sodium arsenite), 0.5 mM As(V) (sodium arsenate), 0.01 mM Cd(II) (cadmium chloride), 0.05 mM Cu(II) (copper sulphate), 1 mM Sb(III) (potassium antimonyl tartrate), 0.5 mM Sb(V) (sodium antimonate), 0.1 mM Zn(II) (zinc sulphate) and 0.25 mM H_2_O_2_ for 2 h. DIC, Differential Interference Contrast. Scale bars = 5 μm.

Since the cortical ER is closely associated with the PM in yeast cells, the ER signal of MpACR3-GFP may overlap with its PM signal. To resolve this, we expressed Mp*ACR3-GFP* in a Δ-s-tether yeast mutant, which lacks six ER-to-PM tethering proteins and thus exhibits a near absence of PM-associated ER (Quon et al. 2018). Interestingly, in the Δ-s-tether strain, MpACR3-GFP remained localized to the ER, but a distinct signal was also observed at the cell surface (Fig. 2C), suggesting that under non-stress conditions, a subset of MpACR3-GFP proteins is also present at the PM. Upon As(III) exposure, MpACR3-GFP completely disappeared from the ER and accumulated exclusively at the PM (Fig. 2C). Western blot analysis of subcellular fractions further confirmed that in the absence of arsenic stress, MpACR3-GFP is present in both ER- and PM-enriched membrane fractions (Fig. 2D). Importantly, despite its ER retention, MpACR3-GFP protein stability remained unaffected, indicating that MpACR3 is not targeted for ER-associated proteasomal degradation (Supplementary Fig. S8). We also examined the rate of MpACR3-GFP re-localisation from the ER to the PM. Upon the addition of 0.1 mM As(III), MpACR3-GFP gradually accumulated at the cell periphery, with the ER-associated GFP signal nearly disappearing within 45-60 min (Fig. 2E).

To determine whether this intracellular redistribution was a specific response to arsenic or a more general reaction to metalloid-induced stress, such as oxidative stress or interactions with cysteine thiol group, we exposed Mp*ACR3-GFP*-expressing cells to hydrogen peroxide and a range of metal(loid)s, including As(V), Cd(II), Cu(II), Sb(III), Sb(V), and Zn(II) (Fig. 2F). Notably, MpACR3-GFP was retained in the ER under all tested conditions except for treatments with arsenic and antimony compounds, which triggered its redistribution to the PM. This strongly suggests that MpACR3 re-localisation from the ER to the PM is a substrate-specific response rather than a general stress reaction.

### MpACR3 enhances arsenic resistance in *A. thaliana*

To investigate the properties of the MpACR3 transporter in plants, we transformed wild-type *A. thaliana*, which lacks an *ACR3* orthologue, with Mp*ACR3-GFP* under the control of the constitutive CaMV 35S promoter. Two independent transgenic lines, *pro35S:*Mp*ACR3-GFP* (T21 and T27), were selected for further analysis. Under normal conditions, wild-type and *pro35S:*Mp*ACR3-GFP* plants exhibited identical phenotypes when grown in soil (Supplementary Fig. S9). Importantly, MpACR3-GFP protein was detected exclusively in extracts from transgenic plants (Fig. 3A). Interestingly, exposure to both As(III) (5 μM) and As(V) (100 μM) for 2 weeks led to increased MpACR3-GFP protein levels compared to control plants (Fig. 3A). However, *MpACR3-GFP* mRNA levels remained unchanged under these conditions (data not shown), suggesting that arsenical exposure enhances the stability of the MpACR3-GFP protein.

**Figure 3.**
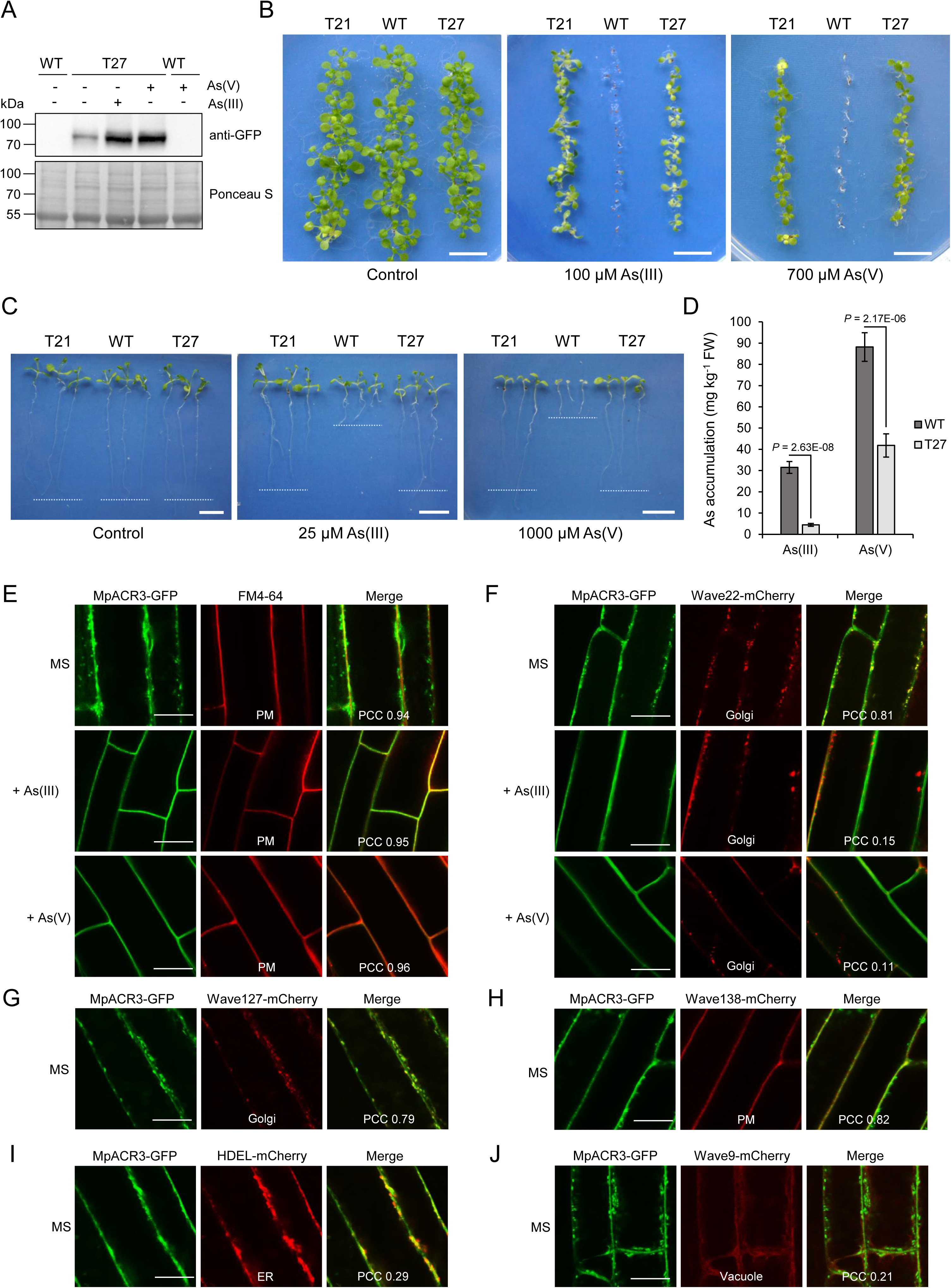
Overexpression of Mp*ACR3-GFP* in *A*. *thaliana*. **A)** Arsenic increases MpACR3-GFP protein levels. Western blot analysis of MpACR3-GFP protein levels in *A. thaliana pro35S:*Mp*ACR3-GFP* transgenic line (T27) grown in the presence of 5 μM As(III) or 100 μM As(V) for 2 weeks or left untreated (MS medium). Protein extracts from wild-type *A. thaliana* (WT, Col-0) were used as a negative control. Ponceau S staining served as a protein loading control. **B** to **D)** Mp*ACR3-GFP* enhances arsenic tolerance in *A*. *thaliana*. **B)** Germination of WT and transgenic *pro35S:*Mp*ACR3-GFP* (T21 and T27 lines) seeds in the absence or presence of As(III) and As(V). Plates were photographed after 2 weeks of growth. Scale bars = 1 cm. **C)** Comparative growth of WT and transgenic (T21 and T27) seedlings grown for 4 d in MS medium followed by 7 d in the absence or presence of As(III) and As(V). Scale bars = 1 cm. **D)** Arsenic accumulation in WT and T27 seedlings grown in the presence of 10 μM As(III) or 200 μM As(V) for 2 weeks. Error bars represent mean value ± standard deviation of mean (*n* = 3). One-way ANOVA was used to calculate the *P*-value. FW, fresh weight. **E** to **J)** Subcellular localisation of MpACR3-GFP in *A. thaliana* root cells. **E** to **H)** MpACR3-GFP co-localized with the plasma membrane (PM) and Golgi bodies under normal conditions (MS medium), whereas in the presence of As(III) or As(V), MpACR3-GFP was exclusively localized to the PM. **I** and **J)** MpACR3-GFP did not co-localized with the ER and vacuole markers. 4-day-old seedlings of transgenic *A*. *thaliana* lines expressing MpACR3-GFP and markers of membrane compartments were transferred to MS medium without or with 25 µM As(III) or 1000 µM As(V) and allowed to grow for an additional 1 day before confocal microscopy analysis. Transgenic *A*. *thaliana* lines are described in the Materials and methods section. Pearson’s correlation coefficient (PCC) was used for quantifying co-localisation of MpACR3-GFP with the indicated markers of membrane compartments. Scale bars = 16 μm.

To assess the impact of *pro35S:*Mp*ACR3-GFP* expression on *A. thaliana* growth under arsenic stress, we examined seed germination, seedling growth, and arsenic accumulation on MS medium supplemented with As(III) or As(V). In the absence of arsenic, no differences were observed between wild-type and *pro35S:*Mp*ACR3-GFP* plants in any of the assays (Fig. 3, B to D). However, in the presence of 100 μM As(III) or 700 μM As(V), *pro35S:*Mp*ACR3-GFP* plants successfully germinated, whereas wild-type germination was inhibited (Fig. 3B). Furthermore, when *A. thaliana* seeds were germinated on standard MS medium for 4 days and then transferred to medium containing 25 μM As(III) or 1000 μM As(V), wild-type plants exhibited more pronounced growth retardation than *pro35S*:Mp*ACR3-GFP* lines (Fig. 3C). Additionally, after 2 weeks of growth on plates supplemented with 10 µM As(III) or 200 μM As(V), concentrations that did not cause severe growth inhibition, transgenic seedlings accumulated significantly lower total arsenic levels compared to wild-type plants (Fig. 3D). These findings indicate that heterologous expression of MpACR3 enhances arsenic tolerance in *A. thaliana*.

### Delayed Golgi-to-PM trafficking of Mp*ACR3* in *A. thaliana* under arsenic-free conditions

The subcellular localisation of MpACR3-GFP in *A. thaliana* was analysed in the rhizodermal cells of roots. The fluorescent signal of MpACR3-GFP was detected at the cell periphery and co-localized with FM4-64, a fluorescent dye commonly used for PM imaging (Fig. 3E). Additionally, MpACR3-GFP was observed in distinct cytoplasmic compartments of varying sizes. Notably, exposure to 25 μM As(III) or 1000 μM As(V) led to the disappearance of intracellular signals, with MpACR3-GFP exclusively co-localizing with the FM4-64-stained PM (Fig. 3E).

To further investigate the intracellular localisation of the MpACR3 transporter, *pro35S:*Mp*ACR3-GFP* transgenic plants were crossed with well-characterized Wave (Geldner et al. 2009) and HDEL (Nelson et al. 2007) mCherry reporter lines. Fluorescence microscopy revealed that MpACR3-GFP co-localized with Golgi markers (Wave line 22R and 127R; Fig. 3, F and G) and the PM marker (Wave line 138R; Fig. 3H). In contrast, it did not co-localize with the ER marker (HDEL-mCherry; Fig. 3I) or the vacuolar marker (Wave line 9; Fig. 3J). Additionally, in protoplasts, MpACR3-GFP co-localized with both the PM (stained with FM4-64) and Golgi bodies (Wave line 22R) (Supplementary Fig. S10).

Upon exposure to 25 μM As(III) or 1000 μM As(V), MpACR3-GFP in Wave22R transgenic root cells no longer co-localized with the Golgi compartment (Fig. 3F). This indicates that under normal conditions, MpACR3-GFP trafficking to the PM is delayed, with a significant portion of the protein retained in the Golgi. However, in the presence of As(III) or As(V), Golgi retention is disrupted, leading to the exclusive accumulation of MpACR3-GFP at the PM. This observation was further confirmed through Lattice Light-Sheet microscopy, which showed that MpACR3-GFP co-localized with the Golgi mCherry marker (Wave127R) only in the absence of arsenic (Supplementary Fig. S11).

To further refine the suborganellar localisation of MpACR3-GFP within Golgi bodies, we transiently co-expressed MpACR3-GFP with markers for different Golgi compartments in tobacco leaf epidermal cells, followed by high-resolution confocal microscopy (McGinness et al. 2022; McGinness et al. 2025; Hawes et al. 2024). MpACR3-GFP was specifically localized to the Golgi and co-localized with the *trans*-Golgi marker ST-mRFP (Fig. 4, A and C). However, it did not overlap with the *cis*-Golgi marker MNS1-mRFP (Fig. 4, B and C) or the *trans*-Golgi network (TGN) marker mRFP-SYP61 (Fig. 4D). These findings suggest that, under normal conditions, MpACR3 is primarily retained in the *trans*-Golgi compartment in plant cells.

**Figure 4.**
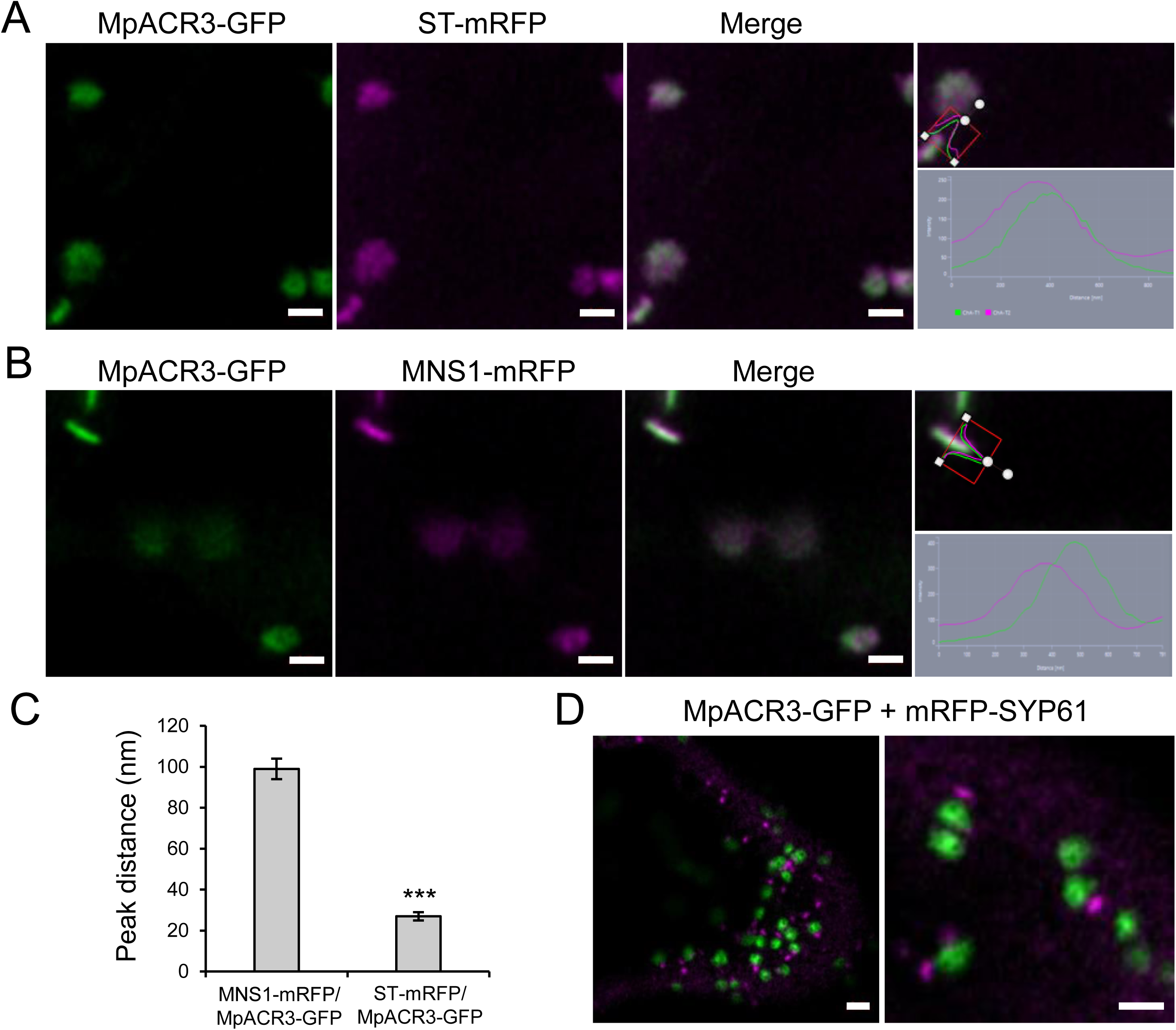
Co-localisation line profile analysis for MpACR3-GFP with *cis-* and *trans*-Golgi marker constructs and TGN marker construct. **A)** Example images and line profile for MpACR3-GFP (green) with the *trans*-Golgi body marker ST-mRFP (magenta) and **B)** with the *cis*-Golgi body marker MNS1-mRFP (magenta). Line profile applied and line profile output are shown. Size bars = 1 µm. **C)** Peak distance analysis graphs showing the markers MpACR3-GFP with MNS1-mRFP and ST-mRFP, respectively. Significance was analysed by Kruskal-Wallis (\*\*\**P* < 0.001). Results shown from *n* = 3 biological replicas with at least 6 technical repeats. For comparison the peak distance difference between MNS1-GFP with MNS1-mRFP is 18 ± 8 nm and MNS1-GFP with ST-mRFP is 122 ± 12 nm (McGinness et al. 2025). **D)** MpACR3-GFP (green) was also co-expressed with the TGN maker mRFP-SYP61 (magenta). Tobacco leaf epidermal cells transiently co-expressing the indicated proteins with fluorescent tags were visualised by Airyscan high-resolution confocal microscopy 3 days after infiltration. Example images are shown. Size bars = 2 µm.

### MpACR3 does not confer Sb(III) tolerance in *A. thaliana*

Unlike in yeast, MpACR3-GFP overexpression did not confer increased Sb(III) resistance in *A. thaliana*, as evidenced by seed germination and seedling growth assays in the presence of Sb(III) (Supplementary Fig. S12, A and B). Additionally, Sb(III) accumulation levels did not significantly differ between *pro35S:*Mp*ACR3-GFP* and wild-type plants (Supplementary Fig. S12C). This aligns with our findings from yeast, where Sb(III) was identified as a poor substrate for MpACR3 (Fig. 1, D and F; Supplementary Figs. S6 and S7). However, similar to arsenic exposure, Sb(III) treatment resulted in an increase in MpACR3-GFP protein levels in *A. thaliana* (Supplementary Fig. S12D). On the other hand, upon Sb(III) exposure, only a subset of MpACR3-GFP relocated to the PM, while intracellular GFP signals remained visible in specific regions of root cells (Supplementary Fig. S12E). This suggests that Sb(III) is also a weak inducer of MpACR3 trafficking in *A. thaliana*.

### Metalloid stress induces upregulation of Mp*ACR3* expression in *M. polymorpha*

To investigate the role of Mp*ACR3* in its native organism, we examined whether its expression is influenced by metalloid exposure in *M. polymorpha* (Cam-1). Using qPCR, we analysed Mp*ACR3* transcript levels in gametophyte tissues under normal conditions and after two weeks of exposure to As(III), As(V), or Sb(III). Mp*ACR3* was constitutively expressed even in the absence of metalloid stress, albeit at low levels (Supplementary Fig. S13). Upon exposure to As(V), transcript levels increased twofold, while Sb(III) induced a slight upregulation. The most pronounced response was observed with As(III), which triggered a threefold increase in Mp*ACR3* expression (Supplementary Fig. S13). These findings suggest that, unlike *ScACR3*, Mp*ACR3* is continuously expressed and moderately upregulated in response to metalloids. The metalloid-dependent regulation of Mp*ACR3* further supports its role in arsenic and antimony detoxification in *M. polymorpha* cells.

### Transcriptomic response of *M. polymorpha* to prolonged As(III) stress

To gain deeper insights into the transcriptomic response to As(III) stress, RNA-seq analysis was conducted on *M. polymorpha* plants grown for one week with or without 2.5 μM As(III). This approach identified 1,283 differentially expressed genes using a false discovery rate (FDR) cutoff of 0.05. Among these, 674 genes were upregulated, while 609 were downregulated (Supplementary Fig. S14; Supplementary Data Set 1). Notably, MpACR3 ranked 23rd among the most significantly regulated genes when ranked by the lowest FDR value, exhibiting a 2.2-fold increase in expression (Supplementary Data Set 1), which aligns with the qPCR results (Supplementary Fig. S14).

Several other genes encoding membrane transporters were significantly upregulated alongside Mp*ACR3*. These included those for a predicted Ca²⁺/H⁺ antiporter (Mp8g17780), a K⁺/H⁺ antiporter (Mp4g10470), and three iron permeases (e.g., Mp4g04160). Genes involved in sulphate and phosphate transport were also enhanced, such as a sulphate permease (Mp1g20200) and seven phosphate transporters (e.g., Mp*PTB4*). Additionally, genes encoding a K⁺/Na⁺ efflux P-type ATPase (Mp3g11080) and four plasma membrane H⁺-ATPases (e.g., Mp*HA18*) were upregulated. Increased expression was also observed in eight genes from the major intrinsic protein (MIP) family, including Mp*TIP2* and Mp*PIP2*.

Arsenic is known to induce oxidative stress in plants (Yadav et al. 2024). In response to As(III), a subset of genes encoding antioxidant enzymes was upregulated, including catalase (Mp*CAT1*), thioredoxin (e.g., Mp3g04870), peroxiredoxin-like (Mp7g05690), and peroxidase (Mp*POD154*, along with 15 other Mp*POD* genes). Conversely, 20 Mp*POD* genes were significantly downregulated. Furthermore, we observed a marked decrease (up to 40-fold) in the expression of 22 genes encoding cytochrome P450 superfamily members (e.g., Mp*CYP829F2*).

As(III) exposure also led to the downregulation of 12 genes for light-harvesting chlorophyll *a*/*b*-binding proteins (e.g., Mp7g05530), along with 10 terpene synthase genes (e.g., Mp*MTPSL15*), and 14 pathogen defence-related genes from the pathogenesis-related (Mp*PR1f*), thaumatin (e.g., Mp2g13870), and Bet v 1 (e.g., Mp8g09000) families. Additionally, genes differentially expressed (both up- and downregulated) under As(III) stress included nine xylanase inhibitor genes, 10 dirigent-like protein genes, 18 cupin genes, 23 transcription factor genes from 11 families, and 48 cell wall-related genes.

In summary, As(III) treatment upregulated genes associated with detoxification, adaptation to As(III)-induced oxidative stress, protein folding stress, and both abiotic and biotic stress responses. Importantly, As(III) toxicity in plants may stem not only from its harmful effects on proteins and other biomolecules but also from the disruption of genes involved in primary and secondary metabolic pathways.

### Overexpression of MpACR3 increases arsenic and antimony tolerance in *M. polymorpha*

To investigate the function and subcellular localisation of MpACR3 protein in *M*. *polymorpha*, we generated a transgenic line expressing the *pro2x35S:*Mp*ACR3-mVenus* transgene in a wild-type background (Cam-1). We first examined MpACR3-mVenus protein levels in plants grown with and without metalloids. Similar to MpACR3-GFP in *A. thaliana*, the level of MpACR3-mVenus protein increased upon exposure to As(III), As(V), and Sb(III) (Fig. 5A). Given this observation, we hypothesized that the CaMV 35S promoter might be upregulated under stress conditions. To test this, we quantified Mp*ACR3-mVenus* mRNA levels using qPCR with primers specific to the *mVenus* region. The results confirmed elevated Mp*ACR3-mVenus* transcript levels in metalloid-treated plants compared to controls (Supplementary Fig. S15), suggesting that metalloid stress enhances CaMV 35S promoter activity, leading to increased MpACR3-mVenus protein expression. Furthermore, *M. polymorpha* overexpressing Mp*ACR3-mVenus* exhibited increased tolerance to As(III), As(V), and Sb(III) (Fig. 5B) while accumulating lower levels of arsenic and antimony than wild-type plants (Fig. 5, C to E). These findings indicate that MpACR3 functions in the efflux of As(III) and Sb(III) in *M. polymorpha*.

**Figure 5.**
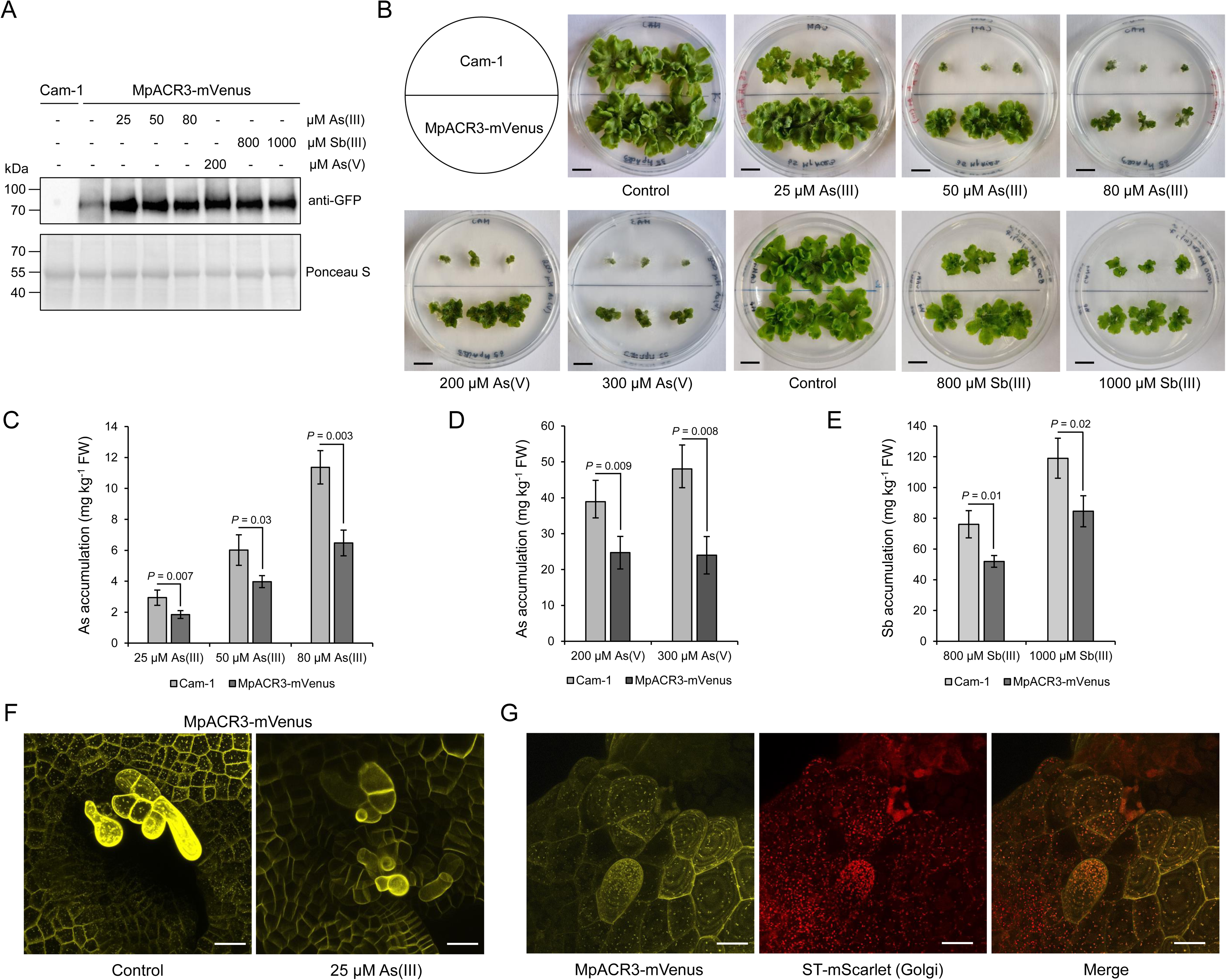
Overexpression of Mp*ACR3-mVenus* in *M*. *polymorpha*. **A)** Western blot analysis of MpACR3-mVenus protein levels in *M*. *polymorpha* transgenic line expressing the *pro35Sx2:*Mp*ACR3-mVenus* transgene grown in the presence of indicated concentrations of As(III), As(V), and Sb(III) for 3 weeks or under normal conditions. Protein extract from wild-type *M*. *polymorpha* (Cam-1) was used as a negative control. Ponceau S staining served as a protein loading control. **B)** Comparative growth of the wild-type Cam-1 and the transgenic MpACR3-mVenus line grown in the absence or presence of the indicated concentrations of As(III), As(V), and Sb(III) for 3 weeks. Scale bars = 1 cm. **C** to **E)** Metalloid accumulation in Cam-1 and MpACR3-mVenus gametophytes grown in the presence of indicated concentrations of metalloids for 3 weeks. Error bars represent mean value ± standard deviation of mean (*n* = 3). One-way ANOVA was used to calculate the *P*-value. FW, fresh weight. **F** and **G)** Subcellular localisation of MpACR3-mVenus in *M*. *polymorpha* gemmae with the use of confocal microscopy. Scale bars = 25 μm. **F)** Yellow fluorescence MpACR3-mVenus protein signal was observed at the cell periphery and in intracellular punctate structures under control conditions, while the exposure to As(III) caused relocation of MpACR3-mVenus from the cytoplasmic compartments to the cell periphery. **G)** MpACR3-mVenus co-localizes intracellularly with the Golgi under control conditions. Co-localisation of MpACR3-mVenus with Golgi bodies was determined in the *M*. *polymorpha* transgenic line expressing the *proUbE2:52aaST-mScarlet* transgene, which encodes a Golgi marker in the form of a transmembrane domain of rat sialyltransferase fused with red fluorescent protein (ST–mScarlet).

### Arsenic triggers the relocation of MpACR3 from the Golgi to the PM in *M. polymorpha*

Here, we demonstrate that under normal conditions, MpACR3-GFP localizes to the ER in *S. cerevisiae* (Fig. 2) and to the Golgi in *A. thaliana* (Figs. 3 and 4). However, upon As(III) stress, MpACR3-GFP accumulates at the PM in both yeast and *A. thaliana* cells. To further investigate its subcellular localisation in *M. polymorpha*, we used confocal microscopy to examine MpACR3-mVenus in gemmae with or without As(III) treatment. Under normal conditions, MpACR3-mVenus fluorescence was detected both at the cell periphery and in intracellular punctate structures. Following As(III) exposure, the intracellular signal was lost, and MpACR3-mVenus was exclusively localized at the cell periphery, consistent with PM accumulation (Fig. 5F).

Since MpACR3 localizes to the Golgi in *A. thaliana* (Figs. 3 and 4), we hypothesized that the same occurs in *M. polymorpha*. To test this, we generated a transgenic line co-expressing the *proUbE2:52aaST-mScarlet* Golgi marker alongside *pro35Sx2:*Mp*ACR3-mVenus*. Under normal conditions, MpACR3-mVenus co-localized with ST-mScarlet-labeled Golgi bodies (Fig. 5G), confirming its Golgi localisation. In conclusion, our findings suggest that in plant cells, MpACR3 localizes to both the Golgi and PM under normal conditions. However, under arsenic stress, it bypasses Golgi retention and accumulates exclusively at the PM, likely facilitating metalloid detoxification.

### Regulation of MpACR3 trafficking by its N-terminal cysteine-rich domain

The predicted structure of MpACR3 features a large cytosolic N-terminal domain comprising 123 amino acid residues (Supplementary Fig. S3), including five cysteine residues (Fig. 6A). Importantly, C29, C47, and C71 are highly conserved among plant ACR3 transporters with extended N-terminal tails (104-163 residues), such as the moss *Physcomitrium patens* ACR3 (PpACR3) (Fig. 6A; Supplementary Fig. S4). In contrast, ACR3 sequences from bacteria, fungi, and certain plants, primarily chlorophytes and ferns, exhibit much shorter (less than 100 residues) N-terminal tails, which typically lack cysteine residues (Fig. 6A; Supplementary Figs. S2 and S5). Proteins involved in arsenic detoxification are often regulated by As(III) binding to vicinal cysteine residues (Shi et al. 1996; Kumar et al. 2015; Lee and Lewin, 2022). Given this, we hypothesized that the N-terminal cysteine-rich domain may modulate MpACR3 trafficking by detecting cytoplasmic As(III).

**Figure 6.**
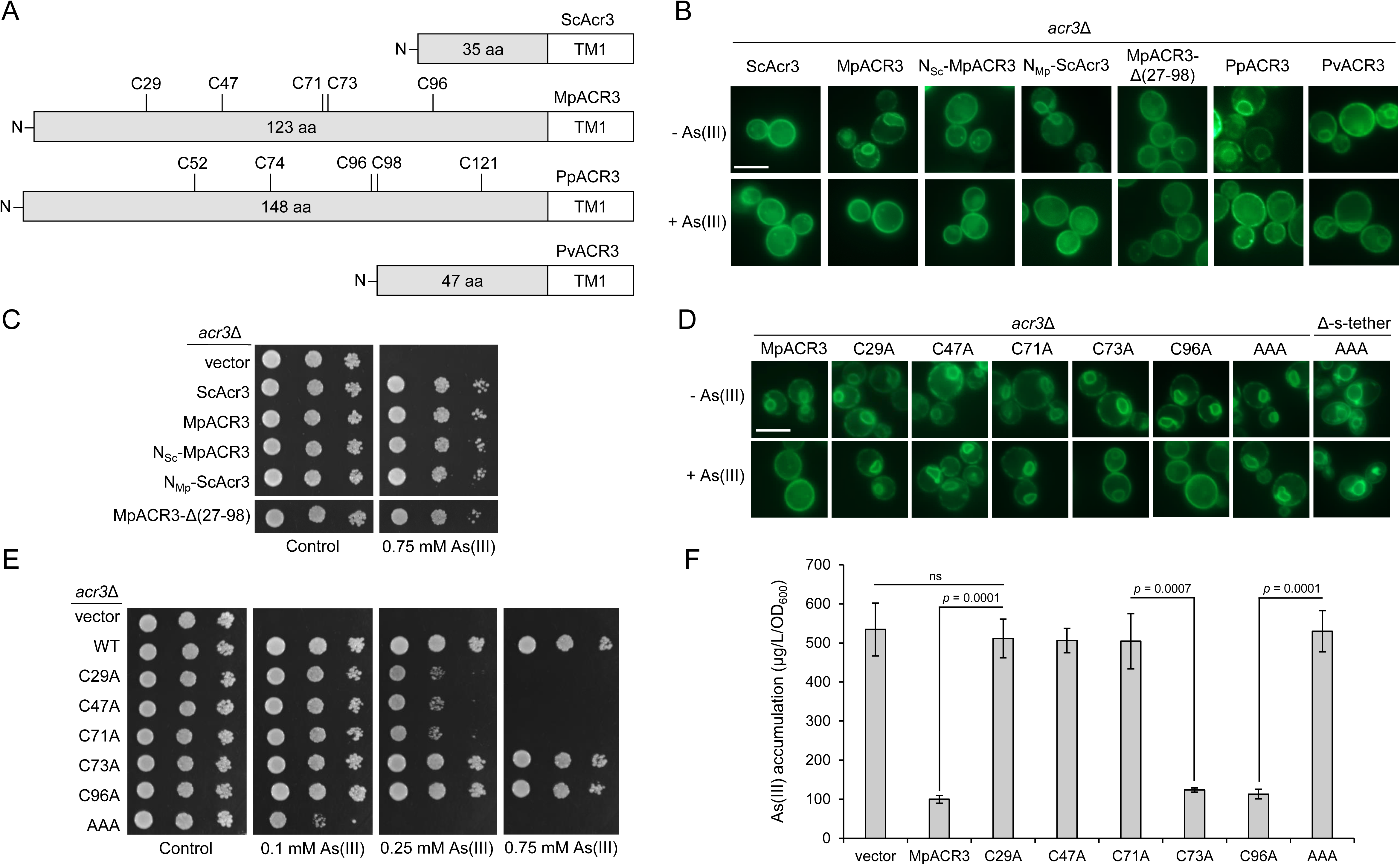
The N-terminal cysteine-rich domain regulates As(III)-induced relocation of ACR3 proteins from the ER to the PM in yeast cells. **A)** Schematic showing the length of the N-terminal cytosolic tails of the yeast *S*. *cerevisiae* (ScACR3), the liverwort *M*. *polymorpha* (MpACR3), the moss *Physcomitrium patens* (PpACR3), and the fern *Pteris vittata* (PvACR3) ACR3 proteins. The positions of cysteine residues and the first transmembrane span (TM1) are indicated. **B)** The N-terminal cysteine-rich region determines ER retention of MpACR3 and PpACR3 proteins in the absence of As(III) in yeast cells. The *acr3*Δ strain was transformed with plasmids expressing GFP-tagged ScACR3, MpACR3, PpACR3, and PvACR3 proteins, N-terminal tail-swapped variants of MpACR3 (N_Sc_-MpACR3) and ScACR3 (N_Mp_-ScACR3), and the MpACR3-Δ27-98 variant lacking the cysteine-rich fragment in N-terminal tail. The localisation of ACR3-GFP proteins was monitored under normal conditions or after exposure to 0.1 mM As(III) for 2 h by fluorescence microscopy. Scale bars = 5 μm. **C)** Tail-swap variants and the MpACR3-Δ27-98 variant showed wild-type resistance to As(III). **D)** C29, C47, and C71 residues are required for As(III)-induced relocation of MpACR3 from the ER to the PM. The subcellular localisation of the indicated variants of MpACR3, including single cysteine to alanine variants and a triple alanine variant (C29A C47A C71A or AAA) was determined in *acr3*Δ cells, as well as in the Δ-s-tether strain for the AAA variant, by fluorescence microscopy. Yeast transformants were exposed to 0.1 mM As(III) for 2 h or left untreated. Scale bars = 5 μm. **E)** Transformants expressing cysteine to alanine MpACR3 variants, defective in As(III)-induced relocation from the ER to the PM, showed decreased tolerance to As(III) compared to wild-type MpACR3 (WT). **F)** Relocation-defective cysteine to alanine variants accumulated high levels of As(III). Accumulation of As(III) in the indicated *acr3*Δ transformants was measured after exposure to 1 mM As(III) for 2 h using an atomic absorption spectrometer. Error bars represent mean value ± standard deviation of mean (*n* = 3). One-way ANOVA was used to calculate the *P*-value. ns, not significant.

To investigate the role of the N-terminal cysteine-rich domain in MpACR3 localisation, we engineered plasmids encoding N-terminally swapped variants: MpACR3-GFP with the *S. cerevisiae* ACR3 N-terminal region (N_Sc_-MpACR3-GFP) and ScACR3-GFP with the *M. polymorpha* ACR3 N-terminal region (N_Mp_-ScACR3-GFP). Additionally, we generated a truncated MpACR3 variant lacking the cysteine-rich region (MpACR3-Δ27-98-GFP). These constructs were heterologously expressed in yeast *acr3*Δ cells. In the absence of As(III), both N_Sc_-MpACR3-GFP and MpACR3-Δ27-98-GFP localized exclusively to the PM, similar to ScACR3, and were no longer retained in the ER. Conversely, N_Mp_-ScACR3-GFP was retained in the ER under normal conditions but relocated to the PM upon As(III) exposure (Fig. 6B). Similarly, PpACR3-GFP, which also contains an N-terminal cysteine-rich domain (Fig. 6A), followed the same localisation pattern as MpACR3-GFP and N_Mp_-ScACR3-GFP (Fig. 6B). In contrast, PvACR3-GFP from the fern *P. vittata*, which possesses a short N-terminal tail, predominantly localized to the PM, with a fraction also present in the ER, regardless of As(III) exposure (Fig. 6B). Despite these differences in localisation, all tested variants (N_Sc_-MpACR3-GFP, N_Mp_-ScACR3-GFP, and MpACR3-Δ27-98-GFP) conferred wild-type levels of As(III) resistance, confirming their retained functionality (Fig. 6C). Collectively, these findings demonstrate that the N-terminal cysteine-rich domain governs the subcellular trafficking of plant ACR3 proteins, ensuring their proper sorting to the PM.

To determine which cysteine residues in the N-terminal domain regulate MpACR3 trafficking, potentially through direct As(III) binding, we individually substituted each cysteine with alanine, generating the variants C29A, C47A, C71A, C73A, and C96A. Under normal conditions, all MpACR3 variants localized to the ER and PM, similar to wild-type MpACR3 (Fig. 6D). However, upon As(III) exposure, the C29A, C47A, and C71A variants failed to re-localize from the ER to the PM (Fig. 6D). Additionally, these variants significantly reduced As(III) resistance compared to wild-type MpACR3 (Fig. 6E) and did not decrease As(III) accumulation in yeast cells (Fig. 6F). The C73A variant exhibited a mild defect in As(III)-induced re-localisation (Fig. 6D) but maintained wild-type As(III) tolerance (Fig. 6, E and F). Meanwhile, the C96A variant displayed no abnormalities in any assay.

To further investigate the role of multiple cysteine residues, we generated a triple-cysteine-to-alanine mutant (C29A C47A C71A, referred to as AAA) and expressed it in *acr3*Δ cells as well as in the Δ-s-tether strain, which exhibits ER-PM separation. Similar to the single-cysteine variants, the AAA mutant primarily localized to the ER, with only a fraction reaching the PM, regardless of As(III) presence (Fig. 6D). Notably, *acr3*Δ cells expressing the AAA variant were more sensitive to As(III) than those expressing single-cysteine mutants (Fig. 6E), suggesting that disrupting the cysteine-rich N-terminal structure may further impair MpACR3’s transport function. Overall, these findings strongly indicate that C29, C47, and C71 are essential for As(III)-induced MpACR3 trafficking to the PM.

### The N-terminal domain of MpACR3 functions as an intracellular arsenic sensor

To determine whether As(III) directly binds to the N-terminal domain of MpACR3, we expressed a 1–123 amino acid N-terminal fragment fused to a 7×HA tag (7HA-MpACR3-N-tail) in *acr3*Δ cells, along with the AAA (C29A C47A C71A) and Cys-null (C29A C47A C71A C73A C96A) variants. Protein extracts from these cells were treated with As(III) for 60 min or left untreated, followed by incubation with an arsenite-biotin conjugate probe. Samples were then subjected to pulldown using streptavidin-agarose resins and analysed via Western blot with anti-HA antibodies.

Our results showed that the As-biotin conjugate bound specifically to the N-terminal domain of wild-type MpACR3, but not to the AAA or Cys-null variants (Fig. 7A). Furthermore, pretreatment of protein extracts with As(III) abolished the binding of As-biotin to the MpACR3 N-terminus (Fig. 7A). These findings strongly suggest that As(III) directly interacts with the N-terminal domain of MpACR3 through C29, C47, and C71, highlighting its role as an intracellular arsenic sensor.

**Figure 7.**
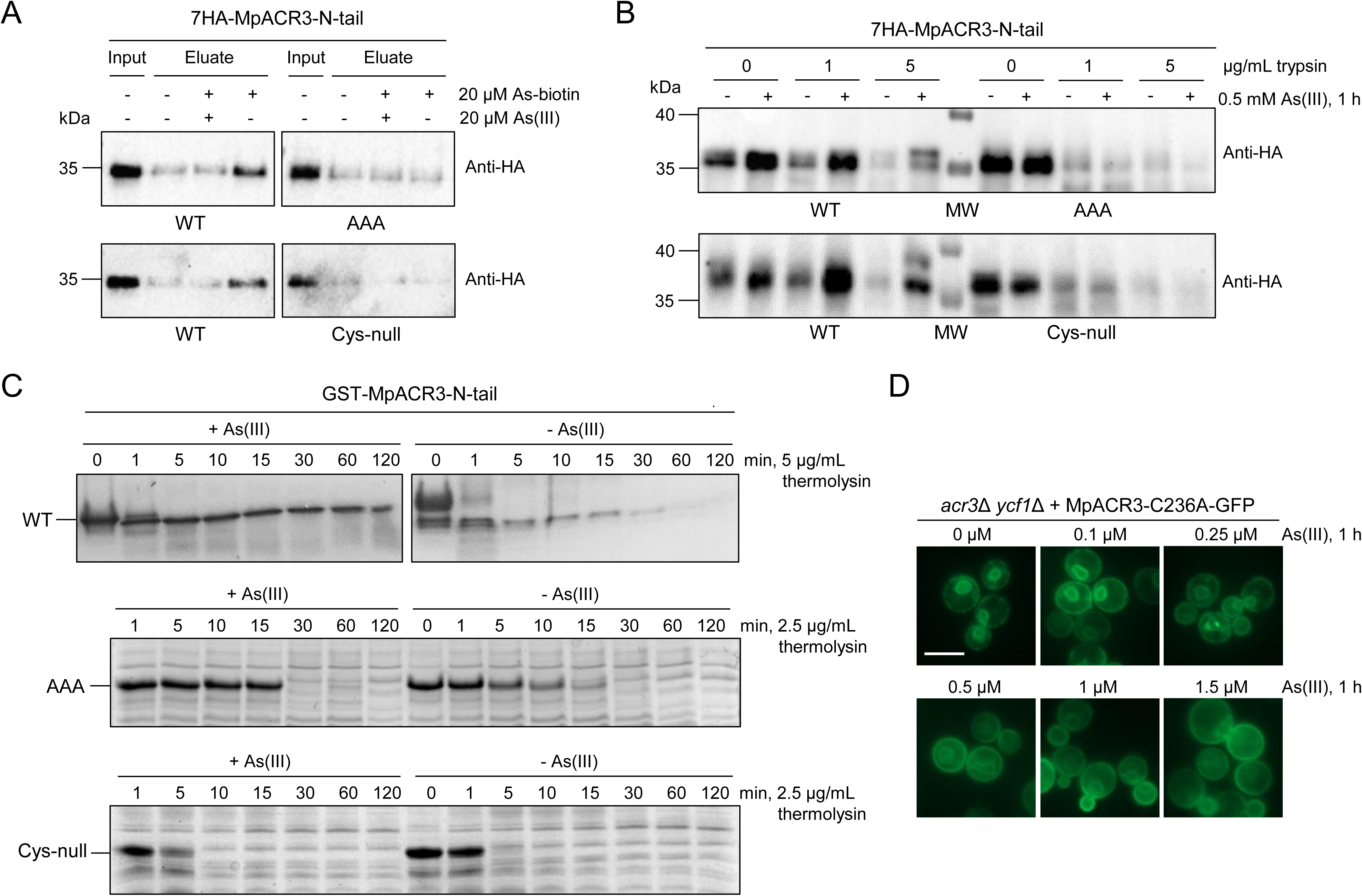
The N-terminal cysteine-rich domain of MpACR3 functions as an arsenic sensing domain. A) As-biotin conjugate binds to the N-terminal cysteine-rich domain of MpACR3. Lysates of *acr3*Δ cells expressing the HA-tagged N-terminal domain of MpACR3 (WT), the AAA (C29A C47A C71A) protein variant, or the Cys-null (C29A C47A C71A C73A C96A) protein variant were pre-incubated or not with 20 μM As(III) and then incubated with 20 μM As-biotin conjugate, followed by pulldown with streptavidin-agarose resins and Western blot analysis with anti-HA antibodies. **B** and **C)** As(III) binding to the N-terminal cysteine-rich domain of MpACR3 induces its conformational change both in vivo and in vitro. **B)** Lysates were prepared from *acr3*Δ cells expressing the indicated variants of the N-terminal domain of MpACR3 that were treated or not with 0.5 mM As(III) for 60 min. The lysates were then exposed to increasing concentrations of trypsin for proteolysis, followed by Western blot analysis with anti-HA antibodies. MW, molecular weight marker. **C)** GST-fused protein variants of the MpACR3 N-terminal domain were purified from *Escherichia coli* lysates by affinity purification using glutathione-Sepharose columns. Purified GST fusion proteins were treated or not with 20 μM As(III) and then subjected to time course of limited thermolysin digestion. Proteolysis patterns were visualized using SDS-PAGE electrophoresis. **D)** Arsenic-hypersensitive yeast cells expressing the transport-inactive MpACR3-C236A-GFP protein can be used as a biosensor to detect environmentally relevant As(III) concentrations. The *acr3*Δ *ycf1*Δ strain was transformed with a plasmid expressing the MpACR3-C236A-GFP variant protein and then exposed to indicated concentrations of As(III). The subcellular localisation of MpACR3-C236A-GFP was visualized by fluorescence microscopy.

### As(III) binding induces a conformational change in the MpACR3 N-terminal domain

Ligand binding can significantly influence the susceptibility of a protein segment to limited proteolysis. For example, As(III) binding induces a conformational change in the yeast transcription factor ScYap8, reducing its susceptibility to trypsin digestion and enabling activation of *ScACR3* transcription (Kumar et al. 2015). Based on this, we hypothesized that As(III) binding to the MpACR3 N-terminal domain induces a structural change that influences its intracellular retention.

To test this, we prepared protein extracts from yeast cells expressing wild-type 7HA-MpACR3-N-tail or its cysteine-deficient variants. These extracts were either treated with As(III) or left untreated before incubation with increasing concentrations of trypsin. Proteolysis patterns were analysed via anti-HA immunoblotting (Fig. 7B; Supplementary Fig. S16). Our results showed that As(III) exposure increased the resistance of wild-type 7HA-MpACR3-N-tail to trypsin digestion compared to untreated samples. In contrast, both the AAA and Cys-null variants remained highly susceptible to proteolysis, regardless of As(III) treatment. Notably, even in the absence of As(III), wild-type 7HA-MpACR3-N-tail exhibited slightly greater resistance to trypsin digestion than its cysteine-deficient counterparts. These findings suggest that As(III) binding stabilizes the MpACR3 N-terminal domain, triggering a conformational change that may regulate its intracellular localisation.

To further investigate the structural effects of As(III) binding, we performed a protease resistance-based protein stability assay using recombinantly expressed and purified GST-MpACR3-N-tail and its cysteine-deficient variants. These proteins were incubated with thermolysin, a nonspecific endoproteinase, in the presence or absence of As(III). In the ligand-free state, GST-MpACR3-N-tail was highly susceptible to thermolysin digestion. However, when bound to As(III), it became remarkably resistant to proteolysis, even over extended incubation times (Fig. 7C). Since substrate binding in the protease active site requires an extended conformation (Tyndall et al. 2005), these findings suggest that As(III) binding induces a stable, highly ordered fold in the MpACR3 N-terminal domain. As expected, the two cysteine-deficient variants, GST-MpACR3-AAA-N-tail and GST-MpACR3-Cys-null-N-tail, were more readily degraded than wild-type GST-MpACR3-N-tail. However, in the presence of As(III), GST-MpACR3-AAA-N-tail displayed slightly increased resistance to thermolysin digestion compared to its ligand-free state, suggesting that, at least in vitro, As(III) may also interact with C73 and C96. Despite using a reduced protease concentration, the Cys-null variant remained highly susceptible to rapid and complete degradation, regardless of As(III) presence (Fig. 7C). This accelerated proteolysis suggests that cysteine residues in the MpACR3 N-terminal domain not only mediate As(III) binding but may also contribute to structural stability.

To explore the potential of MpACR3 as a biological sensor for detecting As(III) in the environment, we expressed a transport-inactive variant, MpACR3-C236A-GFP, in the *acr3*Δ *ycf1*Δ double mutant, which lacks both PM and vacuolar As(III) detoxification pathways. This ensured maximal cytoplasmic As(III) accumulation. Cells were then exposed to varying concentrations of As(III) for 1 h. Notably, MpACR3-C236A-GFP re-localized from the ER to the PM at As(III) concentrations as low as 0.5-1 µM (Fig. 7D). These findings highlight the sensitivity of the MpACR3 N-terminal domain to As(III) and suggest its potential application in developing a cost-effective, biologically based technology for environmental As(III) detection.

Taken together, these results support the role of the MpACR3 N-terminal domain as an intracellular arsenic sensor, with As(III) binding to C29, C47, and C71 triggering a conformational change that likely influences MpACR3 intracellular retention.

### Molecular modelling and structural analysis of the MpACR3 arsenic sensing domain

To further explore the mechanistic aspects and the nature of a potential conformational transition induced by As(III) binding to C29, C47, and C71, we performed molecular modelling. As a starting structure, we employed the MpACR3 AlphaFold model structure (Jumper et al. 2021), from which we extracted the 24-105 amino acid residue region. We first processed the model structure with Maestro BioLuminate program (Salam et al. 2014), which resulted in two disulphide bridges between the side group (SG) atoms of C47-C71 and C29-C96. However, we deemed these disulphide bridges unlikely due to the cytosolic localisation of the domain. As a result, we modified the structure and added hydrogen atoms to the corresponding SG atoms, creating the “WT” (wild-type) model. Additionally, we generated a triple cysteine-to-alanine mutant (C29A, C47A, C71A), referred to as “Triple Ala.” Both the WT and Triple Ala structures were subjected to separate two microseconds-long all-atom unrestrained molecular dynamics (MD) simulations. Also, we conducted a restrained two microseconds-long all-atom MD simulation. In the restrained simulation, the distances between the SG atoms of the C29-C47, C47-C71, and C29-C71 pairs were maintained within a range of 3.5 to 3.8 Å, corresponding to SG-SG distances when As(III) is bound (for further details, see Materials and methods). We analysed the three MD trajectories in terms of the average structures (Fig. 8A; Supplementary Fig. S17): the root mean square deviation (RMSD) of the protein heavy atoms (Supplementary Fig. S18); the root mean square fluctuation (RMSF) for each residue (Supplementary Fig. S19); and the evolution of the SG-SG distances (Supplementary Fig. S20). We also compared the average structures topology (Supplementary Fig. S21) and molecular surface properties, including electrostatic surface potential (Fig. 8B) and hydrophobicity (Fig. 8C).

**Figure 8.**
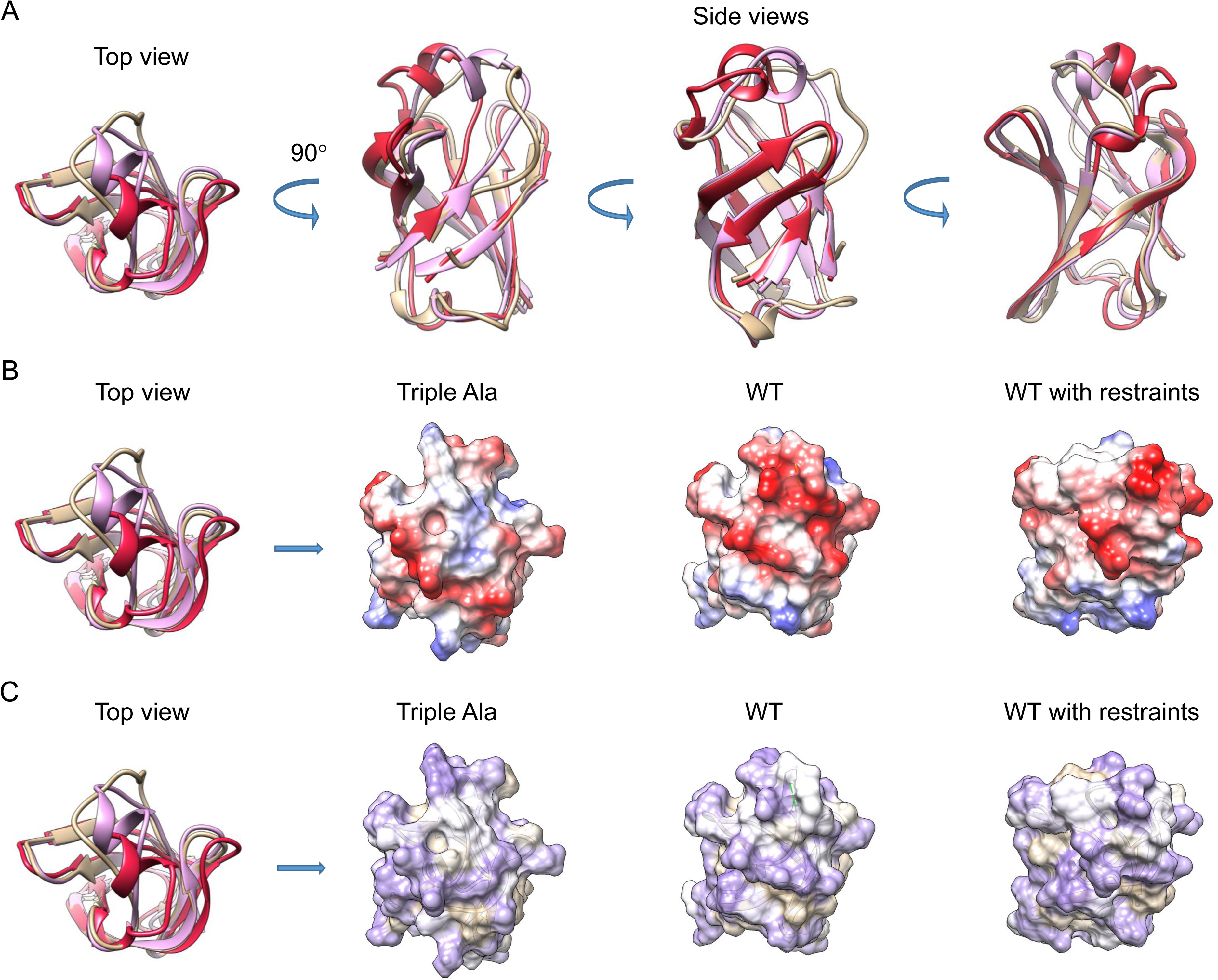
Structure modelling of the MpACR3 arsenic-sensing domain. **A)** Comparison of MD trajectory average structures of the N-terminal domain: Triple Ala (C29A, C47A, C71A) variant (tan), wild-type (WT; pink), and WT with Cys-Cys restraints (dark red). The main difference is the formation of a helical structure in the 30–45 residue region. **B)** Electrostatic surface potentials viewed from the top, highlighting structural differences in the same region for the Triple Ala variant, WT, and WT with restraints. Colour coding: red – negative, white – neutral, blue – positive. **C)** Molecular surface comparison based on amino acid hydrophobicity, shown from the top for all three models. Colour coding: purple – most hydrophilic, tan – most hydrophobic.

Comparing the evolution of the three model structures during the MD trajectories, we observed the formation of an α-helical region around 35-40 amino acid residues in the WT model. This conformational behaviour can be explained by two hydrogen bonds present in the WT structure: the C71 HG – F69 O bond, which in turn enables the formation of the C29 HG – N65 OD1 bond (Supplementary Fig. S22A). When the SG-SG restraints were switched on, the α-helical region expanded further to 32-45 residues. In contrast, in the Triple Ala model the 32-45 region remained in a random coil conformation throughout the entire trajectory (Fig. 8; Supplementary Figs. S19, S21, and S23).

The switching on of the SG-SG restraints made the model structure not only more compact but also influenced its surface properties (Fig. 8). The α-helical conformation of the 32-45 region created a negatively charged patch of the molecular surface, potentially modulating protein-protein interactions. Furthermore, when comparing the RMSF plots for the WT and WT with restraints models, we observed increased conformational flexibility in the 85-95 region. This flexibility arose from the disappearance of two hydrogen bonds when the C29, C47, and C71 residues were in positions suitable for coordinating an As(III) atom (Supplementary Fig. S22B), namely: K28 H-E97 O and R46 H-L87 O. We hypothesized that this increased flexibility may be advantageous for protein-protein interactions.

### Identification of an arginine-based sorting signal in the MpACR3 N-terminal domain

To elucidate the molecular mechanisms governing MpACR3 intracellular localisation, we analysed its N-terminal domain for known sequence motifs critical for ER or Golgi retention (Michelsen et al. 2005; Gao et al. 2014; Luján and Campelo 2021). This analysis revealed an arginine-based sorting signal of the RXR type (where X represents any amino acid) within the highly conserved 45-LRCRF-47 sequence, with the arsenic-sensing residue C47 positioned centrally (Fig. 9A; Supplementary Fig. S4). Arginine-based sorting signals play a key role in the retention of ER and Golgi membrane proteins and are widely conserved among eukaryotes (Michelsen et al. 2005; Banfield, 2011; Luján and Campelo 2021). Additionally, functional sorting signals often contain a bulky hydrophobic residue, typically leucine, preceding an arginine cluster (RR, RXR, or RRXR). These motifs interact with the coat protein complex I (COPI), facilitating retrograde transport within the Golgi and from the Golgi to the ER (Zerangue et al. 2001; Michelsen et al. 2005; Beck et al. 2009).

**Figure 9.**
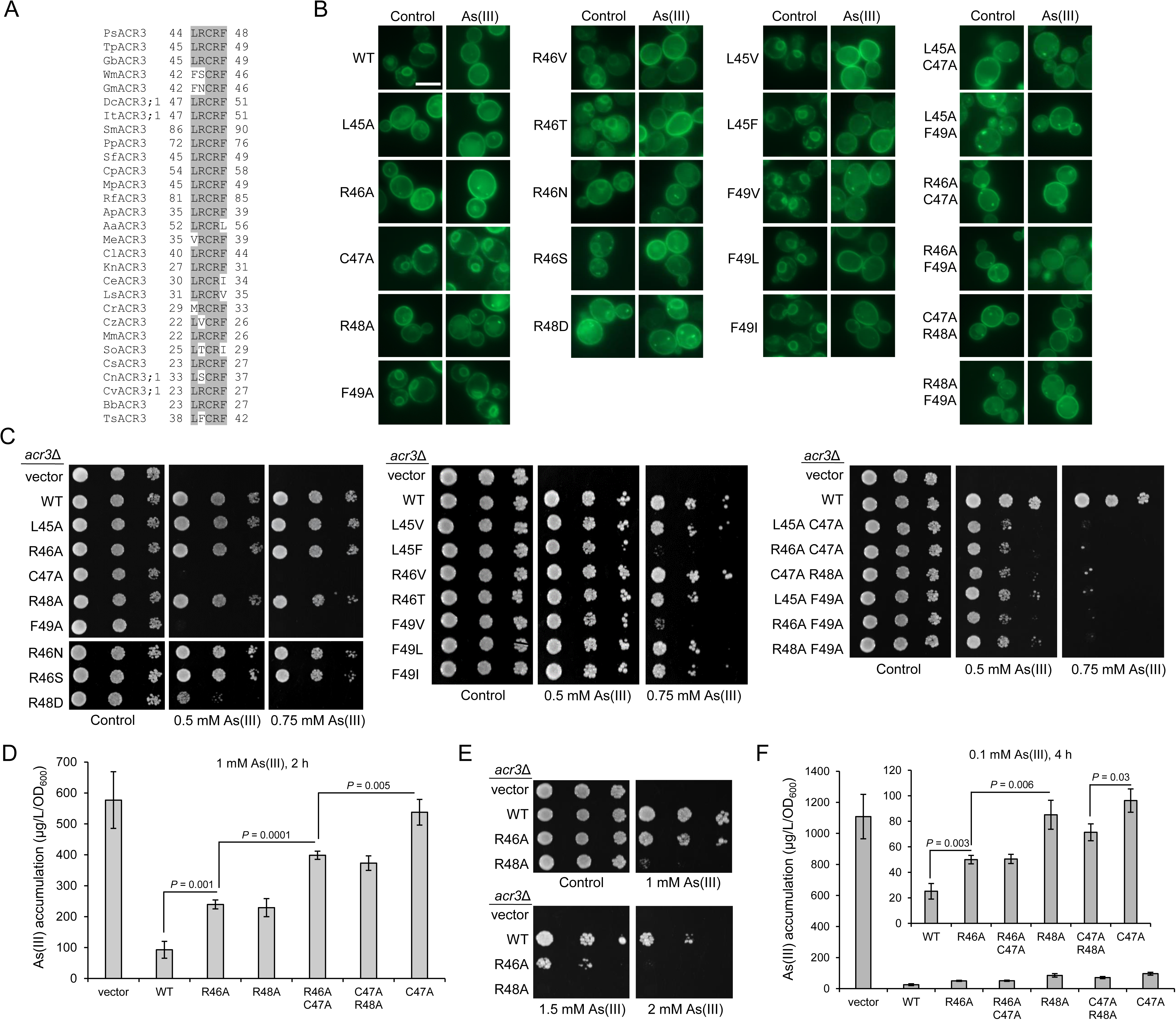
Functional analysis of the LRCRF motif in MpACR3 intracellular retention and function. **A)** Multiple sequence alignment of LRCRF regions of plant ACR3 proteins. Sequence details are provided in Supplementary Table S1. **B)** The localisation of MpACR3-GFP variants with a mutated LRCF motif in *acr3*Δ cells was monitored under normal conditions or after exposure to 0.1 mM As(III) for 2 h by fluorescence microscopy. Scale bar = 5 μm. **C)** As(III) sensitivity of *acr3*Δ transformants expressing the LRCRF motif variants. **D)** As(III) accumulation in *acr3*Δ cells expressing the LRCRF motif variants. Accumulation of As(III) was measured after exposure to 1 mM As(III) for 2 h using an atomic absorption spectrometer. **E)** The growth of the *acr3*Δ strain expressing MpACR3-GFP or its R46A and R48A variants in the presence of high concentrations of As(III). **F)** As(III) accumulation in *acr3*Δ cells expressing the LRCRF motif variants under exposure to low concentrations of As(III). Error bars represent mean value ± standard deviation of mean (*n* = 3). One-way ANOVA was used to calculate the *P*-value.

To determine the role of the LRCRF motif in MpACR3 intracellular retention, we performed alanine-scanning mutagenesis and analysed the subcellular localisation of the resulting variants in yeast cells. Microscopic observations revealed that the L45A, R46A, and R48A variants failed to localize to the ER in the absence of As(III), confirming that these residues are essential for MpACR3 retention under normal conditions (Fig. 9B). In contrast, similar to C47A, the F49A variant remained ER-localized regardless of As(III) presence. This suggests that F49 may play a structural role in either facilitating As(III) binding to the sensor or stabilizing its conformational change upon As(III) interaction (Fig. 9A). Consequently, MpACR3 variants defective in ER retention retained their ability to confer As(III) tolerance. However, expression of the F49A variant failed to complement the *acr3*Δ mutant’s sensitivity to As(III), indicating its critical role in MpACR3 function (Fig. 9C).

Sequence alignment of plant ACR3 proteins revealed variations in the LRCRF region (Fig. 9A; Supplementary Fig. S4). To identify the consensus sequence responsible for regulating plant ACR3 sorting, we generated a series of MpACR3 variants: L45F, L45V, R46N, R46S, R46T, R46V, F49I, F49L, and F49V. Fluorescence microscopy showed that all variants, except R46N, R46S, and R46T, retained strong GFP signals in the ER under normal conditions, which disappeared upon As(III) exposure, mirroring wild-type MpACR3-GFP behaviour (Fig. 9B). In contrast, R46N, R46S, and R46T mutants exhibited impaired ER retention, as evidenced by increased PM localisation, the presence of punctate cytoplasmic structures, and GFP accumulation inside the vacuole, suggesting endocytic degradation of MpACR3-GFP (Fig. 9B). These findings indicate that replacing R46 with asparagine, serine, or threonine, but not valine, substantially weakens the ER retention signal, at least under conditions of Mp*ACR3* overexpression in yeast. Importantly, despite these localisation differences, all tested variants remained functional, as confirmed by their ability to complement the *acr3*Δ mutant’s As(III) sensitivity (Fig. 9C).

Given the high conservation of the second arginine residue in the LRCRF motif (Fig. 9A; Supplementary Fig. S4), we generated the R48D variant to assess the significance of its positive charge. As expected, this variant no longer predominantly localized to the ER under normal conditions. However, in both the absence and presence of As(III), its GFP signal was detected not only at the PM but also in the ER and punctate cytoplasmic structures, likely representing Golgi bodies or endosomes (Fig. 9B). Functionally, the R48D variant exhibited significantly reduced As(III) tolerance compared to wild-type MpACR3 or the R48A variant, suggesting that arginine residues in the LRCRF motif may contribute not only to ER retention but also to MpACR3 function at the PM.

To determine whether substitutions at L45, R46, or R48 could rescue the PM accumulation defects of the C47A and F49A variants and restore As(III) resistance, we generated double variants. As expected, all tested double mutants, including R46A C47A and R46A F49A, predominantly localized to the PM, regardless of As(III) presence (Fig. 9B). Despite their presence at the PM, these double variants exhibited only partial restoration of As(III) tolerance, failing to reach wild-type levels (Fig. 9C). The impaired function of R46A C47A and C47A R48A was further confirmed by As(III) accumulation assays, where yeast expressing these variants showed reduced arsenic transport activity at 1 mM As(III) (Fig. 9D). Interestingly, yeast cells expressing the R46A or R48A single variants accumulated more As(III) than the wild-type MpACR3 transformant (Fig. 9D). Moreover, these variants failed to confer resistance at high As(III) concentrations (Fig. 9E). However, consistent with their ability to promote yeast growth at low As(III) concentrations (Fig. 6E; Fig. 9, C and E), expression of R46A, C47A, R48A, and their double variants prevented excessive arsenic accumulation in yeast exposed to 0.1 mM As(III) (Fig. 9F). Despite this, noticeable differences in transport activity remained among the variants and wild-type MpACR3, suggesting that the conformation of the N-terminal arsenic sensor and surface charge distribution influence MpACR3 function at the PM.

In summary, we propose that the consensus motif for intracellular retention of plant ACR3 proteins is Φ-R/V-X1-R-X2, where Φ represents a bulky hydrophobic residue. However, for this signal to be inactivated upon As(III) exposure, a cysteine at X1 and a bulky hydrophobic residue at X2 (Φ-R/V-C-R-Φ) are required. Additionally, our findings suggest that the N-terminal arsenic sensor, which includes the LRCRF motif, may also play a role in regulating the transport function of PM-localized MpACR3. Further investigations on other members of the plant ACR3 family are necessary to conclusively define this motif.

### Role of the LRCRF motif and its functional implications in plant cells

To determine whether MpACR3 sorting and function follow the same mechanism in plant cells, we transiently expressed wild-type MpACR3-mVenus and its R46A, C47A, and R46A C47A variants in tobacco leaf epidermal cells. Under normal conditions, wild-type MpACR3 and the C47A variant localized to the Golgi and PM, whereas R46A and R46A C47A variants were exclusively found at the PM (Fig. 10A). Upon As(III) exposure, wild-type MpACR3 but not the C47A variant re-localized from the Golgi to the PM (Fig. 10A). Next, we generated two *M. polymorpha* transgenic lines overexpressing C47A and R46A C47A variants of MpACR3-mVenus. Microscopic analysis showed that, unlike wild-type MpACR3, the R46A C47A variant exhibited only PM localisation in *M. polymorpha*, while the C47A variant failed to re-localize from the Golgi to the PM after As(III) treatment (Fig. 10B). Importantly, expression of both variants did not enhance *M. polymorpha* growth in the presence of As(III) or As(V) (Fig. 10C). Furthermore, arsenic content analysis showed that both MpACR3 variant-expressing lines accumulated more arsenic than the wild-type MpACR3-overexpressing line (Fig. 10, D and E). Notably, similar to yeast cells, *M. polymorpha* cells expressing the R46A C47A variant accumulated less arsenic than those expressing C47A alone, but only under As(V) exposure (Fig. 10E). Based on these results, we conclude that in plants, the LRCRF region plays a key role in MpACR3 intracellular retention and As(III) sensing, and likely impacts the transport function of PM-localized MpACR3.

**Figure 10.**
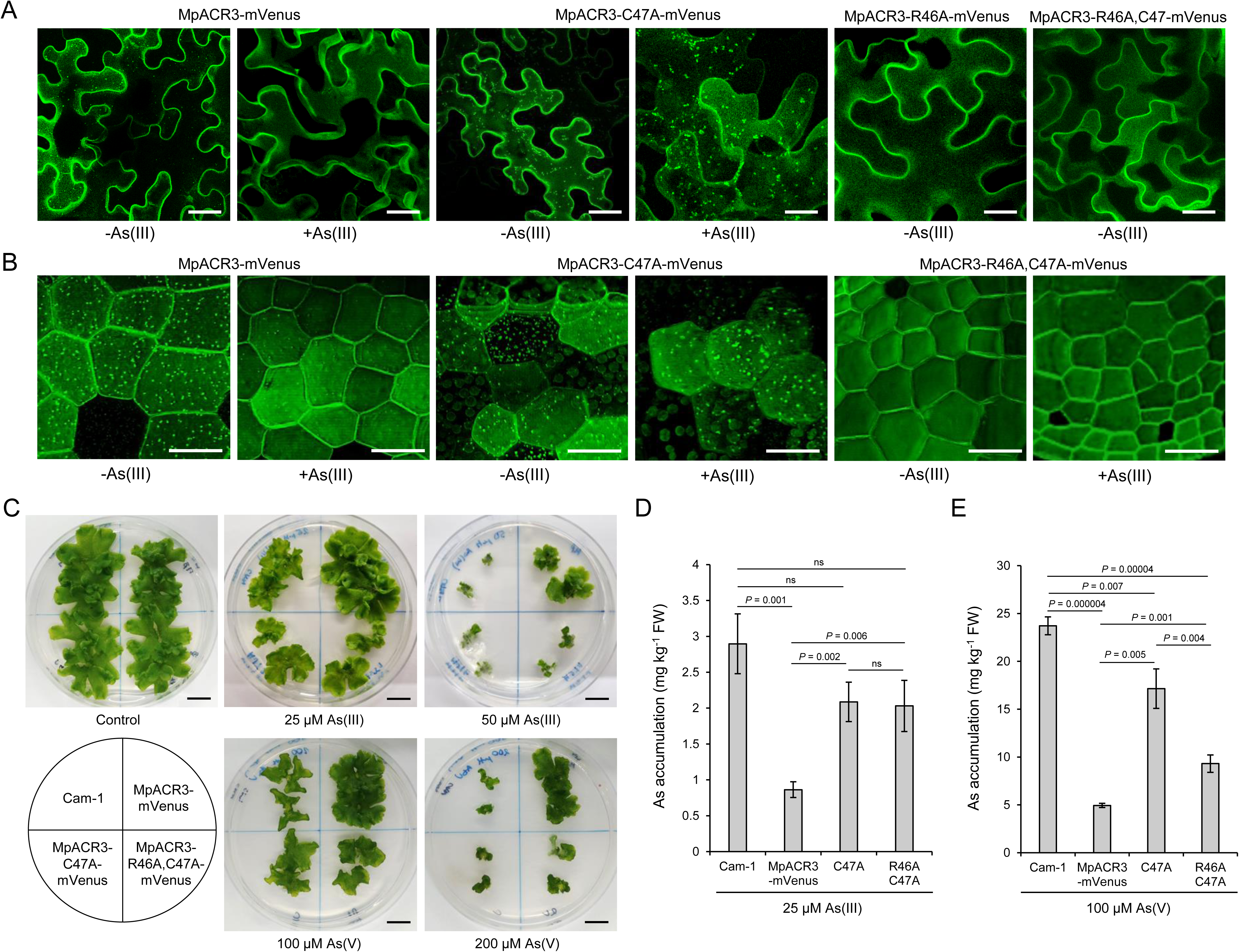
The LRCRF motif orchestrates MpACR3 sorting and function in plant cells**. A)** Tobacco leaf epidermal cells transiently expressing wild-type MpACR3 and its indicated variants tagged with mVenus were visualised by confocal microscopy 3 days after infiltration and then exposed to 100 μM As(III) or left untreated. Scale bars = 30 μm. **B)** The subcellular localisation of MpACR3-mVenus and its indicated variants in *M. polymorpha* transgenic lines in the absence or presence of 25 μM As(III) was visualized using the ZEISS Lattice Lightsheet 7 microscopy. *M. polymorpha* gemma cells are displayed in 3D space. Scale bars = 30 μm. **C)** Comparative growth of the parental Cam-1 and the indicated transgenic MpACR3-mVenus lines grown in the absence or presence of As(III) and As(V) for 3 weeks. Scale bars = 1 cm. **D** and **E)** Arsenic accumulation in the parental Cam-1 and indicated transgenic lines of *M. polymorpha* was measured after exposure to 25 μM As(III) **D)** or 100 μM As(V) **E)** for 3 weeks using an atomic absorption spectrometer. FW, fresh weight; ns, not significant. Error bars represent mean value ± standard deviation of mean (*n* = 3). One-way ANOVA was used to calculate the *P*-value.

## Discussion

Adaptation of plants to arsenic-contaminated environments has required the evolution of specialized mechanisms for arsenic transport, sequestration, and detoxification. These mechanisms enable plants to survive and thrive despite exposure to toxic levels of arsenic in soil and water. In this study, we functionally characterized the arsenite transporter ACR3 from *M. polymorpha* (MpACR3) and identified a unique regulatory strategy employed by this membrane protein. We demonstrated that MpACR3 utilizes its N-terminal cysteine-rich domain as an arsenic-sensing module, which dynamically responds to arsenic exposure to regulate MpACR3 intracellular trafficking and functional activity. Our findings provide new insights into how plants fine-tune arsenic transport at the molecular level and suggest potential strategies for engineering plants with enhanced arsenic tolerance or developing biological arsenic sensors.

Two recent studies have provided preliminary insights into MpACR3 function, suggesting that it plays a key role in arsenic tolerance by mediating As(III) efflux from the cytoplasm to the extracellular space (Dutta et al. 2024; Li et al. 2024). We showed that Mp*ACR3* overexpression in the yeast *S. cerevisiae*, the angiosperm *A. thaliana*, and the liverwort *M. polymorpha* confers resistance to both As(III) and As(V), while decreasing intracellular arsenic accumulation under treatment with either form (Figs. 1, 3 and 5). These findings are consistent with Dutta et al. (2024), who demonstrated that MpACR3 overexpression in the yeast *acr3*Δ strain enhanced As(III) tolerance while reducing intracellular arsenic accumulation. Furthermore, they reinforce the results of Li et al. (2024), who reported that MpACR3 knockout in *M. polymorpha* increased sensitivity to both As(III) and As(V), accompanied by higher arsenic accumulation in transgenic plants exposed to As(V).

Similar to ScAcr3 (Maciaszczyk-Dziubinska et al. 2010; 2011), MpACR3 also provides moderate Sb(III) tolerance in both *S. cerevisiae* and *M. polymorpha* by lowering intracellular Sb(III) levels (Figs. 1 and 5; Supplementary Fig. S6). In contrast, the overexpression of Mp*ACR*3 in *A*. *thaliana* did not lead to enhanced growth under Sb(III) exposure or significant reduction in Sb(III) accumulation in the transgenic plants (Supplementary Fig. S10). A recent study on rice has revealed that, in contrast to As(III), Sb(III) is primarily retained in the roots, with minimal translocation to the shoots (Huang et al. 2024). We hypothesize that this retention might be a result of Sb(III) forming conjugates with phytochelatins and/or glutathione, which seem to be more stable than As(III)-thiol conjugates (Sun et al. 2000; Percy and Gailer 2008). This increased stability may contribute to Sb(III) immobilization in the cytosol and/or its sequestration in the vacuole, resulting in plant resistance to this metalloid. Consequently, the MpACR3-mediated efflux of free Sb(III) may be inadequate to enhance antimony tolerance in transgenic *A. thaliana* plants. Moreover, our previous research demonstrated that heterologous expression of the phytochelatin synthase (*PCS*) gene from wheat (*Triticum aestivum*) or fission yeast (*Schizosaccharomyces pombe*) fully restored Sb(III) resistance in *S. cerevisiae ycf1*Δ cells but only partially rescued the As(III) sensitivity of *acr3*Δ cells (Wysocki et al. 2003). On the other hand, mutation of the *A. thaliana PCS1* gene resulted in increased sensitivity to both As(III) and Sb(III) (Kamiya and Fujiwara 2011). Noteworthy, the human ABC transporter MRP1/ABCC1, which exports metalloid-glutathione conjugates, localized to the PM when expressed in *A. thaliana* and facilitated Sb(III) efflux. However, despite its transport activity, it did not significantly enhance the growth of transgenic plants exposed to Sb(III) (Gayet et al. 2006).

Unlike *A. thaliana*, *M. polymorpha* appears to rely less on phytochelatins for arsenic detoxification. Recent studies have shown that the primary function of the *M. polymorpha* phytochelatin synthase (Mp*PCS*) gene is cadmium detoxification rather than arsenic (Li et al. 2022). Furthermore, our RNA-seq analysis revealed that Mp*PCS* expression remains unchanged in response to As(III) exposure (Supplementary Data Set 1). Additionally, Zhao et al. (2003) reported that phytochelatins play a minimal role in arsenic detoxification in *P. vittata*, as only a small portion (1-3%) of arsenic is complexed with phytochelatins. These findings suggest that phytochelatins contribute only marginally to arsenic tolerance in both *M. polymorpha* and *P. vittata*, with efficient As(III) efflux primarily mediated by ACR3 transporters. This aligns with our previous observations that co-expression of *ScACR3* and *PCS* genes from wheat and *S. pombe* does not enhance arsenic tolerance but instead reduces arsenic resistance levels (Wysocki et al. 2003). The role of phytochelatins in antimony tolerance has yet to be examined in both liverworts and ferns.

ACR3 transporters from bacteria and yeast act as As(III)/H⁺ antiporters (Maciaszczyk-Dziubinska et al, 2011; Villadangos et al, 2012), and our study shows that MpACR3 operates through the same mechanism (Fig. 1K). Like the bacterial CgAcr3 (Fu et al. 2009; Villadangos et al. 2012) and yeast ScAcr3 (Maciaszczyk-Dziubinska et al. 2014; Markowska et al. 2015), MpACR3 relies on highly conserved cysteine (C236) and glutamic acid (E407) residues in transmembrane regions TM4 and TM9, respectively, to mediate arsenic transport activity (Fig. 1, G to J). In CgAcr3 (C129 and E305) and ScAcr3 (C151 and E353), the equivalent sites are believed to play a crucial role in substrate binding, illustrating the remarkable conservation of arsenic transport mechanisms across the ACR3 family. However, eukaryotic ACR3 transporters seem to possess distinct structural and functional characteristics that distinguish them from their bacterial counterparts. In contrast to bacterial Acr3 proteins, ScAcr3 requires an additional glutamic acid residue (E380) in TM10 for proper functioning and exhibits limited Sb(III) transport (Markowska et al. 2015). Moreover, MpACR3 also shows weak Sb(III) transport activity (Fig. 1, D and F; Fig. 5E; Supplementary Fig. S7) and is non-functional in the absence of the corresponding glutamic acid residue (E434) (Fig. 1, I and J).

The ability to detect toxic metal(oid)s or excessive levels of essential metals and rapidly activate or upregulate efflux transporters and downregulate uptake pathways is vital for the survival of all organisms. This is especially critical for land plants, which are immobile and must adapt to their environment. The regulation of membrane transporters includes transcriptional control, trafficking and subcellular localisation, membrane potential and pH modulation, ligand-mediated regulation, and post-transcriptional modifications (Zbieralski et al. 2024; Niño-González and Duque 2025).

In bacteria, expression of As(III) efflux transporter genes located in the *ars* operon is repressed by the action of ArsR repressor under normal conditions (Wu and Rosen 1993). As(III) binding to three vicinal cysteine residues in ArsR triggers the conformation change of ArsR dimer and its release from the promoter, leading to the expression activation of the *ars* operon (Shi et al. 1996). In *S. cerevisiae*, the Yap8 transcription factor, which activates the expression of the *ScACR3* gene (Wysocki et al. 2004), also acts as the As(III) sensor (Kumar et al. 2015). As(III) binding to three cysteine residues in Yap8 leads to its stabilization and activation (Kumar et al. 2015). In contrast to bacteria and yeast, Mp*ACR3* is expressed under normal conditions and its expression is further increased in the presence of arsenic and antimony compounds (Li et al. 2024; Supplementary Fig. S13; Supplementary Data Set 1). The upregulation of Mp*ACR3* expression may be driven by a transcription factor that directly senses arsenic or may stem from the activation of a broader general or oxidative stress response.

Beyond transcriptional regulation, our findings indicate that MpACR3 activity is modulated post-translationally through ligand binding and sorting to the PM. Under normal conditions, MpACR3 localizes to the ER and PM in *S. cerevisiae* cells (Fig. 2) and to the Golgi and PM in plant cells (Fig. 3, E to H; Fig. 4; Fig. 5, F and G; Supplementary Figs. S10 and S11). High-resolution microscopy further pinpoints its specific localisation to the *trans*-Golgi cisternae (Fig. 4). Consistently, Li et al. (2024) reported that the MpACR3-Citrine fusion protein predominantly localizes to the PM, with the additional presence in unidentified cytosolic structures. Upon exposure to arsenic or antimony, MpACR3 rapidly relocates to the PM, losing its retention in the ER or Golgi (Fig. 2; Fig. 3, E and F; Fig. 5F; Fig. 10, A and B; Supplementary Fig. S11). Moreover, we demonstrate that the plant-specific large N-terminal domain governs the intracellular retention of both MpACR3 and the moss *P. patens* PpACR3, but not the fern *P. vittata* PvACR3, which lacks this domain (Fig. 6, A and B). Mutational and biochemical analyses of the MpACR3 N-terminal domain strongly support its role as an As(III) sensor (Figs. 6 and 7; Supplementary Fig. S16). We propose that As(III) binding coordinates three conserved cysteine residues (C29, C47, and C71), triggering a conformational change in the sensor and directing MpACR3 sorting to the PM. MpACR3 variants lacking any of these cysteines, or the conserved F49 residue, which likely stabilizes this conformation, failed to rapidly relocate to the PM in response to As(III) treatment (Fig. 6D; Fig. 9B; Fig. 10, A and B). Although As(III) sensing via cysteine residues is a known mechanism, as seen in bacterial ArsR and yeast Yap8, where As(III) binding triggers conformational changes leading to gene expression regulation (Shi et al. 1996; Kumar et al. 2015; Wysocki et al. 2004), our findings reveal a novel role for this process in the direct control of membrane protein trafficking. MpACR3 uses its N-terminal domain as an As(III) sensor to modulate its own subcellular localisation, a regulatory strategy not previously described in eukaryotic arsenic detoxification systems.

We also investigated how the N-terminal As(III) sensor regulates intracellular retention and identified a highly conserved 45-LRCRF-49 motif, which includes C47 and F49, both essential for MpACR3 relocation to the PM (Figs. 9 and 10). Notably, this sequence aligns with a known arginine-based sorting signal (LRXR), which plays a crucial role in the retention of membrane proteins within the ER and Golgi (Michelsen et al. 2005; Banfield 2011; Lujan and Campelo 2021). This motif has been shown to interact with COPI to facilitate retrograde transport within the Golgi and from the Golgi to the ER (Zerangue et al. 2001; Michelsen et al. 2005; Beck et al. 2009). Consistent with this, single substitution variants MpACR3-L45A, MpACR3-R46A, and MpACR3-R48A, as well as double variants MpACR3-L45A,C47A, MpACR3-R46A,C47A, and MpACR3-C47A,R48A, were exclusively localized to the PM, confirming that the LRXR motif governs MpACR3 intracellular retention in the absence of As(III) (Figs. 9 and 10). The clustering of arginine residues appears to be a key factor in the sorting signal’s interaction with COPI (Zerangue et al. 2001; Michelsen et al. 2005). To further investigate this, we analyzed sequence variations in the LRCRF region across plant ACR3s and performed mutational studies. Our results showed that the first arginine can be substituted with valine without affecting ER retention of MpACR3 in yeast cells (Fig. 9, B and C). Nevertheless, it remains to be tested in both yeast and plant cells whether the MpACR3 N-terminal domain physically interacts with COPI in the absence or presence of As(III) and whether one or both arginine residues are required for this interaction.

Upon As(III)-induced conformational change in the MpACR3 sensor, the LRCR sorting signal may become masked through post-translational modifications, binding of accessory proteins, or interactions with the cytosolic domains of other membrane proteins, including MpACR3 itself (Michelsen et al. 2005; Banfield 2011; Lujan and Campelo 2021). Notably, molecular modelling of the MpACR3 N-terminal domain suggests that As(III) binding to C29, C47, and C71 triggers several structural changes, including: compaction of the overall structure, expansion of the α-helical 35-40 region to 32-45 residues (formed by two discontinuous α-helices) toward the 45-LRCRF-49 motif, resulting in formation of a negatively charged patch on the protein’s surface, and increased conformational flexibility in the 85-95 region (Fig. 8; Supplementary Figs. S17, S21 and S22). Interestingly, these structural changes do not appear to restrict access to the LRCR sorting signal (Supplementary Fig. S24). Instead, they may enhance protein-protein interactions, potentially facilitating MpACR3’s re-localisation.

Intriguingly, while the N-terminal As(III) sensor plays a similar role in MpACR3 intracellular retention in both yeast and plant cells, we observed a key difference in its localisation: MpACR3 is retained in the ER in yeast (Fig. 2) but localizes to the Golgi in plant cells (Figs. 3 to 5). This discrepancy may stem from differences between these organisms in membrane lipid and protein composition, Golgi apparatus structure, the number of COPI subclasses, and the subcellular localisation of enzymes that modify newly synthesized membrane proteins. For example, in *S. cerevisiae*, the Golgi cisternae are dispersed throughout the cytoplasm (Suda and Nakano 2012), whereas in most eukaryotes, they are stacked. Moreover, *S. cerevisiae* possesses a single COPI complex, whereas plants are thought to have two distinct COPI subpopulations, COPIa and COPIb, composed of different subunits. These subpopulations are proposed to mediate ER and Golgi retention, respectively (Donohoe et al. 2007; Gao et al. 2014). Supporting this model, multiple paralogues of COPI subunits have been identified in the *A. thaliana* genome (Gao et al. 2014). In contrast, the *M. polymorpha* genome encodes only a single copy of each COPI subunit, suggesting that distinct COPI subpopulations are unlikely to exist in this species.

Cell surface trafficking of integral membrane proteins can also be regulated by S-palmitoylation, a process in which a C16 acyl chain is added to cytosolic cysteine residues near transmembrane spans. Palmitoylation has been proposed to influence the conformation of transmembrane regions, association with membrane domains, formation of protein complexes, and inhibition of lysine ubiquitylation near the palmitoylation site. Consequently, the absence of palmitoylation or enzymatic depalmitoylation may lead to the intracellular retention of membrane proteins (Blaskovic et al. 2013; Hemsley 2015; Li and Qi 2017). Importantly, we previously proposed that ScAcr3 undergoes palmitoylation at C90, in the proximity of TM2, to prevent ubiquitylation at a neighbouring lysine and subsequent retention in the ER. Mutation of this site led to ScAcr3 retention in the ER, whereas an additional substitution of K91 with arginine suppressed this effect (Maciaszczyk-Dziubinska et al. 2014). Interestingly, these residues are conserved in MpACR3, corresponding to C175 and K176 (Supplementary Figure S2). MpACR3 also contains two additional cysteine residues near TM5 and TM18, C254 (equivalent to C169 in ScAcr3) and C384 (not conserved in ScAcr3), which may also undergo palmitoylation (Supplementary Figure S2). Protein S-acyl transferases (PATs), which catalyse S-palmitoylation, are transmembrane proteins localized to various compartments, including the ER, Golgi, and PM (Ohno et al. 2006; Li and Qi 2017). It is intriguing to speculate that in plant cells, MpACR3 is palmitoylated by a Golgi-localized PAT, with this modification serving as a sorting signal for PM targeting. However, under normal conditions, the N-terminal sensing domain may inhibit palmitoylation of MpACR3, leading to its retention in the Golgi. It would be interesting to investigate whether the absence of MpACR3 palmitoylation results in its ubiquitination and degradation, as we observed increased levels of MpACR3 protein in plant cells in the presence of arsenic (Figs. 3A and 5A).

Functional analysis of LRCR motif variants, which localize exclusively to the PM, revealed that they are partially functional (Figs. 9 and 10). This finding suggests that the As(III)-induced conformational transition of the N-terminal domain not only regulates MpACR3 trafficking but also modulates MpACR3 function at the PM. Interestingly, recent studies have shown that a large cysteine-rich cytoplasmic loop in ScAcr3, found exclusively in species closely related to *S*. *cerevisiae*, plays a regulatory role in As(III) transport by modulating the activity of the aquaglyceroporin Fps1 (Lee and Levin 2022). This regulatory mechanism relies on the physical interaction between the ScAcr3 cytoplasmic loop and the Rgc1/2 regulatory proteins bound to Fps1, facilitating the opening of the Fps1 channel and enhancing As(III) efflux in response to As(V) treatment.

In contrast, under As(III) exposure, methylated As(III) species bind to cysteine residues within the ScAcr3 cytoplasmic loop, causing the release of Rgc1/2 activators from Fps1. This leads to Fps1 channel closure and the inhibition of As(III) uptake (Lee and Levin 2022). It is intriguing to speculate that MpACR3, through its N-terminal domain, may also regulate arsenic influx by modulating the activity of other PM proteins. Similarly, further research is needed to understand the functional significance of MpACR3 presence at the PM in the absence of arsenic (Figs. 3, 5, and 10). Addressing this may require analysing Mp*ACR3* expression from its native promoter.

In summary, our functional analysis of MpACR3 not only uncovers a novel adaptation mechanism to arsenic stress in plants but also holds potential for the development of a biological sensor for detecting As(III) in environmental samples. Furthermore, the MpACR3 arsenic sensing domain may be utilized as a valuable tool for investigating the elusive mechanisms of membrane protein trafficking in eukaryotes.

## Materials and methods

### Yeast material, transformation, and growth conditions

The *S. cerevisiae* strains used in this study include RW104 *(acr3*Δ*::kanMX)* and RW107 *(acr3*Δ*::kanMX ycf1*Δ*::loxP*) (Wysocki et al. 2001) in the W303-1a background (*MAT***a** *leu2-3 112 trp1-1 can1-100 ura3-1 ade2-1 his3-11,15*), as well as the Δ-s-tether strain (*MAT***a** *ice2*Δ*::natMX4 ist2*Δ*::hisMX6 scs2*Δ*::TRP1 scs22*Δ*::hisMX6 tcb1*Δ*::kanMX6 tcb2*Δ*::kanMX6 tcb3*Δ*::hisMX6*), a generous gift from Christopher T. Beh (Simon Fraser University, British Columbia, Canada), in the SEY6210 background (*MAT*α *leu2*-*3,112 ura3*-*52 his3*Δ*200 trp1*Δ*901 lys2*-*801 suc2*Δ*9*) (Quon et al. 2018). The *acr3*Δ*::kanMX* strain expressing the ER marker *proTPI1-SS-mCherry-HDEL* at the *TRP1* locus was generated via transformation with *Pac*I-linearized pZJOM90 plasmid, a gift from Zhiping Xie (Addgene plasmid #133647; http://n2t.net/addgene:133647) (Zhu et al. 2019).

Yeast cells were grown at 30°C in selective synthetic dextrose (SD) medium supplemented with adenine and appropriate amino acids. To assess metalloid sensitivity, yeast cell cultures were serially diluted and plated on selective SD medium containing varying concentrations of sodium arsenite [As(III)], sodium arsenate [As(V)], or potassium antimony tartrate [Sb(III)] (Sigma-Aldrich). For Sb(III)-containing plates the medium pH was adjusted to 7.0 to enhance Sb(III)-induced growth inhibition. Plates were incubated at 30°C and photographed after 2 d. To examine the subcellular localisation of MpACR3-GFP in budding yeast, cells were cultured in liquid SD medium with metalloids or heavy metals for up to 2 h before microscopic observation.

### Plant material and growth conditions

Male gametophytes of *Marchantia polymorpha* Cam-1 and transgenic lines carrying the *pro35Sx2:*Mp*ACR3-mVenus* construct (see Gene cloning section) were cultured on solid 0.5× Gamborg B5 medium (Duchefa Biochemie, pH 5.8) in Petri dishes. Growth conditions included continuous light (150 μmol m^-2^ s^-1^) at 21 ± 1°C in an MLR-352-PE Climatic Test Chamber (PHCbi). For metalloid sensitivity and uptake measurements, gametophytes derived from fresh gemmae were cultured on solid 0.5× Gamborg B5 medium with or without the indicated concentrations of As(III), As(V), or Sb(III) for 3 weeks. In Sb(III)-containing plates, the medium pH was adjusted to 7.0 (Fig. 5B). For confocal microscopy analysis (Fig. 5F), *Marchantia* lines were plated on solid 0.5× Gamborg B5 medium supplemented with 25 μM As(III), and gemmae were imaged 24 h after metalloid or mock treatment. For Lattice Lightsheet 7 (LLS7) microscopy (Fig. 10B), gemmae were incubated directly on glass slides in water or water supplemented with 25 μM As(III) for 20 min before imaging.

All experiments in this study were conducted using the *Arabidopsis thaliana* Columbia-0 (Col-0) ecotype. The following marker lines, previously characterized by Geldner et al. (2009), were obtained from the Salk Institute: Wave9R (VAMP711-mCherry, vacuole marker), Wave22R (SYP32-mCherry, Golgi marker), Wave127R (MEMB12-mCherry, Golgi marker), and Wave138R (PIP1;4-mCherry, plasma membrane marker). The ER marker line mCherry-HDEL (Nelson et al. 2007) was generously provided by Marcela Rojas-Pierce (North Carolina State University, US). The *pro35S:*Mp*ACR3-GFP* transgenic line was generated during this study (see Gene cloning section). For co-localisation studies, the *pro35S:*Mp*ACR3-GFP* T27 line was crossed with the relevant mCherry marker lines described above.

*A. thaliana* plants were cultivated under a 16 h light/8 h dark photoperiod at 22°C, either in soil or in vitro on 0.5× Murashige and Skoog (MS) medium (Sigma-Aldrich) supplemented with 1% sucrose. To compare phenotypic differences between wild-type *A*. *thaliana* and transgenic *pro35S:*Mp*ACR3-GFP* lines (T21 and T27), plants were grown in soil. Upon completing their growth cycle, measurements were taken for main stem height, number of flowers on the main stem, total number of branches, and total number of flowers. For the germination assay, sterilized seeds from wild-type and *pro35S:*Mp*ACR3-GFP* transgenic lines were sown on MS medium with or without 100 µM As(III) or 700 µM As(V) and allowed to grow for 14 d. For the growth assay, wild-type and transgenic *pro35S:*Mp*ACR3-GFP* seeds were initially sown on MS medium without arsenic. After 4 d, seedlings were transferred to MS medium supplemented with either 25 µM As(III), 1000 µM As(V), or no arsenic (control) and grown for an additional 7 d. For confocal microscopy (Fig. 3, E to J) and Lattice Lightsheet 7 (LLS7) microscopy (Supplementary Fig. S11), double heterozygous plants (heterozygous for both the *mCherry* reporter and the *pro35S:*Mp*ACR3-GFP* transgene) were first grown on MS medium without arsenic. After 4 d, seedlings were transferred to MS medium with or without 25 µM As(III) or 1000 µM As(V) and incubated for one additional day before imaging.

### Quantitative RT-PCR analyses

The gametophytes were grown on solid 0.5× Gamborg B5 medium for 2 weeks in the presence or absence of As(III), As(V), or Sb(III). Material for total RNA isolation was collected by cutting off the terminal fragments of *M. polymo*rpha gametophytes with a sterile scalpel. Total RNA was extracted from 50 mg of plant tissues using the Beadbeat Total RNA Mini kit (A&A Biotechnology, Gdansk, Poland). First-strand cDNA from RNA samples was then synthesized using the High-Capacity cDNA Reverse Transcription Kit (Applied Biosystems TM). Quantitative PCR (qPCR) was performed using cDNA as a template, the 2× PCR Master Mix SYBR kit (A&A Biotechnology, Gdansk, Poland), and the CFX Connect Real-Time PCR Detection System (Bio-Rad, Hercules, CA, US) in a total volume of 15 μl. Primers used for qPCR are listed in Supplementary Table S2. The amplification conditions were as follows: 1 min at 95°C, 40 cycles of 10 s at 95°C, 15 s at 60°C, and 20 s at 72°C. Finally, melting-curve analysis was conducted to verify the specificity of the reaction. The comparative threshold cycle (CT) method for relative quantification (ΔΔCT method) was used to analyse the data.

### RNA sequencing and analysis

*M*. *polymorpha* Cam-1 plants were grown under control conditions (solid 0.5x Gamborg B5 medium) or treated with 2.5 μM As(III) for a week. Samples were collected using a scalpel and ground in liquid nitrogen. Each condition was tested in three independent experiments. Total RNA was extracted using the RNeasy Plant Mini Kit following the manufacturer’s instructions (Qiagen, Germany). Library preparation and RNA sequencing were performed by Novogene Europe. Sequencing reads were mapped to the *M*. *polymorpha* MpTak v6.1 assembly (http://marchantia.info/) using HISAT2 (Kim et al. 2015). Gene expression counts were obtained with HTSeqCount (Anders et al. 2015), and differential expression analysis was conducted using EdgeR (Robinson et al. 2010).

### Gene cloning

The oligonucleotides used for *ACR3* cloning and verification are listed in Supplementary Table S2, while the plasmids utilized in this study are detailed in Supplementary Table S3. To clone the Mp*ACR3* and *PpACR3* genes for heterologous expression in yeast, total RNA was isolated from *M. polymorpha* Cam-1 and sterile explants of *Physcomitrium patens* ssp. *patens* (Gransden 2004 ecotype), courtesy of Agnieszka Hanaka and Piotr Waśko (Maria Curie-Skłodowska University, Lublin, Poland). RNA extraction was performed using the BeadBeat Total RNA Mini Kit (A&A Technology), and the resulting RNA was used as a template for reverse transcription with the Takara PrimeScript RT Reagent Kit. The Mp*ACR3* and *PpACR3* coding sequences were amplified by PCR (CloneAmp HiFi PCR Premix, Takara) using the appropriate cDNA as a template. The amplified sequences were cloned into the pUG35 vector, fused to a C-terminal GFP tag, and placed under the control of the constitutive *MET17* promoter using the In-Fusion HD Cloning Kit (Takara). The resulting pMpACR3 and pPpACR3 plasmids were verified by DNA sequencing. Additionally, the synthetic DNA sequence of the *PvACR3* gene from *P. vittata* (Thermo Fisher Scientific) was cloned into pUG35 using the same approach. For comparison, the pScACR3 plasmid, which overexpresses the *S. cerevisiae proMET17:ScACR3-GFP* fusion gene, was used as described previously (Maciaszczyk-Dziubinska et al. 2014).

Plasmids encoding the N-terminally swapped variants, N_Sc_-MpACR3-GFP and N_Mp_-ScACR3-GFP, were constructed through recombination. This was achieved by combining PCR-generated linear fragments of pMpACR3 or pScACR3, each lacking the respective N-terminal region of ACR3, with fragments encoding the N-terminal tail of ScACR3 or MpACR3, respectively, using the In-Fusion HD Cloning Kit. Similarly, the plasmid encoding the MpACR3-Δ27-98-GFP variant was generated by fusing a PCR-amplified pMpACR3 linear fragment that omits the sequence encoding the N-terminal cysteine-rich region.

For heterologous expression in yeast, the wild-type Mp*ACR3* N-terminal domain (MpACR3-N-tail) and its cysteine-lacking variants (MpACR3-AAA-N-tail and MpACR3-Cys-null-N-tail) were cloned into the *Sal*I-digested pGREG535 vector (EUROSCARF, P30360). These constructs enabled expression under the *GAL1* promoter and tagging with a 7×HA epitope at the N-terminus. For bacterial expression and subsequent protein purification, DNA sequences encoding the wild-type and mutant variants of the Mp*ACR3* N-terminal domain were synthesized and cloned into the pGEX4T-1 plasmid by BioCat GmbH. The resulting plasmids, pGST-MpACR3-N-tail, pGST-MpACR3-AAA-N-tail, and pGST-MpACR3-Cys-null-N-tail, allowed for the inducible expression of N-terminally GST-fused proteins in *E. coli* BL21(DE3) cells.

For plant expression, the synthetic Mp*ACR3* gene sequence was cloned under the control of the *35Sx2* promoter, fused to a C-terminal mVenus tag (*pro35Sx2:*Mp*ACR3-mVenus* and its mutant variants). This was achieved using Type IIS Loop assembly and components from the OpenPlant toolkit (Pollak et al. 2019; Sauret-Güeto et al. 2020). The same strategy was used to generate the *pro35Sx2:*Mp*ACR3-mVenus*, *proUbE2:52aaST-mScarlet* construct for the *Marchantia* co-localisation experiment. Additionally, the Mp*ACR3* coding sequence was amplified from the pMpACR3 plasmid using high-fidelity PCR and cloned into the *Xba*I-digested pGFPGUSPLUS vector (Vickers et al. 2007; a gift from Claudia Vickers; Addgene plasmid #64401), resulting in the *pro35S:*Mp*ACR3-GFP* construct.

### Mutagenesis

Plasmids carrying the Mp*ACR3* gene or its mutant variants served as templates for site-directed mutagenesis, performed using the QuikChange Lightning Kit (Agilent Technologies) to generate single and multiple mutants. The introduced mutations were verified by DNA sequencing. Oligonucleotides used for mutagenesis are listed in Supplementary Table S2.

### Transformation

Yeast transformation was carried out using the lithium acetate/single-stranded carrier DNA/polyethylene glycol method (Gietz and Woods 2002).

For *M. polymorpha* transformation, *Agrobacterium tumefaciens* cells (GV3103) were electroporated with the *pro35Sx2:*Mp*ACR3-mVenus* construct, its mutant variants, or the *pro35Sx2:*Mp*ACR3-mVenus, proUbE2:52aaST-mScarlet* construct using a Bio-Rad Gene Pulser DNA. The transformed bacteria were plated on LB medium containing spectinomycin, rifampicin, and gentamycin, then incubated at 29°C for 3 d. *M. polymorpha* spores (Cam-1 x Cam-2) were sterilized and plated on 0.5× Gamborg B5 medium for 5 d, following the protocol described by Sauret-Güeto et al. (2020). The 5-day-old spores were then collected using a scalpel blade and resuspended in 4 mL of liquid 0.5× Gamborg B5 media in sterile 6-well plates. The medium was supplemented with 0.1% N-Z amino A (Sigma-Aldrich, C7290) 0.03% (w/v) L-glutamine (Alpha Caesar, A14201) 2% (w/v) sucrose (Fisher Scientific, 10634932), and 100 μM acetosyringone. A plate containing transformed *Agrobacterium* colonies was streaked with a sterile pipette tip, and approximately 0.3 mm diameter sphere of bacterial cells was resuspended in the liquid medium containing the *M. polymorpha* sporelings. The 6-well plate was shaken at 120 rpm for 2 d with continuous lighting (150 μmol m^-2^ s^-1^). Following co-cultivation, the sporelings were transferred onto a 70 μm-pore cell strainer and washed with 10 mL of sterile water. They were then placed on solid 0.5× Gamborg B5 medium supplemented with 100 μg/mL cefotaxime (Apollo Scientific, BIC0111) and 20 μg/mL hygromycin (Invitrogen, 10687010). Transformants emerged within a week and were subsequently transferred to fresh plates for gemma cup formation.

The *pro35S:*Mp*ACR3-GFP* construct was electroporated into *A. tumefaciens* cells GV3101 (pMP90) (Koncz and Schell 1986) and used to transform *A*. *thaliana* (Col-0) plants via the floral dip method (Clough and Bent 1998). Seeds from transformed *A*. *thaliana* plants were surface-sterilised and sown on MS medium supplemented with 25 μg/mL Hygromycin B (Sigma-Aldrich). After exposure to light for 6 h, the seeds were kept in the dark for four days before being transferred to a 16-h light/8-h dark photoperiod for further growth. Transformants, identified by their elongated hypocotyls and green true leaves, were then transferred to soil. Seven T1 plants were obtained, and lines T21 and T27, due to the strongest fluorescence of their *pro35S:*Mp*ACR3-GFP* transgene product, were selected for more detailed analyses.

### Transient expression

Transient expression of the Mp*ACR*3 gene in tobacco leaf epidermal cells was carried out as described in (Hawes et al. 2024).

### Protoplast isolation

Protoplasts were isolated from *A*. *thaliana* leaves using a modified Tape-*Arabidopsis* Sandwich method (Wu et al. 2009). Young leaves from 3 to 5-week-old plants were affixed to a strip of autoclave tape, and the lower epidermal surface cell layer was carefully peeled off using price tape. The leaves, still attached to the autoclave tape, were then transferred to a Petri dish and submerged in 2 mL of enzyme solution (1% cellulase ‘Onozuka’ R10, 0.25% macerozyme, 0.4 M mannitol, 10 mM CaCl_2_, 20 mM KCl, 20 mM MES, pH 5.7). The samples were incubated in the dark at room temperature for 1 h with gentle shaking (70 rpm). Protoplasts were collected by centrifugation at 100g for 4 min at 4°C, then washed twice with 2 mL of cold W5 solution (154 mM NaCl, 125 mM CaCl_2_, 5 mM KCl, 5 mM glucose, 2 mM MES, pH 5.7) and centrifuged again under the same conditions. Finally, the protoplast-containing pellet was resuspended in 300 μL of cold MMg solution (0.4 M mannitol, 15 mM MgCl_2_, 4 mM MES, pH 5.7).

### FM4-64 staining

To label the vacuolar membrane in live yeast cells, 0.5 mL of an overnight yeast culture was washed, resuspended in SD medium containing 0.04 mM FM6-64 (Molecular Probes), and incubated in the dark for 2 h in a 30°C water bath. The cells were then analysed using an epifluorescence microscope.

For plasma membrane staining in live *pro35S:MpACR3-GFP Arabidopsis* cells, seedlings were transferred to a 0.08 mM FM4-64 staining solution, incubated on ice for 2 min, and then sequentially washed in two cold washing solutions to remove excess FM4-64. The protoplast-containing pellet was suspended in diluted FM4-64 stock solution (0.04 mM in MMg solution) and incubated on ice for 2 min. The reaction was stopped by washing the protoplasts with cold MMg solution. The protoplasts were then collected by centrifugation at 100g for 4 min at 4°C and resuspended in MMg solution (Rigal et al. 2015).

### Microscopy

The subcellular localisation of MpACR3-GFP fusion proteins in live yeast cells was analysed using an upright wide-field epifluorescence microscope (Axio Imager M2, Carl Zeiss, Germany) equipped with a Zeiss HBO 100 illuminator and a 100× oil immersion objective (Zeiss Plan-Neofluar 100×/1.30). Fluorescence signals were detected using either filter set 38 HE (GFP) or filter set 43 HE (mCherry, FM4-64). Images were captured with a Zeiss AxioCam MRc digital camera and processed using Zeiss ZEN 3.7 software.

A Leica SP8 confocal microscope was used to image *M. polymorpha* sporelings and gemmae (Fig. 5, F and G). A 460-670nm WLL (White Light Laser) was used to excite mVenus (515 nm) and mScarlet (560 nm), with fluorescence signals detected via HyD detectors. Laser power and gain were adjusted per sample, ranging from 5-15% and 50-150, respectively. Imaging was conducted with a Plan-Apochromat 40× water immersion objective.

An Olympus FluoView FV1000 confocal laser scanning microscope was used to image MpACR3-GFP (excitation: 488 nm) in *A. thaliana* protoplasts (Supplementary Fig. S10) and root cells expressing Wave-mCherry, HDEL-mCherry, or FM4-64 markers (excitation: 559 nm) (Fig. 3, E to J). Imaging was performed using a Plan-Apochromat 60× objective.

A Lattice Lightsheet 7 (LLS7) fluorescence microscope was employed for visualizing MpACR3-mVenus (488 nm laser) variants in *M. polymorpha* gemmae (Fig. 10B) and MpACR3-GFP (488 nm laser) in *A. thaliana* root cells co-expressing the Golgi marker (Wave127R) with mCherry (565 nm laser) (Supplementary Fig. S11). Imaging parameters included: Sinc3 100 x 1400, z-step 0.3 µm, 488 nm laser power 4%, exposure time 30 ms for MpACR3-mVenus or Sinc3 30 x 1000, z-step 0.3 µm, 488 nm laser power 2%, 565 nm laser power 6%, exposure time 50 ms for MpACR3-GFP/Wave127R-mCherry. Images were processed using ZEN ZEISS 3.10 (Carl Zeiss Microscopy GmbH), applying medium-strength deconvolution, linear interpolation, deskewing, cover glass transformation, and image subset selection. Final images were displayed in 3D space.

A Leica STELLARIS 8 confocal microscope was used to capture MpACR3-mVenus signals in tobacco leaf epidermal cells (Fig. 10A). A 460-670nm WLL (White Light Laser) was used to excite mVenus, and fluor bescence signals were detected using HyD detectors. Laser power and gain were adjusted per sample, ranging from 5-20% and 50-100, respectively. Imaging was performed with a Plan-Apochromat 40× water immersion objective.

Co-localisation of MpACR3-GFP with the marker proteins of Golgi compartments in tobacco leaf epidermal cells was visualised using Airyscan high-resolution confocal microscopy 3 d post-infiltration (Fig. 4). The analysis methodology was based on (McGinness et al. 2022; McGinness et al. 2025).

### Co-localisation analysis

We quantitatively analysed tracer co-localisation by calculating Pearson’s correlation coefficients (PCC) for individual cells using the co-localisation tool in Olympus FV1000 Viewer. A PCC value of 1 indicates perfect correlation, while a value of 0 signifies no correlation. PCC calculations were performed on images from three independent biological replicates, with a total of 15 cells analysed.

### Metalloid uptake measurements

Yeast transformants were cultivated in selective SD medium until reaching the logarithmic phase, then incubated with varying concentrations of As(III) or Sb(III) for a specified duration. Following optical density measurement at 600 nm (OD_600_), cells were washed 3 times with cold deionized water and lysed in 2 mL of 0.5% nitric acid (EMPLURA, Merck) by boiling for 10 min. After centrifugation, the supernatants were analysed for arsenic or antimony concentrations using an atomic absorption spectrometer (AAS) with a graphite cuvette (contrAA 800 G, Analytik Jena).

For plant samples, a similar procedure was followed. *A. thaliana* and *M. polymorpha* were grown on solid MS or 0.5× Gamborg B5 medium, respectively, supplemented with different concentrations of As(III), As(V), or Sb(III) for 14 or 21 d. Tissue fragments were weighed and snap-frozen in liquid nitrogen before homogenization in 1.5 mL of 0.5% nitric acid using glass beads and a Precellys 24 homogenizer (Bertin Technologies). The homogenized samples were boiled, centrifuged, and the metalloid concentrations in the supernatants were measured as described above.

### Metalloid/H^+^ antiporter assay

Insight-out plasma membrane vesicles were isolated from yeast transformants pre-treated with 0.1 mM of As(III) for 2 h using a dextran/polyethylene glycol two-phase partitioning method (Norling, 2000). Plasma membrane protein content was quantified using the Bio-Rad Bradford Assay.

The As(III)/H^+^ antiport activity was assessed using acridine orange as a pH-sensitive probe. Membrane vesicles (50 μg) were added to 500 μL of assay buffer containing 10 mM MES-Tris (pH 6.0), 330 mM sucrose, 140 mM KCl, 4 mM MgCl_2_, 0.1 mM EDTA, 1 mM DTT, 50 mM KNO_3_, 1 mM of NaN_3_, 0.1 mM Na_2_MoO_4_, and 5 μM acridine orange. Formation of the ΔpH gradient was initiated by adding 2 mM ATP, and the resulting change in acridine orange absorbance at 495nm (A495) was monitored using a Beckman DU640 spectrophotometer. After 3 min, ATP-dependent H^+^-transport activity was inhibited by adding 0.5 mM sodium orthovanadate.

As(III)/H^+^ antiporter activity was determined by its ability to dissipate the preformed, inside-acid pH gradient following the addition of 10 mM As(III) or either 10 mM or 20 mM Sb(III). The acridine orange absorbance (A495) measured after proton gradient formation and immediately after As(III) or Sb(III) addition was normalized across samples. Subsequent changes in A495 were recorded at various time points to quantify gradient dissipation. The generation of the proton gradient was consistent across plasma membrane vesicles from different yeast strains, while As(III)-mediated dissipation of the gradient was representative of results obtained separately for each sample.

### Total protein extracts

Total protein extraction from yeast cells was performed using the trichloroacetic acid (TCA) method (Wright et al. 1989). For *A. thaliana* and *M. polymorpha*, total protein extracts were obtained following the protocol of Martínez-García et al. (1999). Proteins were resolved on 10% SDS-PAGE, transferred to nitrocellulose filters (Bio-Rad), and probed with an anti-GFP antibody (Roche, 11814460001) at a 1:3000 dilution to detect GFP or mVenus-tagged ACR3 proteins in yeast and plant samples. For total protein normalization, nitrocellulose filters were stained with Ponceau S solution (Sigma-Aldrich). Additionally, an anti-3-phosphoglycerate kinase (Pgk1) antibody (Abcam, 22C5D8) at a 1:3000 dilution was used as a loading control for yeast proteins.

### Fractionation

Logarithmic-phase yeast cultures were centrifuged, and the resulting cell pellets were lysed using glass beads in a lysis buffer (20 mM Tris-HCl, pH 7.5, 2 mM MgCl₂, 250 mM sorbitol) supplemented with 1 mM PMSF and a protease inhibitor cocktail (Sigma-Aldrich). Glass beads and cell debris were then removed by centrifugation at 950g for 10 min. Whole-cell extracts were further fractionated by differential centrifugation: first at 7000 for 10 min to obtain an ER/nuclear-enriched pellet, followed by centrifugation at 15,000g for 40 min to isolate a plasma membrane-enriched pellet. The remaining supernatant contained the cytosolic fraction. Pellets were resuspended in MS buffer (2 mM MgCl₂, 10 mM imidazole, 1 mM PMSF, and a protease inhibitor cocktail). The obtained fractions (plasma membrane-enriched, ER/nuclear-enriched, and cytosolic) were solubilized in Laemmli buffer containing 8 M urea, separated by SDS-PAGE, and analysed by Western blot. MpAcr3-GFP was detected using an anti-GFP antibody (Roche, 11814460001). As a quality control for membrane fractionation, the plasma membrane H⁺-ATPase Pma1 was detected using rabbit anti-Pma1 antibodies (kindly provided by Prof. Ramon Serrano) at a 1:15,000 dilution.

### MpACR3 N-terminal domain expression in yeast and trypsin susceptibility assay

Yeast *acr3*Δ cells transformed with pGREG535 plasmids encoding MpACR3 N-terminal domain variants were grown overnight in selective SD medium supplemented with 2% raffinose. The following day, approximately 10^8^ cells from each transformant were transferred to selective SD medium containing 4% galactose and incubated for 3 h at 30°C to induce expression. Cells were then either left untreated or exposed to 0.5 mM As(III) for 1 h before being centrifuged, washed with phosphate-buffered saline (PBS), and resuspended in 600 μL of lysis buffer containing PBS, a protease inhibitor cocktail (Sigma-Aldrich, P8215), and 1 mM PMSF (BioShop). Lysis was performed using 0.5 mm glass beads (Linegal Chemicals) with 3 rounds of bead-beating (Bertin Technologies) at 4000g for 30 s each. Whole-cell extracts were then centrifuged at 850g for 5 min at 4°C, and the resulting lysate was transferred to a fresh tube.

For the trypsin susceptibility assay, 100 μg of yeast lysates was incubated on ice for 10 min with trypsin (Sigma-Aldrich) at concentrations of 0, 1, or 5 μg/mL. Proteolysis was halted by adding 0.5 μg/mL soybean trypsin inhibitor (Sigma-Aldrich) for 15 min. Protein degradation patterns were assessed via Western blot analysis using an anti-HA antibody (Sigma-Aldrich, H6908) at a 1:5000 dilution.

### As-biotin pulldown assay

Lysates of *acr3*Δ yeast cells expressing 7HA-MpACR3-N-tail, 7HA-MpACR3-AAA-N-tail, or 7HA-MpACR3-Cys-null-N-tail from the pGREG535 plasmid were prepared as described above. For the arsenic-biotin pull-down assay, 200 μg of yeast lysate was incubated with 20 µM As-biotin (N-Biotinyl *p*-Aminophenyl Arsenic Acid, Santa Cruz, SC-503570) in 10 mM MES-Tris buffer (pH 6.0) at 25°C for 1 h. In parallel, an additional set of lysates was pre-treated with 20 µM As(III) in 10 mM MES-Tris buffer (pH 6.0) for 30 min before the addition of the As-biotin conjugate. Following incubation, lysates were diluted 10-fold with 10 mM MES-Tris buffer (pH 6.0) and mixed with 30 µl of streptavidin-agarose resin (Thermo Scientific), which had been pre-equilibrated with 0.05% Tween 20 buffer (one wash) and 10 mM MES-Tris buffer (pH 6.0) (two washes). Samples were incubated at 21°C for 1 h with continuous overhead rotation. The resin was then briefly spun down and sequentially washed with NaCl buffer (1 M NaCl, 10 mM MES-Tris buffer, pH 6.0) for 2 min, followed by urea buffer (2 M urea, 10 mM MES-Tris buffer, pH 6.0) for 2 min. As-protein conjugates were eluted using 45 μl of Laemmli buffer (250 mM Tris-HCl, pH 6.8, 500 mM DTT, 10% SDS, 0.01% bromophenol blue, 50% glycerol) after incubation at 95°C for 5 min. All fractions, including the input lysate (not incubated with As-biotin) and the eluate, were denatured at 95°C for 10 min before Western blot analysis. A total of 40 µL of each fraction was analysed by 12% SDS-PAGE, transferred to a nitrocellulose membrane (Bio-Rad), and immunoblotted using an anti-HA antibody (Sigma-Aldrich, H6908) at a 1:5000 dilution.

### Purification of MpACR3 N-terminal domain and thermolysin digestion

*Escherichia coli* BL21(DE3) cells carrying pGST-MpACR3-N-tail, pGST-MpACR3-AAA-N-tail, or pGST-MpACR3-Cys-null-N-tail plasmids were cultured at 37°C until reaching A600 of 0.8. Protein expression was then induced by adding 0.5 mM IPTG, followed by incubation at 30°C for 4 h. Cells were harvested by centrifugation, and the resulting pellet was lysed by sonication (Bandelin Sonopuls) in lysis buffer (PBS, 10 mM 2-mercaptoethanol, 1% Triton X-100, 10% glycerol, protease inhibitors, 1 mM PMSF). Sonication was performed in 10 cycles of 30 s each. The lysate was clarified by centrifugation at 21,000g for 30 min at 4°C. Protein extracts were incubated with Glutathione Sepharose™ 4B resin (GE Healthcare) pre-equilibrated with PBS in a gravity-flow column. After three washes with PBS, bound proteins were eluted using an elution buffer containing 10 mM reduced glutathione (L-glutathione, Sigma-Aldrich) and 50 mM Tris-HCl (pH 8.0). The eluted proteins were concentrated using Amicon® Ultra-15 Centrifugal Filter Units (Merck Millipore) and stored at −80°C until further use.

A stock solution of thermolysin from *Geobacillus stearothermophilus* (Sigma-Aldrich, T7902) was prepared at 2.5 mg/mL in proteolytic buffer (0.1 mM TCEP, 25 mM bis-Tris, pH 6.0). Before the proteolysis assay, the enzyme was further diluted 50-fold in proteolysis buffer. For digestion, 5 µg of purified protein was mixed with 0.5 mL of the diluted thermolysin solution in a final volume of 5 mL on ice, then incubated at 21°C. The reaction was terminated by adding an equal volume of hot 4X SDS sample buffer supplemented with 2 mM PMSF and 100 mM DTT, followed by heat denaturation at 95°C for 5 min. The samples were immediately loaded onto an SDS-PAGE gel for analysis. For MpACR3 variants, the thermolysin stock solution was diluted 100-fold before addition to the reaction mixture.

### Phylogenetic analysis of ACR3 protein sequences

ACR3 protein sequences were retrieved from multiple databases, including NCBI, GigaDB, FernBase, Hornworts Database, Dryad, Phytozome, and Phycocosm. Detailed sequence information is provided in Supplementary Table S1. Sequence alignment was performed using ClustalW 2.1, embedded in Geneious Prime (version 2019.1.1; Biomatters Ltd.), with default settings. Phylogenetic relationships were inferred using the maximum likelihood method via the IQ-Tree web server (Trifinopoulos et al. 2016). Bootstrap analysis with 1000 replicates was conducted to assess branch support values. The resulting phylogenetic tree was visualized using Interactive Tree of Life (iTOL3) (Letunic and Bork 2019).

### Molecular modelling

The MpACR3 AlphaFold model structure (Entry ID: A0A176WR38) (Jumper et al. 2021) was utilized, focusing on the 24–105 amino acid region for further investigation. The structure of this region was initially prepared using the Maestro BioLuminate Schrödinger suite (v. 2021-3) (Salam et al. 2014). This preparation resulted in two disulphide bonds (C47-C71 and C29-C96), which were considered improbable due to the cytosolic localization of the domain. To correct this, we manually edited the 24–105 MpACR3 structure using USCF Chimera (Pettersen et al. 2004), where we removed the disulphide bonds and added hydrogen atoms to the corresponding SG atoms. Additionally, a triple alanine mutant (C29A, C47A, C71A) was generated using Chimera’s rotamer functionality. Subsequently, unrestrained molecular dynamics (MD) simulations were performed for both the wild-type (WT) and triple alanine mutant (Triple Ala) structures using GROMACS MD engine (version 2021.4) (Abraham et al. 2015). The Amber14sb force field (Maier et al. 2015) was applied for protein treatment, while OPC force field parameters (Izadi et al., 2014) were used for water modelling, following the standard simulation protocol.

We first neutralised the systems and then solvated in triclinic rectangular periodic boxes, ensuring a buffer distance of 15 Å to the walls. Additional Na^+^ and Cl^-^ ions were introduced to achieve a final concentration of 150 mM NaCl. We treated ions with the Joung-Cheatham parameters (Joung and Cheatham 2008). Applying periodic boundary conditions, we first energy minimized the systems with 5,000 steps of steepest decent. This was followed by 500 ps equilibration-runs with weak position restraints on heavy atoms of the solute (1,000 kJ/mol) in the NVT and NPT ensembles, adjusting temperature and pressure to 300 K and 1 atm, respectively (Parrinello and Rahman 1981; Berendsen et al. 1984). Finally, after lifting the restraints, we conducted 2 μs MD simulations for each system under constant pressure (1 atm) and temperature (300 K).

To investigate a potential conformational transition induced by As(III) binding to C29, C47, and C71, we performed an additional steered MD simulation. In this simulation, we gradually reduced the distances between the SG atoms of the cysteine residues, maintaining them within a range of 3.5-3.8 Å, which correspond to SG-SG distances observed when As(III) is bound. This distance range was derived from the structural analysis of available crystal structures in which As(III) is coordinated by three cysteine residues.

Using the PLUMED software (Bonomi et al. 2009), we applied a distance bias through a wall quadratic potential to constrain the accessible configurational space during the simulation. The upper and lower wall limits were set at 3.8 Å and 3.5 Å, respectively. Starting with a force constant of 500 kJ/mol*nm^2, we incrementally increased it by 500 kJ/mol*nm^2 every 200 ns until the target distance range was reached at a force constant of 2500 kJ/mol*nm^2. The restraints were then maintained for an additional 2000 ns to allow the protein structure to equilibrate.

We analysed the structural dynamics of the three simulated systems: WT, triple Ala, and WT with restraints, using built-in analysis tools in the GROMACS MD v. 2021.4 package. Our analysis included the RMSD of all heavy atoms, RMSF for each residue, and secondary structure elements. To examine the evolution of secondary structure elements, we utilized the DSSP algorithm. Additionally, we assessed protein electrostatic surface potential and surface hydrophobicity using UCSF Chimera. For visualizing protein topology, we employed the PDBsum web server (Laskowski et al. 2018). Molecular graphics were generated using UCSF Chimera, while data plotting was performed in MATLAB.

## Supporting information

Supplementary Data Set 1

## Acknowledgments

The authors gratefully acknowledge Christopher T. Beh for generously providing the Δ-s-tether strain, Marcela Rojas-Pierce for the ER marker line mCherry-HDEL, Agnieszka Hanaka and Piotr Waśko for supplying sterile *P. patens* explants, and Facundo Romani for his assistance with RNA-seq analysis. The authors also thank the Swedish National Infrastructure for Computing (SNIC) for the generous provision of computing resources. During the preparation of this manuscript, the authors used ChatGPT to improve readability and language.

## Author contributions

K.M., I.B., R.W., and E.M.D. designed the research. K.M., K.Z., E.M.D., and D.W. conducted yeast experiments. I.B. generated and analysed RNA-seq transcriptomic data. I.B. also constructed *M. polymorpha* transgenic lines. I.B. and P.T. analysed MpACR3 subcellular localisation in *M. polymorpha*. A.D. constructed *A. thaliana* transgenic lines and performed growth analyses. A.D. and P.T. performed microscopy analyses in *A. thaliana*. P.T., W.B., and V.K. performed transient expression experiments in tobacco leaf cells. P.T. and W.B. analysed MpACR3 subcellular localisation in tobacco leaf cells. V.K. acquired and analysed high-resolution confocal microscopy data for MpACR3 subcellular localisation. K.M., E.M.D., and J.S. measured metalloid cellular content and conducted the antiporter assay. E.M.D. carried out qPCR analysis, As-biotin pulldown, trypsin susceptibility assays, MpACR3 protein level analysis in *M. polymorpha* and *A. thaliana* transgenic lines, and growth tests of *M. polymorpha*. D.W. performed the fractionation experiment. W.B. purified the MpACR3 N-terminal domain and performed thermolysin digestion assays. P.T. constructed the phylogenetic tree. A.R. conducted molecular modelling and data analysis. R.W. generated multiple sequence alignments. All authors contributed to data interpretation and manuscript drafting. R.W. wrote the final version of the manuscript and obtained funding. E.M.D., J.H., and R.W. supervised the project. All authors have read and approved the final manuscript.

## Data availability

All data and materials that support the findings of this study are included in the manuscript or in the Supplementary Materials. Sequencing data have been deposited in the Gene Expression Omnibus (GEO) repository at the National Center for Biotechnology Information (NCBI) under the accession number GSE276247.

