## Supplementary Figures and Tables for "Arsenic-sensing domain controls ACR3 transporter trafficking and function in *Marchantia polymorpha*"

**Supplementary Figures S1 to S24**

**Supplementary Tables S1 to S3**

### Arsenic-sensing domain controls ACR3 transporter trafficking and function in *Marchantia polymorpha*

Katarzyna Mizio[1], Ignacy Bonter[1,2], Kacper Zbieralski[1], Alicja Dolzblasz[3], Paulina Tomaszewska[1,4], Jacek Staszewski[1], Donata Wawrzycka[1], Anna Reymer[5,6], Wojciech Bialek[7], Verena Kriechbaumer[8], Jim Haseloff[2], Robert Wysocki[1,*], Ewa Maciaszczyk-Dziubinska[1,*]

[1]Department of Genetics and Cell Physiology, Faculty of Biological Sciences, University of Wroclaw, 50-328 Wroclaw, Poland

[2]Department of Plant Sciences, University of Cambridge, Downing Street, Cambridge, CB2 3EA, United Kingdom

[3]Department of Plant Developmental Biology, Faculty of Biological Sciences, University of Wroclaw, 50-328 Wroclaw, Poland

[4]Department of Genetics, Genomics and Cancer Sciences, University of Leicester, LE1 7RH, Leicester, United Kingdom

[5]Department of Chemistry and Molecular Biology, University of Gothenburg, 405 30 Göteborg, Sweden

[6]Institute for Advanced Biosciences (Inserm U1209), Site Santé – Domaine de la Merci, 38700 La Tronche, France

[7]Department of Biophysics, Faculty of Biotechnology, University of Wroclaw, 50-383 Wroclaw, Poland

[8]School of Biological and Medical Sciences and Centre for Bioimaging, Oxford Brookes University, Oxford OX3 0BP, United Kingdom

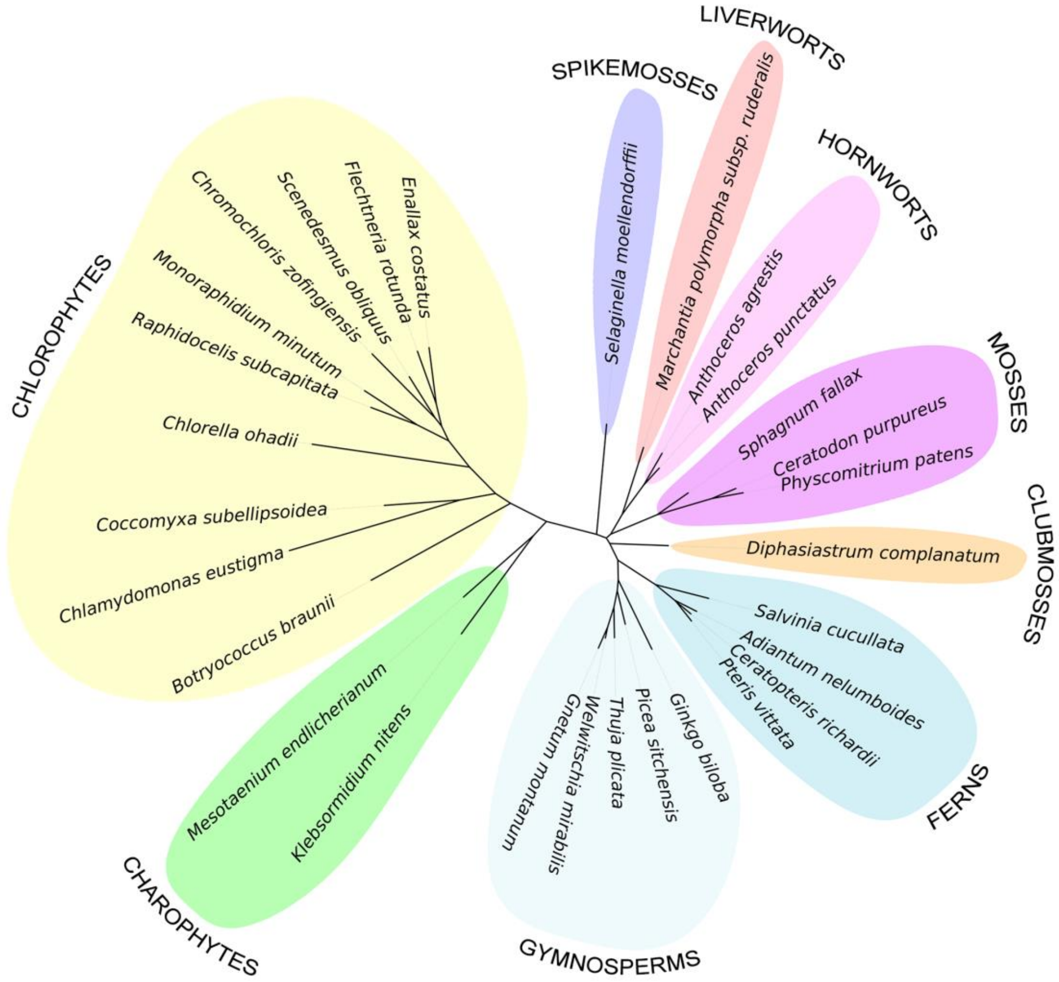


**Supplementary Figure S1.** Phylogenetic tree of the plant members of the ACR3 family. The sequences were aligned using ClustalW 2.1. The phylogenetic relationship was inferred by the maximum likelihood method. 1000 bootstrap replicates were generated to estimate branch support values. Sequence details are provided in Supplementary Table S1.

```
MpACR3  MRGSEMTEMEAERPVDNTGLEAGEVVVKCKPLWDDARTGSAKDELRCRFSKESRNVDVPV  60
PvACR3  ------------------------------------------------------------  0
ScAcr3  ------------------------------------------------------------  0
CgAcr3  ------------------------------------------------------------  0

MpACR3  VDGDNAEGFLCKCSAELSST--GTTVSILLPNGDELSCEALPDSDKVKGVVTAGQIYKQL  118
PvACR3  --------------MENSSAERKQQLALDIADGNDPS----DAAKNPDGRTKLQGLFKQL  42
ScAcr3  ------------------------------MSEDQKSENSVPSKVNMVNRTDILTTIKSL  30
CgAcr3  ----------------------------------------------MTNSTQTRAKPARI  14

                      TM1                              TM2
MpACR3  SLLDRYLFVWIFAVMGLALLIGYYVDGIKKRLS---VVEIASVSLPIAIGLWVMMYPVLC  175
PvACR3  SLLDRYLYVWIFIVMAVSIIFGYYVKGVKKAFQ---VAEITSVSLPIAIGLWVMMYPVLC  99
ScAcr3  SWLDLMLPFTIILSIIIAVIISVYVPSSRHTFDAEGHPNLMGVSIPLTVGMIVMMIPPIC  90
CgAcr3  SFLDKYIPLWIILAMAFGLFLGRSVSGLSGFLG---AMEVGGISLPIALGLLVMMYPPLA  71

                                TM3                       TM4
MpACR3  KVRYELLYKMLRRGRLGKNLAISLVLNWLVGPALMTGLAWATLPDLSDYRTGVILVGIAR  235
PvACR3  KVQYEILGGVLRQAGSLKTISLSVVLNWVVGPALMTGLAWATLPDLPDFRTGVILVGIAR  159
ScAcr3  KVSWESIHKYFYRSYIRKQLALSLFLNWVIGPLLMTALAWMALFDYKEYRQGIIMIGVAR  150
CgAcr3  KVRYDKTKQIATD---KHLMGVSLILNWVVGPALMFALAWLFLPDQPELRTGLIIVGLAR  128

           TM4                  TM5
MpACR3  CIAMVLIWNELAKGDAQLCAIIVAVNSVLQIILFTPVSLFYLKVISGGKGI-----DVGV  290
PvACR3  CIAMVLIWNDLAKGDADYCAILVAINSILQIILFTPVALLYLKVVSRGKGF-----HVSS  214
ScAcr3  CIAMVLIWNQIAGGDNDLCVVLVITNSLLQMVLYAPLQIFYCYVISHDHL--NTSNRVLF  208
CgAcr3  CIAMVLVWSDMSCGDREATAVLVAINSVFQVAMFGALGWFYLQVLPSWLGLPTTTAQFSF  188

                 TM6                                      TM7
MpACR3  WTVAKSVLLFLGVPLLAGFLTRVILRRAAGARWYDTKFLPFIGPWALLGLLYTIFVMFAI  350
PvACR3  WTVAKSVLLFLGVPLAAGVLTRLILMNAFGRKWYESKFLRFIGPWALIGLLYTIFVMFSI  274
ScAcr3  EEVAKSVGVFLGIPLGIGIIIRLGSLTIAGKSNYEKYILRFISPWAMIGFHYTLFVIFIS  268
CgAcr3  WSIVTSVLVFLGIPLLAGVFSRIIGEKIKGREWYEQKFLPAISPFALIGLLYTIVLLFSL  248

                              TM8
MpACR3  QSHEIVNNIGHVLRVVVPLLLYFSITFTSAVAVCRYFNL---------------------  389
PvACR3  QAHQIVDNIGHVVRVAVPLLLYFGILFFGSLGICRWLKV---------------------  313
ScAcr3  RGYQFIHEIGSAILCFVPLVLYFFIAWFLTFALMRYLSISRSDTQRECSCDQELLLKRVW  328
CgAcr3  QGDQIVSQPWAVVRLAIPLVIYFVGMFFISLIASKLSGM---------------------  287

                                TM9                       TM10
MpACR3  -------PYPEIVTQSFTAASNNFELAIAVAVGAFGIDSHEALAATIGPLIEVPVLLILV  442
PvACR3  -------PYPLMVTQCFTAASNNFELAIAVAVGSFGIDSTQALAATIGPLIEVPVLLLFV  366
ScAcr3  GRKSCEASFSITMTQCFTMASNNFELSLAIAISLYGNNSKQAIAATFGPLLEVPILLILA  388
CgAcr3  -------NYAKSASVSFTAAGNNFELAIAVSIGTFGATSAQAMAGTIGPLIEIPVLVGLV  340

MpACR3  YVTLYFRKWLSKEQA---------------  457
PvACR3  YIVGFFQRKGPSV-----------------  379
ScAcr3  IVARILKPYYIWNNRN--------------  404
CgAcr3  YAMLWLGPKLFPNDPTLPSSARSTSQIINS  370
```

**Supplementary Figure S2.** Multiple sequence alignment of ACR3 proteins. CgAcr3, *Corynebacterium glutamicum* (NCBI accession WP_011265757.1); ScAcr3, *Saccharomyces cerevisiae* (NCBI accession DAA11615.1); PvACR3, *Pteris vittata* (NCBI accession ADP20955.1); MpACR3, *Marchantia polymorpha* (NCBI accession OAE35577.1). Predicted transmembrane regions (TM) are marked with black bars. Conserved residues that are crucial for arsenite transport activity and analysed in this work are highlighted in black. Multiple sequence alignment was generated using the Clustal Omega server (https://www.ebi.ac.uk/Tools/msa/clustalo/). Transmembrane regions were determined using the Consensus Constrained TOPology prediction (CCTOP; http://cctop.ttk.hu).

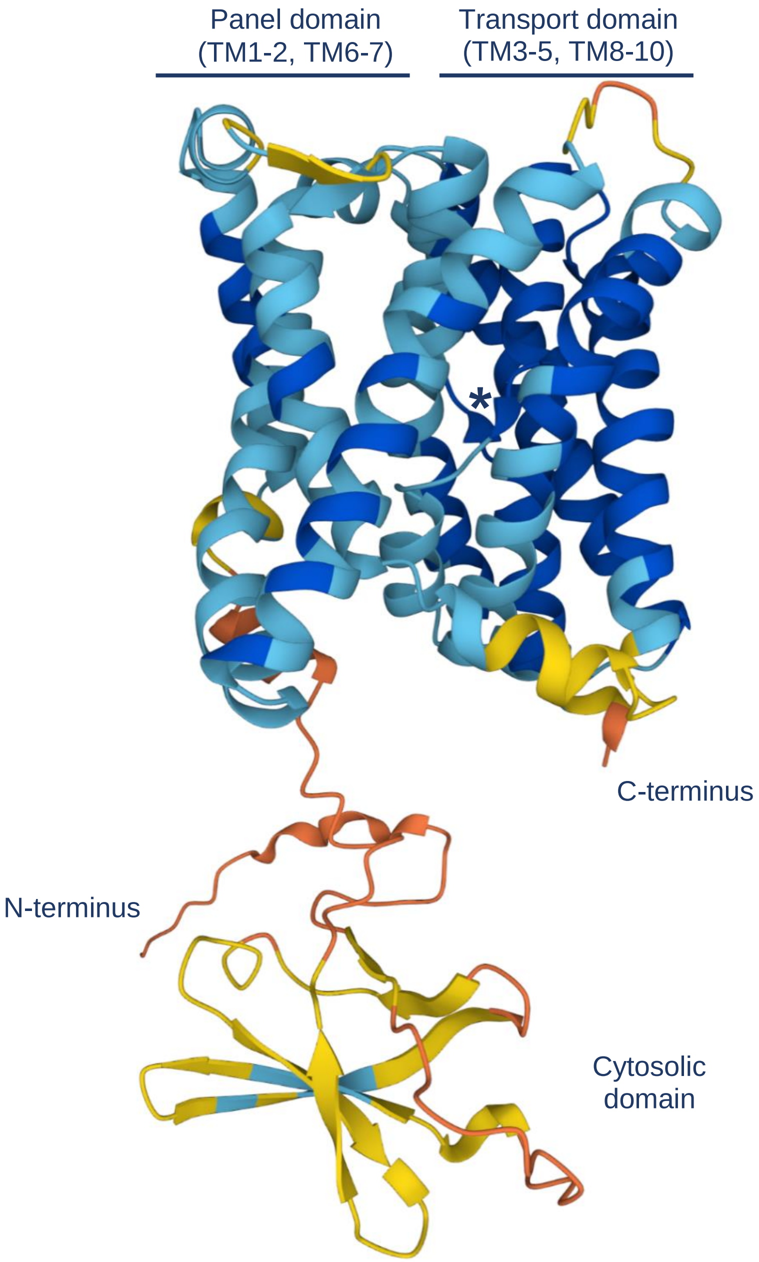


**Supplementary Figure S3.** AlphaFold-predicted structure of MpACR3 (Entry ID A0A176WR38). The structure was coloured based on the AlphaFold per-residue confidence score (pLDDT), where the intervals of confidence are defined as follows: very high (pLDDT > 90, dark blue colour), high (90 > pLDDT > 70; blue colour), low (70 > pLDDT > 50, yellow colour) and very low (pLDDT < 50; peach colour). The asterisk indicates the TM4-TM8 crossover region.

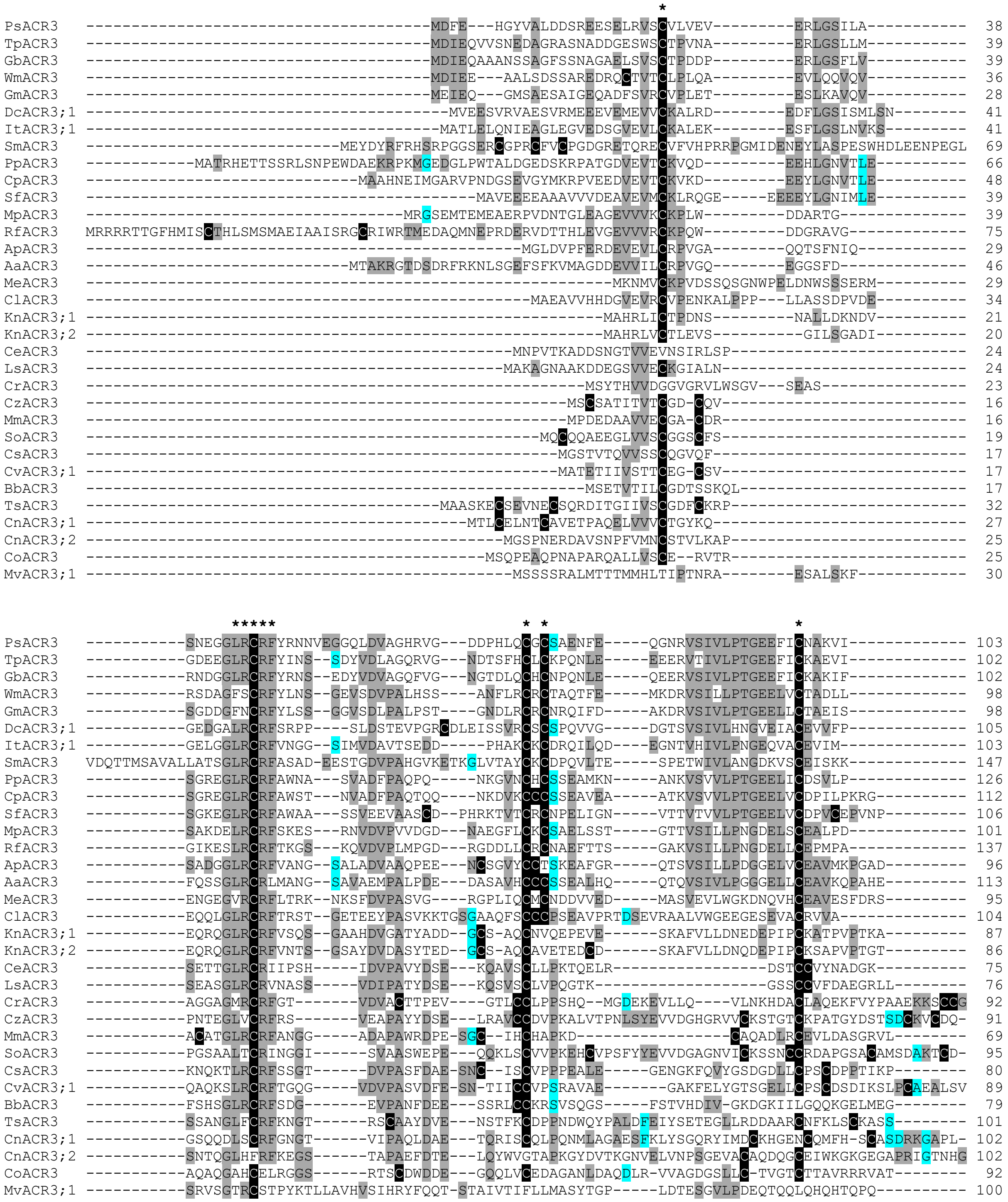

**Supplementary Figure S4.** Multiple sequence alignment of long (residues >100), cysteine-rich N-terminal tails of plant ACR3 proteins. At least 25% sequence identity in a column is highlighted. Cysteine residues are highlighted in black. Conserved residues of MpACR3 analysed in this work are marked with asterisks. Multiple sequence alignment was generated using the Clustal Omega server (https://www.ebi.ac.uk/Tools/msa/clustalo/) and manually adjusted. Sequence details are provided in Supplementary Table S1.

```
PsACR3   -----------------------------TDSYKHEK-EDKSIIRRLSLLDR 125
TpACR3   -----------------------------SGIRKHEK-EEKGLFQRLSFLDR 124
GbACR3   ------------------------------DSGNHGK-EEQRIFQRLSFLDR 123
WmACR3   -----------------------------AEKCNHEK-QGRSIFERLSILDR 120
GmACR3   ------------------------------ESSKHEN-AERGIFSRLSFLDR 119
DcACR3;1 -----------------------------ENGLETEK-SNGKIIKGLSFLDR 127
ItACR3;1 ---------------------------------PSEKHVTGGVYKKLSFLDR 122
SmACR3   ----------------------------------PEK--EEAVYAKLSFLDR 163
PpACR3   -----------------------------SKPGADVP-NVGSVYKRLSFVDR 148
CpACR3   -----------------------------SDTVK-----VASVYKRLSFIDR 130
SfACR3   -----------------------------SDRVTKEK--STPIYNRLSLLDR 127
MpACR3   -----------------------------SDKVKGVV-TAGQIYKQLSLLDR 123
RfACR3   -----------------------------ADEVKEVV-AGGEIFRKLSLLDR 159
ApACR3   ------------------------------------DSAAAGLFKKLSLLDR 112
AaACR3   ------------------------------------KAVAVGLFKKLSLLDR 129
MeACR3   -------------------EAATTSSAAAGIGDGVSSTTSPSIIKQLSFLDR 128
ClACR3   -------------------------ALEEKNAPAVKELSAAAVFRSMSLLDR 131
KnACR3;1 ------------------------LDIGSAEIQADSSTSFKAVVGALGWLDR 115
KnACR3;2 ----------------------FETEPVGQEASASSPANFRSVVGALGWLDR 116
CeACR3   --------------------SVPSDAAKLDDDGEDDNDLEKDPLISLSFVDR 105
CrACR3   -----SCGPALSAEPSGPLATVAISQRALLAPPAAAATSDDSWLAGLSWTDR 142
LsACR3   ---------------------GRSAEHSDSPPTKVDALDEMDPLMSLGIVDR 104
CzACR3   ------------------AQVIDPKQDGTGGDPPTPAAGVGAVLSKLSWLDR 125
MmACR3   -----------YRGPAAGAKAPGAGAGADHADAPAAAAAAPPPPRLPWLDR 110
SoACR3   AKTCDEPFLMNGGGSCCGDKGADAAPGAAPTAAAAGAAAGRGVMSGLSWLDR 142
CsACR3   -------------GVPSSSSIFEVDGCGAGEDGQGSATTATSVFKKLSLLDR 119
CvACR3;1 ---------------------DQGDAPERDRITQPCNTAAGGVLKQLSFLDR 120
BbACR3   -----------PKEAILPGQPEEPPISAIKAEDEPDEVSGSAILKKLSWIDR 120
TsACR3   ----------VPSGIHSFGIVRAPQIGLTAEDTCLPSDPPKSVIKSLSWLDR 143
CnACR3;1 -----------SCASDRKGAPLEIGATNESTDSP-SSVSAIGLIRSLSFLDR 131
CnACR3;2 -----KGEGAPRIGTNHEGCASVLAISGNGEEKEAPPVSAKSVFKQMSFLDQ 136
CoACR3   --------GSLAVAAPPEAGGGSSEVDPAAEADEAAATPSGAVMHGLSWVER 136
MvACR3;1 -----------------------------PDEQCAQP-QFRGLFRKLSILDR 122
```

**Supplementary Figure S4 continued.** Multiple sequence alignment of long (residues >100), cysteine-rich N-terminal tails of plant ACR3 proteins.

```
CrACR3   -----------------------------MEEQKKAHVAAMESSNSYPKEMSISISNGQSDTAAPSSAGATGDEGRKLKGLFKQLSLLDR 61
AnACR3   ------------------------------MENSPPRQKQAADHSPSPNEKQMALD--IPDADKPS-L---PSESTKLQGLFKQLSVLDR 54
PvACR3   ------------------------------MENSSAERKQ-----------QLALD--IADGNDPSDAAKNPDGRTKLQGLFKQLSLLDR 47
SaACR3   ------------------------------MNTES---------SDFPN-----LENGR--PHSQERDNVPPDG--KSKGLFWQLSLLDR 42
AsACR3   ------------------------------MEISSVE-KPVAIDVKSPDNDAAAAA----QMQSSSRPPPEP-ER-KLKGLFGQLSLLDR 53
MvACR3;2 ------------------------------MDPYTH---PDPEPGSLHKEDPQQL-----QHQAPHSQPDEQ-CDQQFRGLFRKLSILDR 52
DcACR3;2 MGKMAPCVAGFPDHPRLIARRCPAVAIWKFLLFGAHVVPRFLQVVGDGTSVSIVLHNGVEIACEVVFPENGMDTEKSNGKIIKGLSFLDR 90
ItACR3;2 ----------------------------------------------MSKVLLAEDKTVHIVLPTGEEIACEAFEKDASGGVYKKLAFLDR 44
CvACR3;2 --------------------MPDETPSFKLFSGKKYIMDCKDGNDCHILHRCSAGGKGGPLEIDANKQVTDGSSDVTAIGLIKSLSFLDR 70
SuACR3   --------------------------------------------MSQQSAVELARKDILPADPDSLGTKKPISEDMGGLSLLASLSFLDR 46
RsACR3   ------------------------------------------MPPKHDKSGSGGHRSGEQAGLDAAEQGGAPAAAGGALARAAALSWLDR 48
FrACR3   -----------------------------MPVSIVLNLFVCPATDQPEILEATDDANISSEGIGAIGTPPHNAAKPKKSAVLGGLSWLDR 61
EcACR3   ---------MVAATRSVCRDEPSSGCAKSPVKTCDEASLLGSAPLAKDPDSLDASMKGDASKCPAAASIEGPPPATASSSVLAGLSWIDK 81
CnACR3;3 MEQWLDWPTEESNCISCVPPPEALQHDDGKFKVYGHDGEVLCSSCDPTIKPSSCCSPPAFEVQSSEAGEEDRNGSVTAGGVFKKLSWLDC 90
```

**Supplementary Figure S5.** Multiple sequence alignment of short (less than 100 residues) N-terminal tails of plant ACR3 proteins. At least 25% sequence identity in a column is highlighted. Cysteine residues are highlighted in black. Multiple sequence alignment was generated using the Clustal Omega server (https://www.ebi.ac.uk/jdispatcher/msa/clustalo) and manually adjusted. Sequence details are provided in Supplementary Table S1.

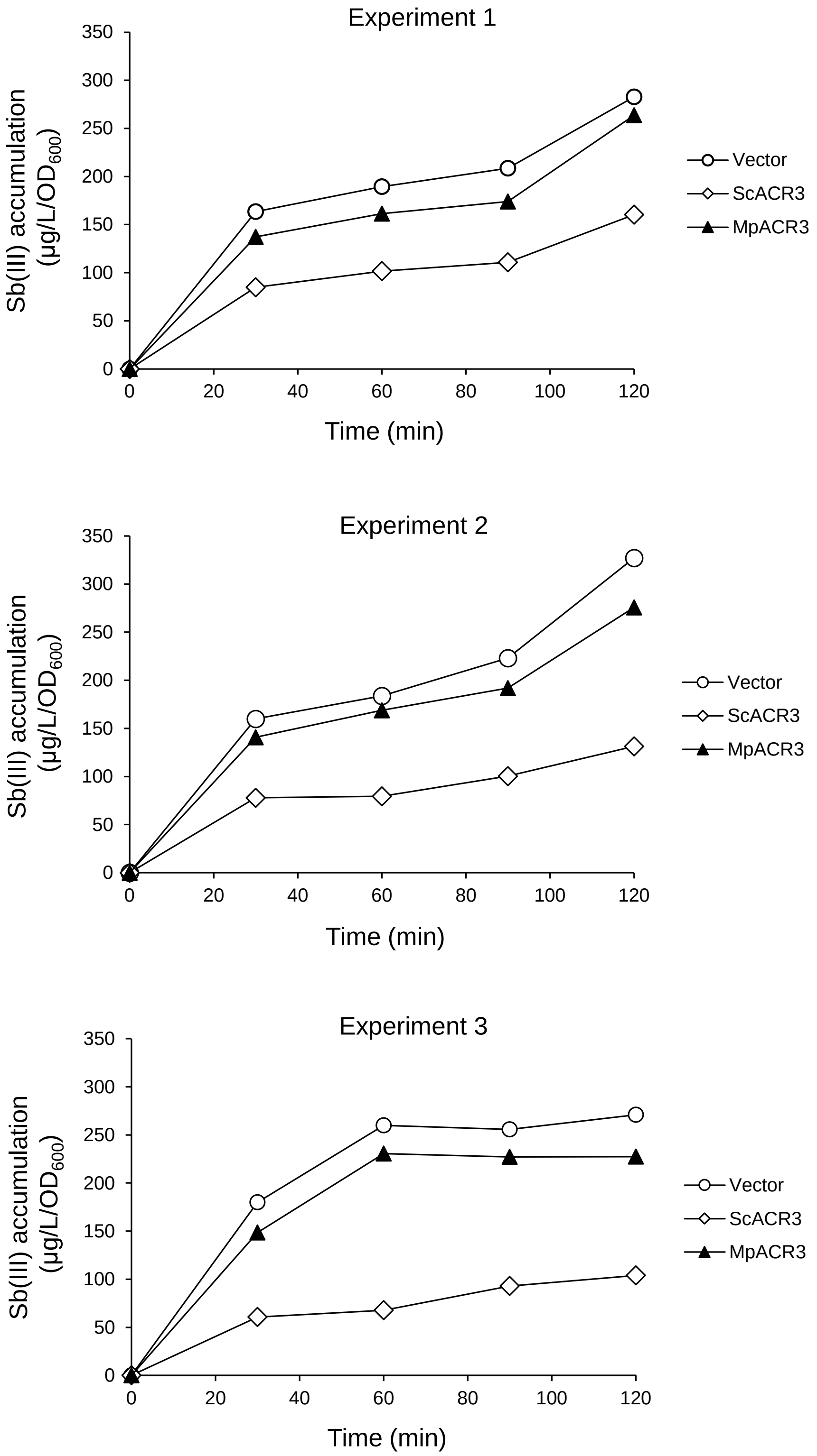


**Supplementary Figure S6.** Three independent experiments of Sb(III) accumulation in the *acr3*Δ mutant containing a control vector, pScACR3 or pMpACR3 plasmid (supports Fig. 1).

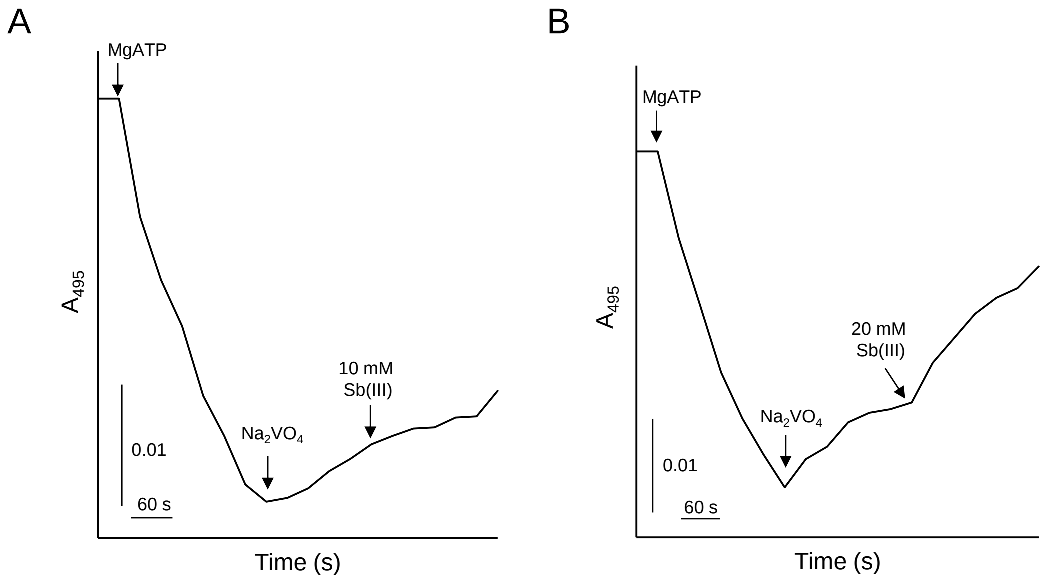


**Supplementary Figure S7.** MpACR3 shows an Sb(**III**)/H$^+$ antiport activity only with a very high concentration of antimony (supports Fig. 1). Sb(**III**)-induced H$^+$ transport across the membranes of inside-out vesicles prepared from *acr3*Δ cells expressing MpACR3-GFP was monitored by measuring changes in the absorbance of acridine orange used as a pH-sensitive probe. Vesicle acidification was initiated by the addition of 2 mM of ATP (first arrow), accompanied by a decrease of absorbance. After 3 min, 0.5 mM of sodium orthovanadate was added to inhibit H$^+$-ATPase and maintain a steady-state acidic-inside pH gradient (second arrow). At the indicated time point, Sb(**III**) was added at the concentration of 10 mM **A)** or 20 mM **B)** to initiate Sb(**III**)-dependent proton movement (third arrow), which acidified the environment and recovered the absorbance.

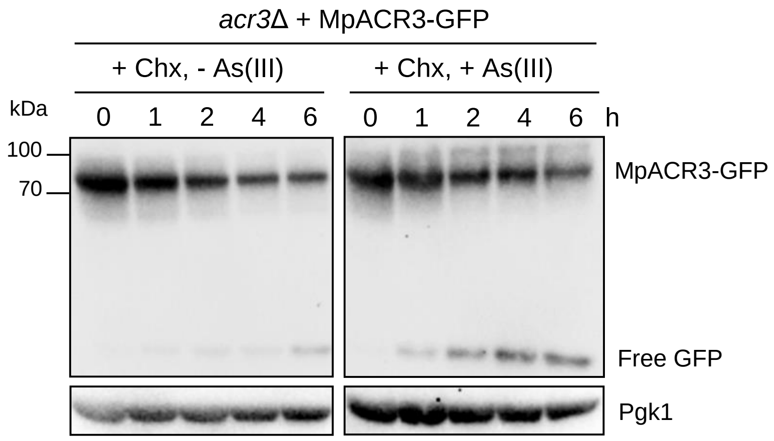


**Supplementary Figure S8.** Stability of the MpACR3-GFP protein in the absence or presence of 0.1 mM As(III) in yeast cells (supports Fig. 2). At time point zero 0.1 mg/ml cycloheximide (Chx) was added to inhibit protein synthesis. Total protein extracts were analysed by Western blot using the anti-GFP antibody to detect MpACR3-GFP. Anti-Pgk1 Western served as a loading control.

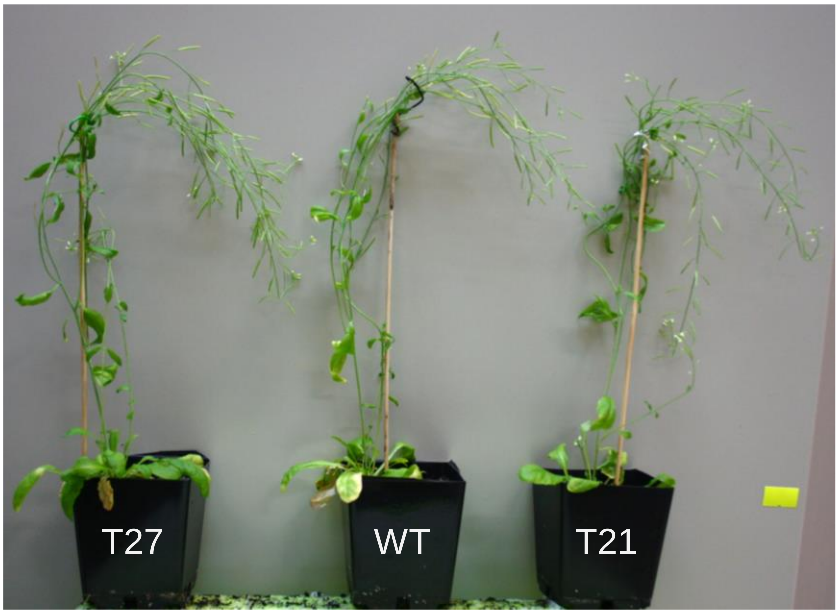


| Line | Main stem height (cm) | Number of flowers on main stem | Total number of branches | Total number of flowers | Number of analysed plants |
|---|---|---|---|---|---|
| WT | 30.4 ± 2.9 | 27.6 ± 4.5 | 3.7 ± 1.0 | 76.9 ± 15.1 | 22 |
| T27 | 29.3 ± 4.4 | 29.1 ± 5.7 | 3.9 ± 0.9 | 82.4 ± 25.5 | 21 |
| T21 | 33.1 ± 4.8 | 32.9 ± 6.8 | 3.2 ± 0.9 | 87.4 ± 24.3 | 21 |
| (±) standard deviation of mean | | | | | |

**Supplementary Figure S9.** No phenotypic differences between *pro35S:*Mp*ACR3-GFP* transgenic lines (T21 and T27) and control wild-type (WT, Col-0) *A. thaliana* (supports Fig. 3). Plants were grown on soil in LD (long-day, 22°C). Indicated growth parameters were measured/counted after plants terminated their growth.

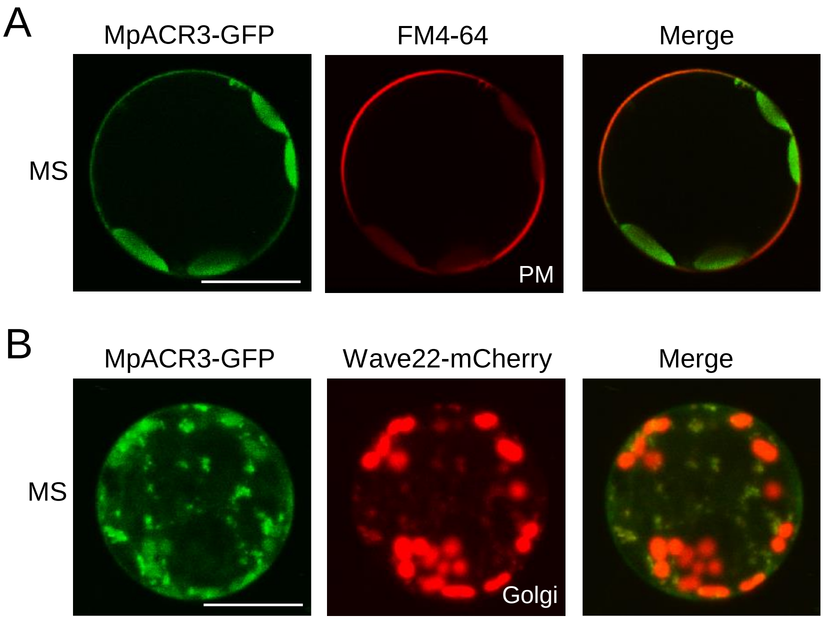


**Supplementary Figure S10**. Subcellular localisation of MpACR3-GFP in *A. thaliana* protoplasts (supports Fig. 3). MpACR3-GFP co-localized with the FM4-64-stained plasma membrane (PM) **A)** and Golgi bodies labelled with the Golgi marker Wave22-mCherry **B)** in protoplasts isolated from leaves under normal conditions (MS medium)**.** Scale bars = 20 μm.

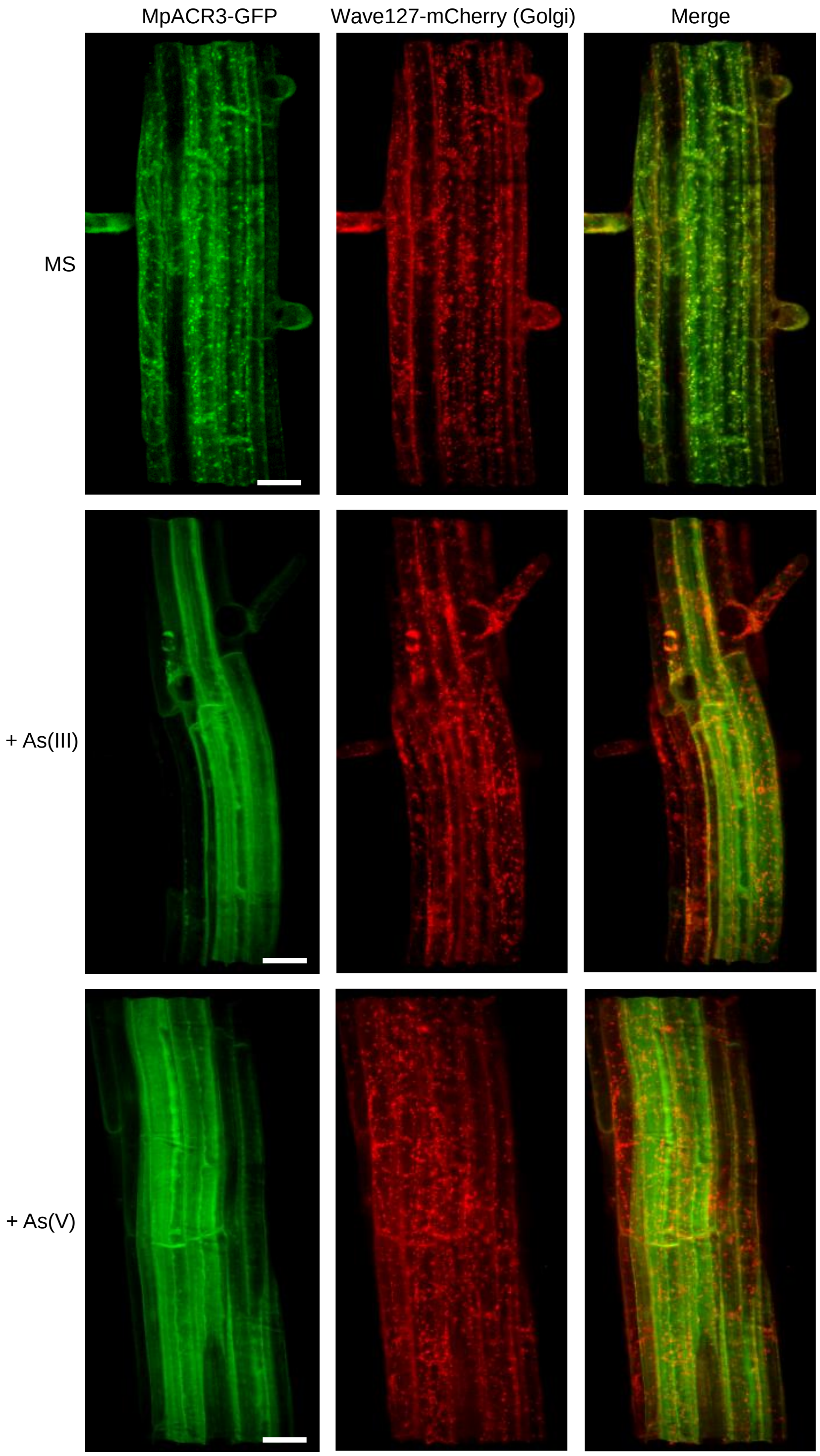


**Supplementary Figure S11.** Imaging of *A. thaliana* roots expressing Mp*ACR3-GFP* and the Golgi marker (Wave127-mCherry) with the use of the ZEISS Lattice Lightsheet 7 microscopy (supports Fig. 3). Roots are displayed in 3D space. 4-day-old seedlings was transferred to MS medium without or with 25 µM As(III) or 1000 µM As(V) and allowed to grow for an additional 1 day before the ZEISS Lattice Lightsheet 7 microscopy analysis. Scale bars = 50 µm.

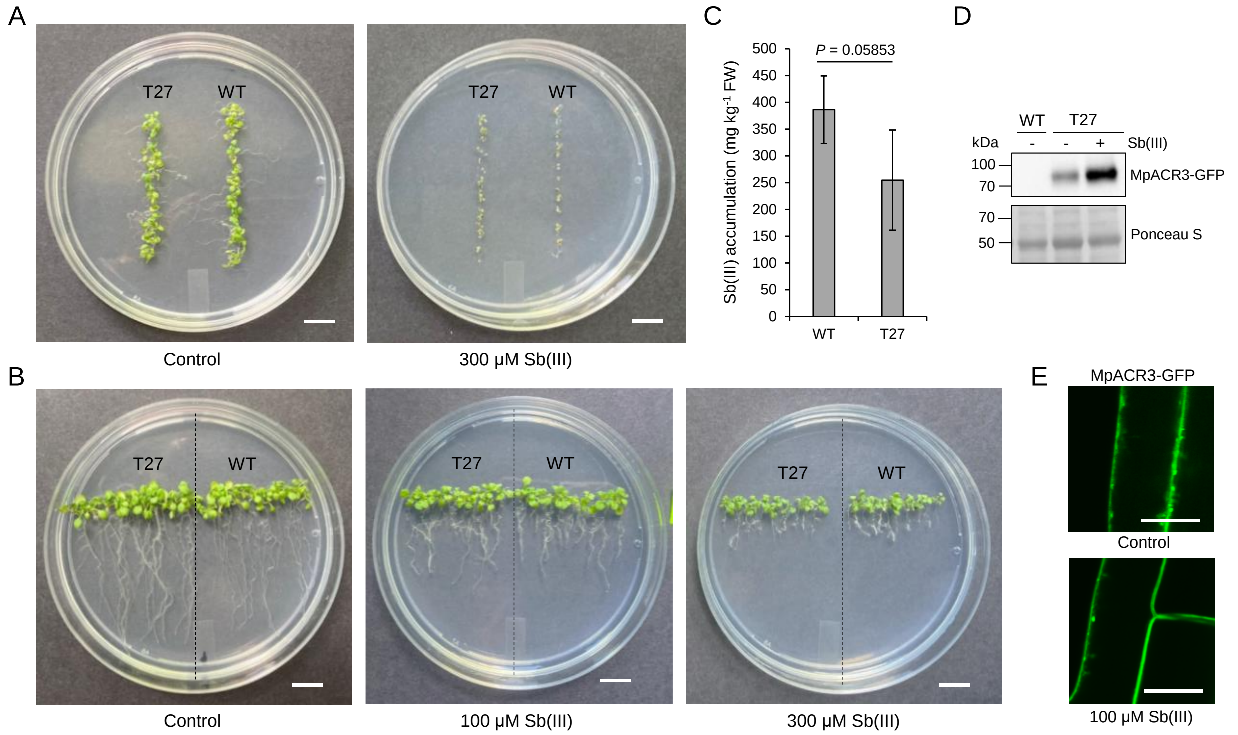


**Supplementary Figure S12.** Effect of Mp*ACR3* overexpression on Sb(**III**) tolerance in *A. thaliana* (supports Fig. 3). **A** and **B)** *MpACR3* overexpression did not improve *A. thaliana* growth in the presence of Sb(**III**). **A)** Germination of wild-type (WT) and transgenic T27 (*35S:*Mp*ACR3-GFP*) *A. thaliana* seeds in the absence or presence of 300 μM Sb(**III**). Plates were photographed after 2 weeks of growth. Scale bars,1 cm. **B)** Comparative growth of 4-day-old WT and T27 seedlings grown in the absence or presence of Sb(**III**) for 7 days. Scale bars, 1 cm. **C)** Mp*ACR3* overexpression did not significantly decrease Sb(**III**) accumulation in *A. thaliana*. Sb(**III**) content in WT and T27 plants was measured after exposure to 100 μM Sb(**III**) for 2 weeks using an atomic absorption spectrometer. Error bars represent mean value ± standard deviation of mean ($n$ = 3). One-way ANOVA was used to calculate the $P$-value. FW, fresh weight. **D)** Western blot analysis of MpACR3-GFP protein levels in T27 transgenic plants grown in the absence or presence 100 μM Sb(**III**) for 2 weeks. Protein extract from WT *A. thaliana* was used as a negative control. Ponceau S staining served as a protein loading control. **E)** Partial loss of MpACR3-GFP intracellular signal in response to Sb(**III**) treatment. 4-day-old seedlings of transgenic T27 line were transferred to MS medium without (control) or with 100 μM Sb(**III**) and allowed to grow for an additional 1 day before confocal microscopic analysis. Scale bars, 16 μm.

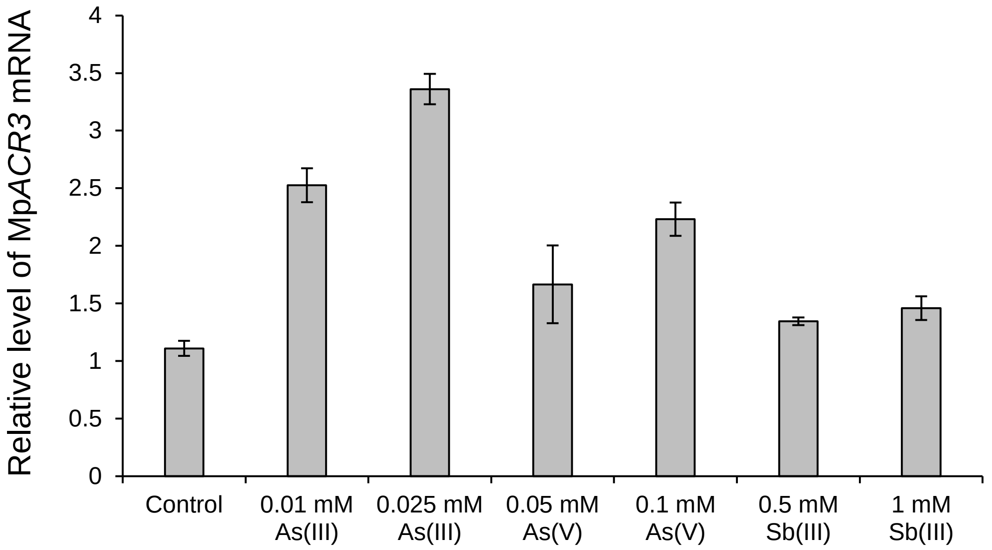


**Supplementary Figure S13.** *MpACR3* expression is increased in the presence of metalloids in *M. polymorpha* (supports Fig. 5). Total RNA was isolated from *M. polymorpha* Cam-1 gametophytes grown on solid 0.5x Gamborg B5 medium in the absence (control) or presence of indicated concentrations of As(III), As(V), or Sb(III) for two weeks. Mp*APT* (adenine phosphoribosyltransferase) was used as a reference gene to normalize expression data of Mp*ACR3*. Error bars represent mean value ± standard deviation of mean ($n$ = 3, three independent biological experiments using cDNA from at least three individual plants). One-way ANOVA was used to calculate the $P$-value. Control vs 0.01 mM As(III), $P$ = 0.011; control vs 0.025 mM As(III), $P$ = 0.0035; control vs 0.05 mM As(V), $P$ = 0.023; control vs 0.05 mM As(III), $P$ = 0.0025; control vs 0.5 mM Sb(III), $P$ = 0.023; control vs 1 mM Sb(III), $P$ = 0.023.

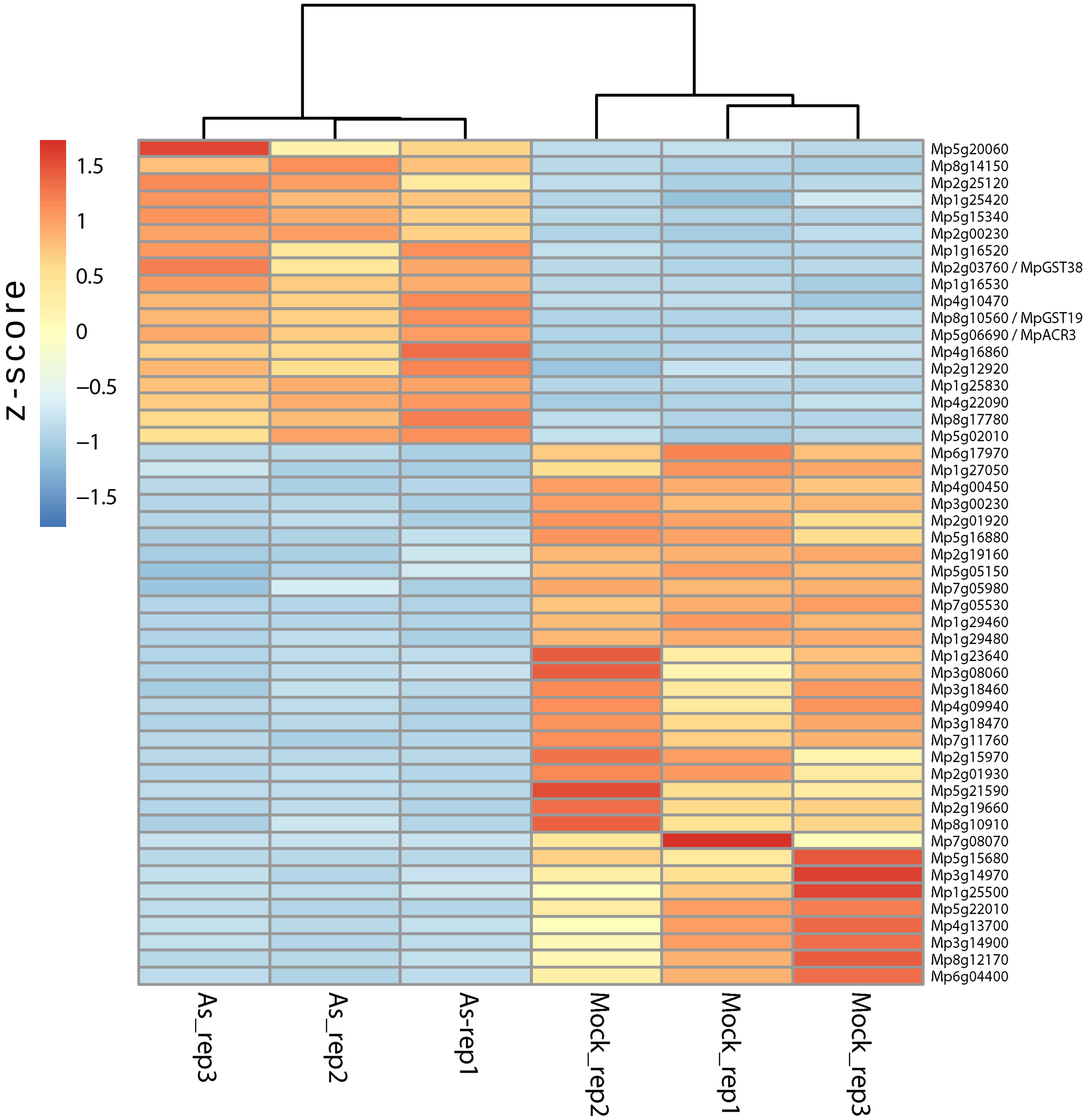


**Supplementary Figure S14.** Transcriptomic analysis of *M. polymorpha* plants treated with arsenic. Cam-1 plants were grown on solid 0.5× Gamborg B5 medium in the presence or absence of 2.5 µM As(III) for one week followed by RNA isolation and sequencing. Differentially expressed genes were sorted based on the lowest False Discovery Rate value generated with EdgeR. The top 50 candidates were clustered and plotted on a heat map with blue and red values indicating lower and higher transcript levels, respectively. Three biological repeats are shown for As(III) and control conditions.

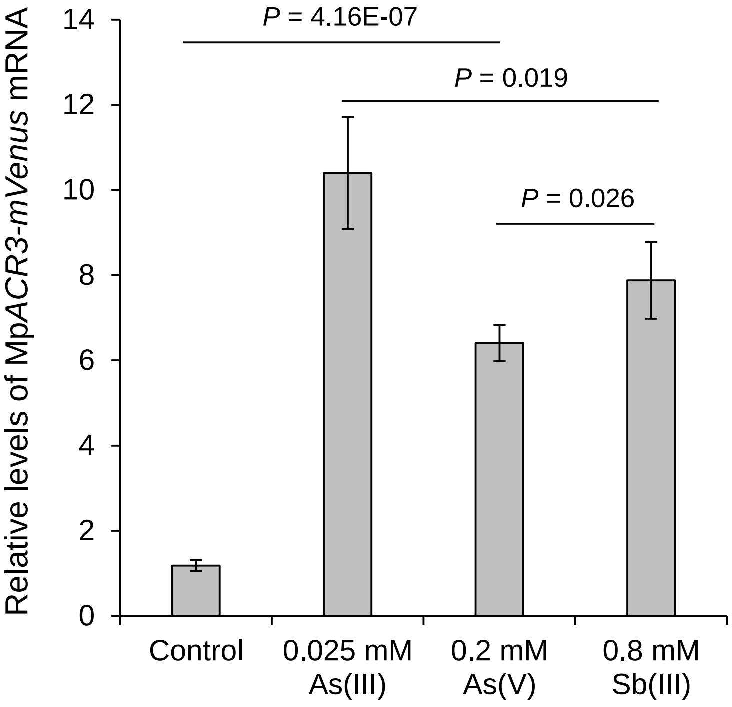


**Supplementary Figure S15.** Mp*ACR3-mVenus* expression is increased in the presence of metalloids in *M. polymorpha* (supports Fig. 5). Total RNA was isolated from transgenic *M. polymorpha* gametophytes expressing the *pro35Sx2:*Mp*ACR3-mVenus* transgene grown on solid media in the absence (control) or presence of indicated concentrations of As(III), As(V), or Sb(III) for 2 weeks. Mp*APT* (adenine phosphoribosyltransferase) was used as a reference gene to normalize expression data of Mp*ACR3-mVenus*. Error bars represent mean value ± standard deviation of mean ($n$ = 3). One-way ANOVA was used to calculate the *P*-value.

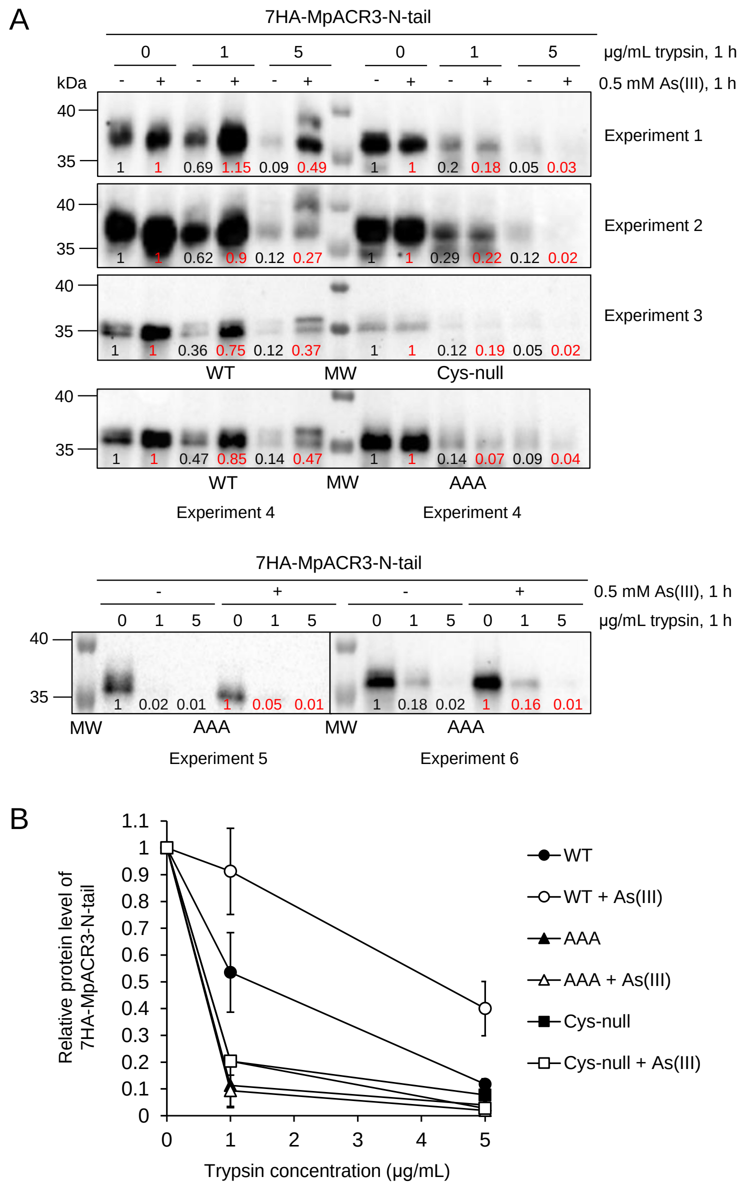


**Supplementary Figure S16.** Trypsin susceptibility of N-terminal cysteine-rich domain of MpACR3 (supports Fig. 7). **A)** Protein lysates were prepared from *acr3Δ* cells expressing the indicated variants of the N-terminal tail of MpACR3 (WT, Cys29A Cys47A Cys71A or AAA, Cys29A Cys47A Cys71A Cys73A Cys96A or Cys-null) that were treated or not with 0.5 mM As(III). The lysates were then exposed to increasing concentrations of trypsin for proteolysis, followed by Western blot analysis with anti-HA antibodies. The numbers in the Western blot lanes represent the relative protein level, determined from band intensity using ImageJ software and normalized relative to mock-treated protein samples. **B)** Quantification of 7HA-MpACR3-N-tail protein levels. Error bars represent mean value ± standard deviation of mean ($n$ = 4, WT; $n$ = 3, AAA and Cys-null variants). One-way ANOVA was used to calculate the $P$-value. WT vs WT + As(III), $0.001 < P < 0.01$; WT vs AAA, $0.007 < P < 0.025$; WT + As(III) vs AAA + As(III), $0.0005 < P < 0.001$; WT vs Cys-null, 1 µg/mL trypsin, $P = 0.018$; WT vs Cys-null, 5 µg/mL trypsin, not significant; WT + As(III) vs Cys-null + As(III), $0.0008 < P < 0.002$; AAA vs Cys-null, not significant.

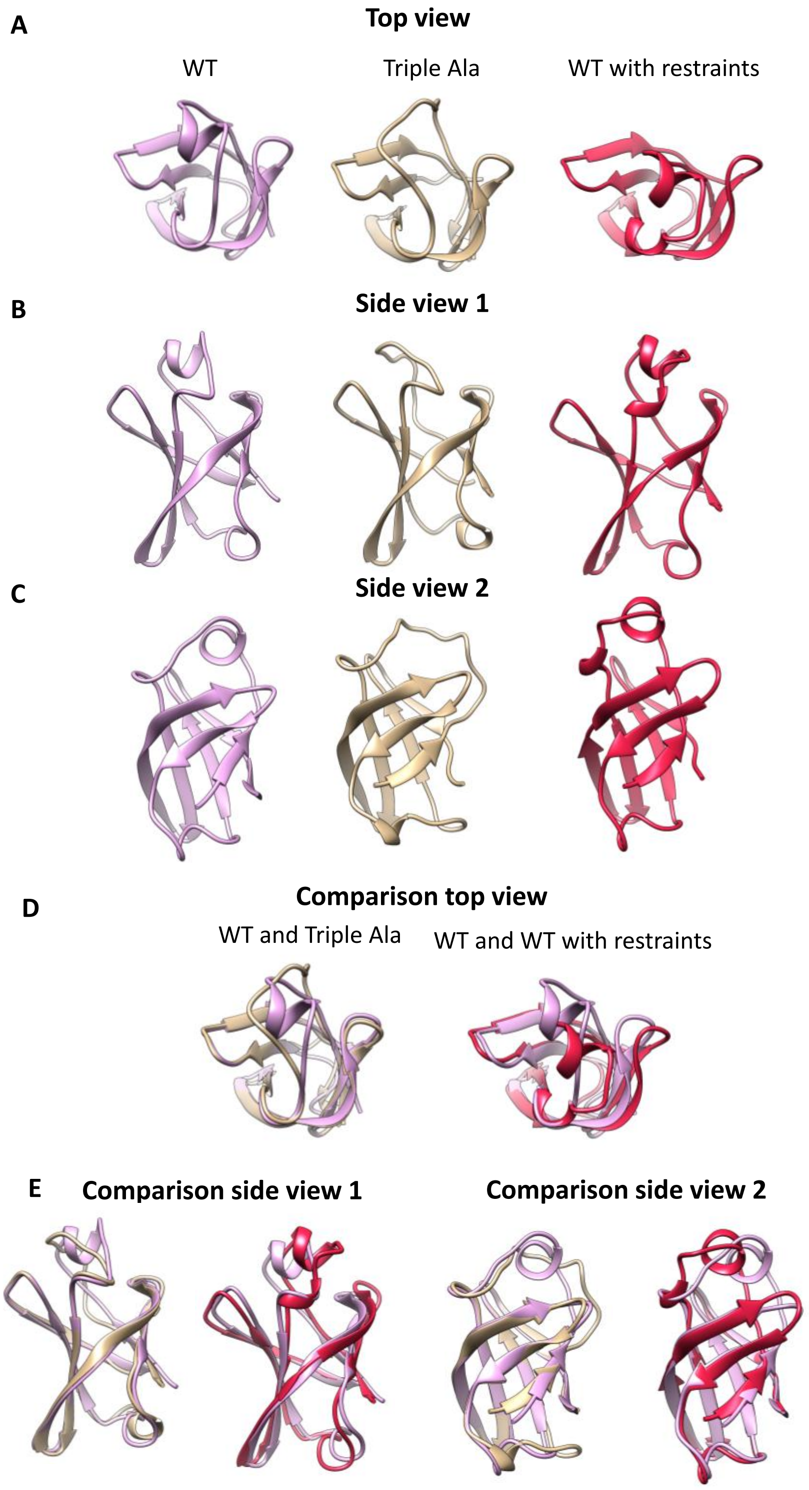


**Supplementary Figure S17.** Comparison of MD trajectory average structures of MpARC3 protein (supports Fig. 8). Triple Ala variant is in tan color, wild-type is in pink colour, and wild-type with the Cys-Cys restraints is in dark red colour.

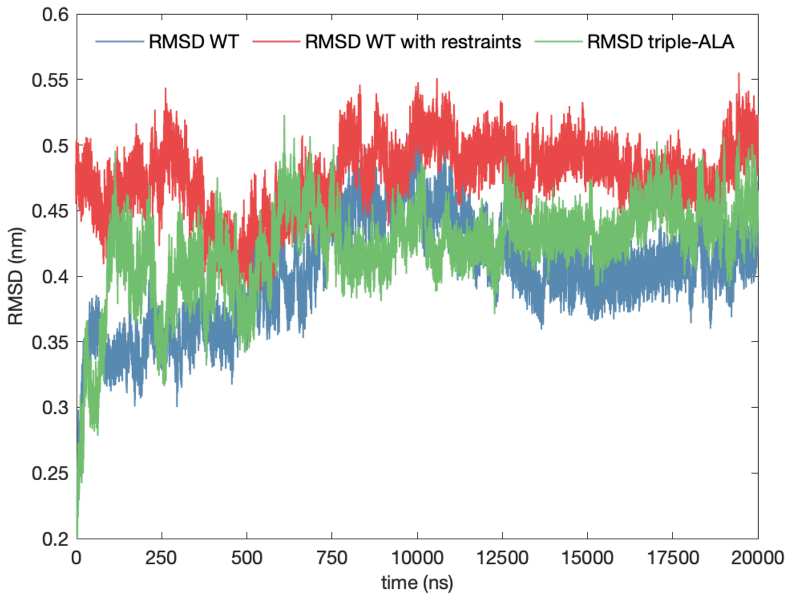


**Supplementary Figure S18.** Root mean square deviation (RMSD) of heavy atoms of the MpACR3 protein, Triple Ala variant, wild-type, and wild-type with the Cys-Cys restraints observed during 2000 ns all-atom MD trajectories (supports Fig. 8).

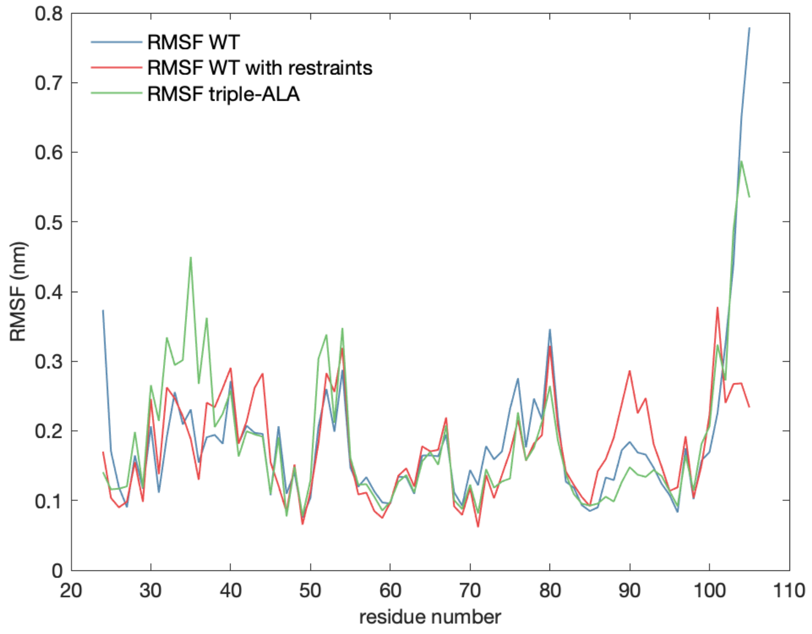


**Supplementary Figure S19.** Root mean square fluctuations (RMSF) per residue of the MpARC3 protein, Triple Ala mutant, wild-type, and wild-type with the Cys-Cys restraints observed during MD trajectories (supports Fig. 8).

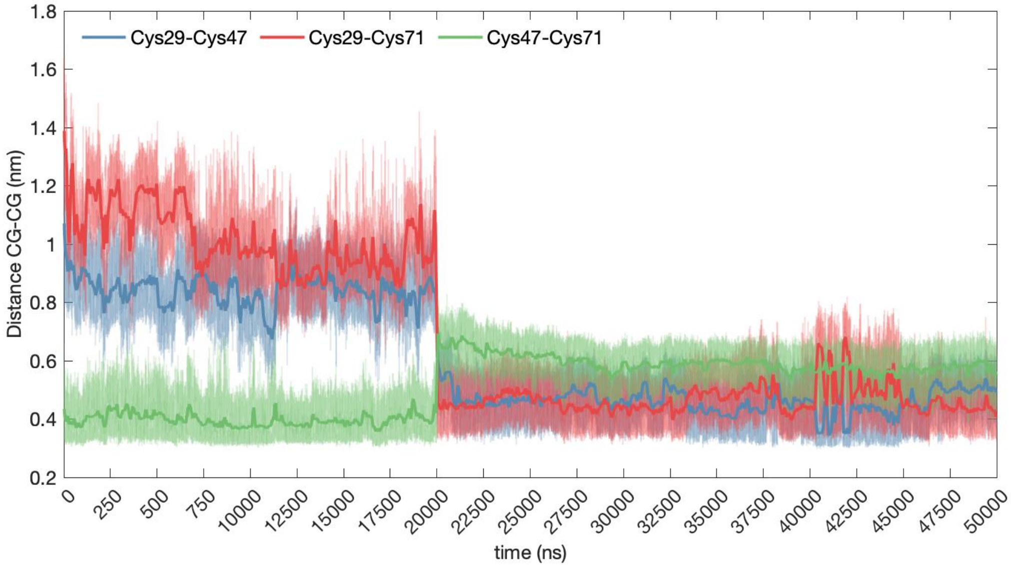


**Supplementary Figure S20.** Evolution of distances for SG-SG atoms for Cys29-Cys47, Cys29-Cys71, Cys47-Cys71 pairs (supports Fig. 8). First 2000 ns correspond to the MD simulation of the MpARC3 protein wild type system without restraints. The SG-SG distance restraints are switched on at 2000 ns and smoothly increased during the period 2000-3000 ns, and then were kept constant during the last 2000 ns.

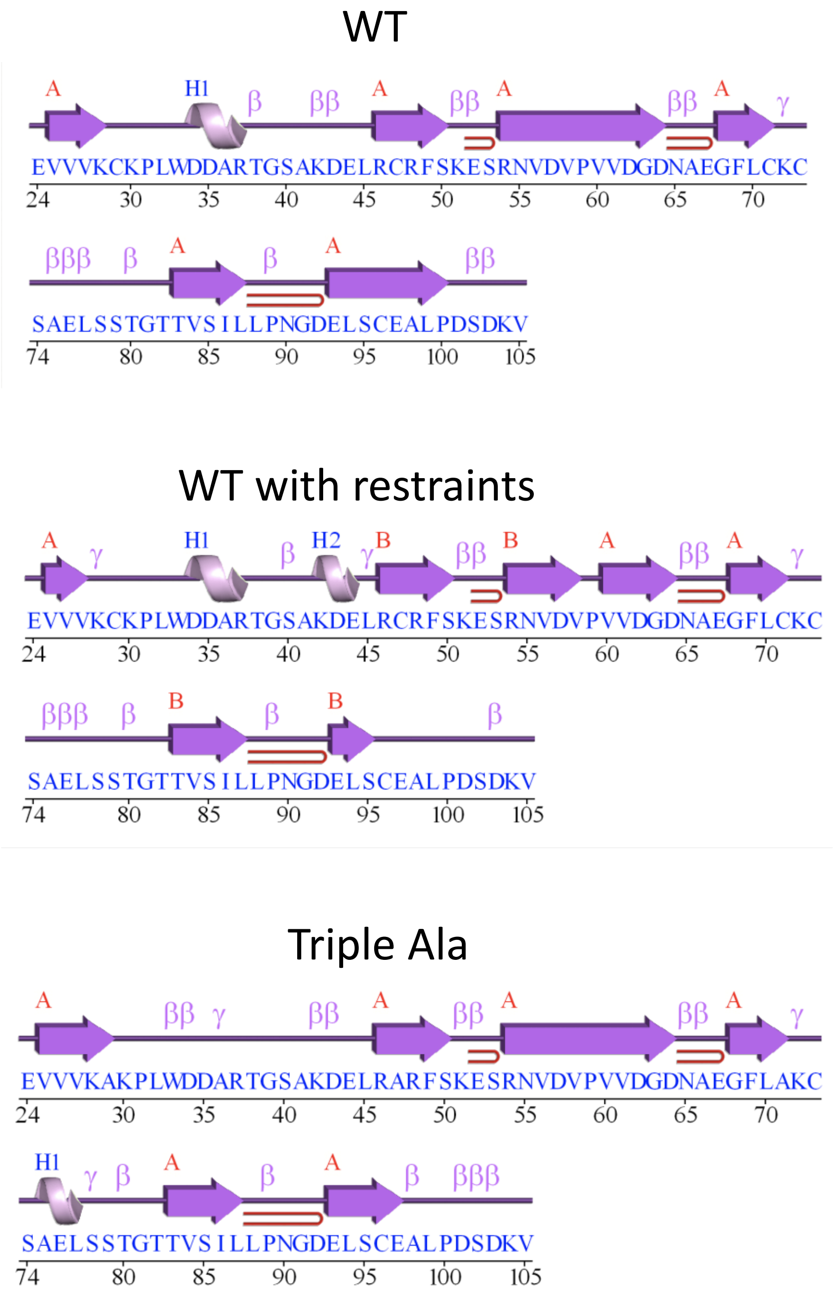


**Supplementary Figure S21.** Comparison of the secondary structures for wild-type, wild-type with the Cys-Cys restraints, and Triple Ala mutant obtained using the PDBsum webserver (supports Fig. 8).

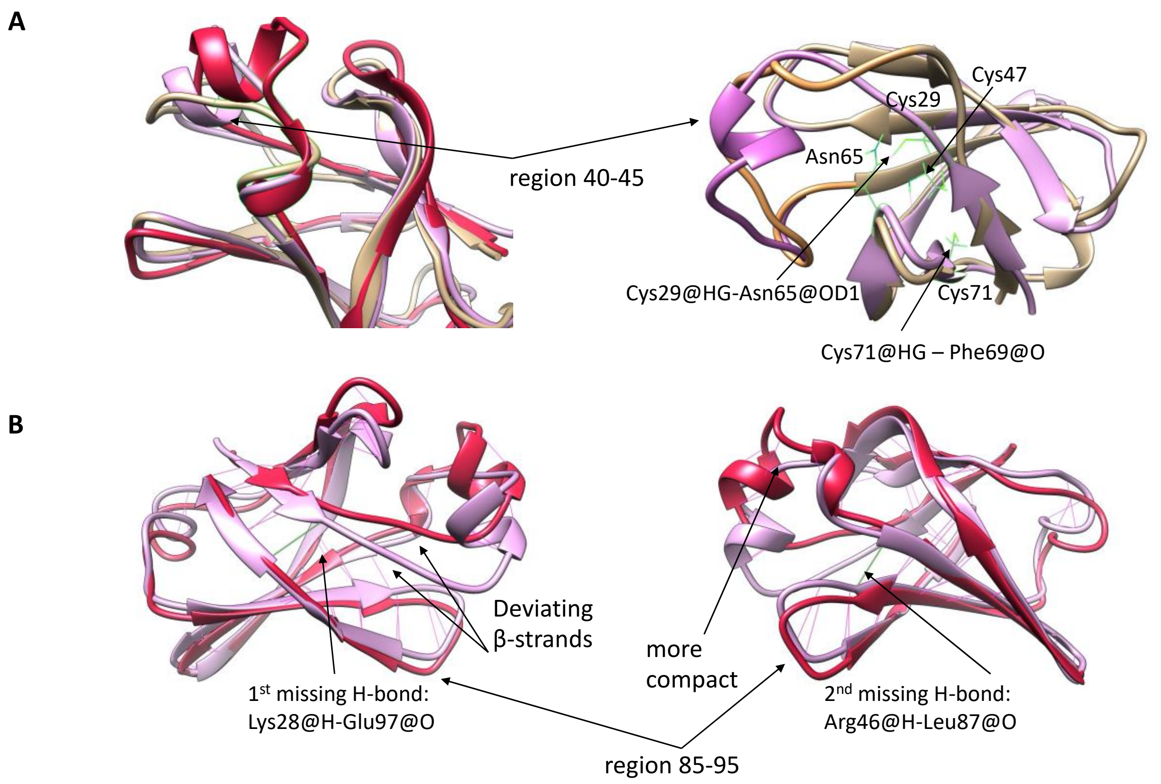


**Supplementary Figure S22.** Observed conformational differences in Triple Ala (tan coloured), wild-type (plum coloured), and wild-type with SG-SG restraints (red coloured) models during the 2μs MD trajectories (supports Fig. 8).

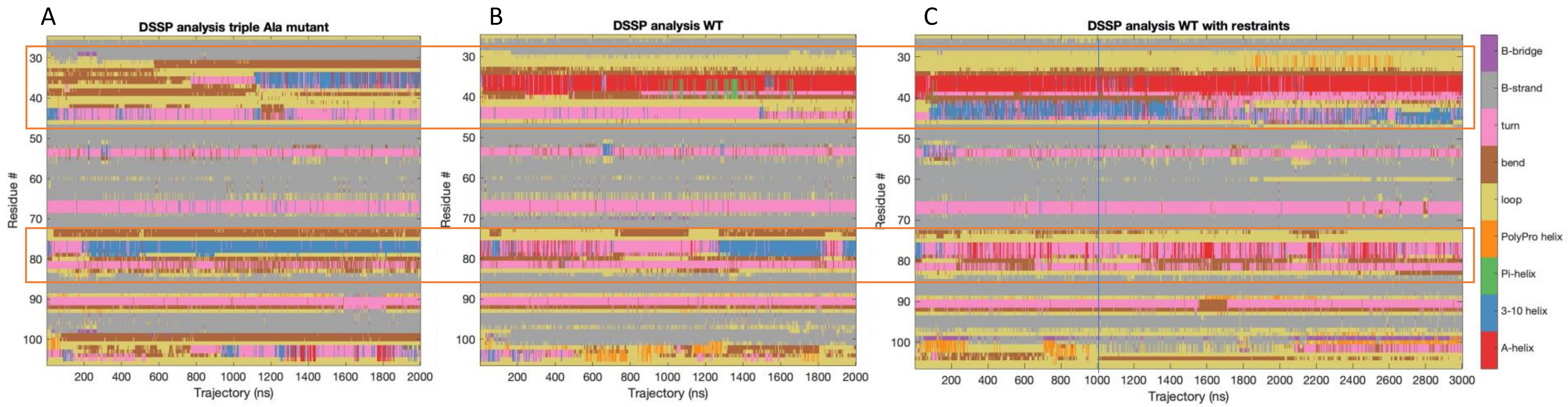


**Supplementary Figure S23.** Evolution of the secondary structure of MpARC3 protein, Triple Ala **A),** wild-type **B)**, and wild-type with the Cys-Cys restraints **C)** during MD trajectories (supports Fig. 8). For the restrained wild-type system trajectory, the Cys-Cys restraints were smoothly increased during the first 1000 ns marked with a blue vertical line (see Materials and methods), then the restraints were kept constant during the remaining 2000 ns. The colour-map to the right shows the protein secondary structure elements. The major differences in the secondary structure between Triple Ala mutant and wild-type systems include a. a. r. 32-45 and 75-85. For the region a. a. r. 32-45 we observe an appearance of a short $\alpha$-helical region (a. a. r. 35-40). When the restraints are switched on, the helical region extends to a. a. r. 32-45. For region a. a. r. 75-85, the 3-10 helical structure in the Triple Ala mutant is instead a turn in the wild-type system with restraints.

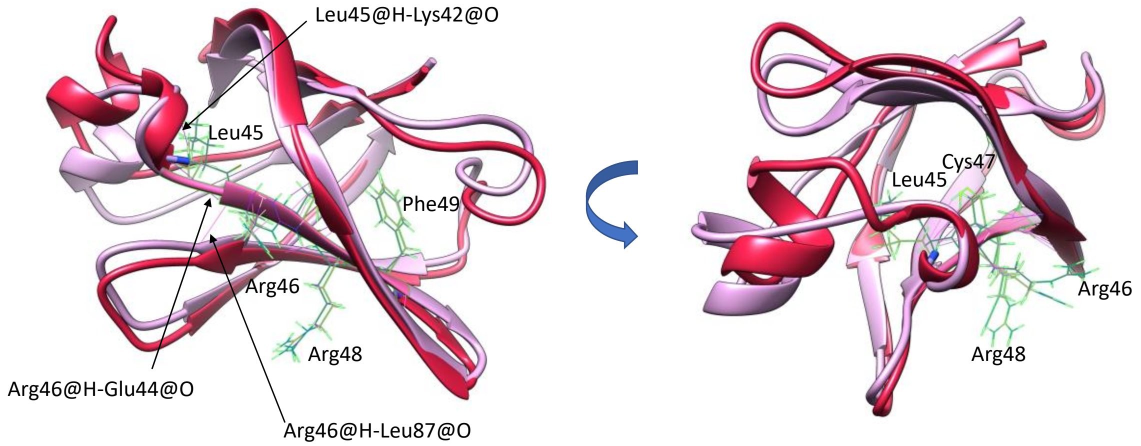


**Supplementary Figure S24.** Comparison of accessibility and orientation of Arg46 and Arg48 residues of the LRCRF motif (supports Fig. 8).

**Supplementary Table S1.** Plant ACR3 proteins analysed in silico in this study.

| Protein name | Organism | Taxonomic group | Database | Protein identifier |
|---|---|---|---|---|
| PsACR3 | *Picea sitchensis* | Gymnosperms | NCBI[1] | ADE76924.1 |
| TpACR3 | *Thuja plicata* | Gymnosperms | Phytozome[2] | Thupl.29379155s0008 |
| GbACR3 | *Ginkgo biloba* | Gymnosperms | Gigascience[3] | Gb_32402 |
| WmACR3 | *Welwitschia mirabilis* | Gymnosperms | Data Dryad[4] | W.mirabilis.22940 |
| GmACR3 | *Gnetum montanum* | Gymnosperms | Data Dryad[5] | TnS000269885t05 |
| PvACR3 | *Pteris vittata* | Ferns | NCBI | ADP20955.1 |
| CrACR3 | *Ceratopteris richardii* | Ferns | NCBI | KAH7279060.1 |
| AnACR3 | *Adiantum nelumboides* | Ferns | NCBI | MCO5594610.1 |
| SaACR3 | *Salvinia cucullata* | Ferns | Fernbase[6] | Sacu_v1.1_s0024.g008994 |
| AsACR3 | *Alsophila spinulosa* | Ferns | Fernbase | Aspi01Gene69197.t1 |
| MvACR3;1 | *Marsilea vestita* | Ferns | Fernbase | Mvestita_S10g02657-RA |
| MvACR3;2 | *Marsilea vestita* | Ferns | Fernbase | Mvestita_S10g02658-RA |
| DcACR3;1 | *Diphasiastrum complanatum* | Lycophytes | NCBI | KAJ7529772.1 |
| DcACR3;2 | *Diphasiastrum complanatum* | Lycophytes | NCBI | KAJ7529768.1 |
| ItACR3;1 | *Isoetes taiwanensis* | Lycophytes | Fernbase | Itaiw_v1_scaffold_14_t11603-RA |
| ItACR3;2 | *Isoetes taiwanensis* | Lycophytes | Fernbase | Itaiw_v1_scaffold_2_t02514-RA |
| SmACR3 | *Selaginella moellendorffii* | Lycophytes | NCBI | XP_024531516.1 |
| PpACR3 | *Physcomitrium patens* | Mosses | NCBI | XP_024362327.1 |
| CpACR3 | *Ceratodon purpureus* | Mosses | NCBI | KAG0559408.1 |
| SfACR3 | *Sphagnum fallax* | Mosses | NCBI | KAH8961229.1 |
| MpACR3 | *Marchantia polymorpha* | Liverworts | NCBI | OAE35577.1 |
| RfACR3 | *Riccia fluitans* | Liverworts | NCBI | KAL2645082.1 |
| AaACR3 | *Anthoceros agrestis* | Hornworts | Hornwort genomes[7] | AagrOXF_evm.model.utg000091l.433__434.1 |
| ApACR3 | *Anthoceros punctatus* | Hornworts | Hornwort genomes | Apun_evm.model.utg000254l.51.1 |
| MeACR3 | *Mesotaenium endlicherianum* SAG 12.97 | Charophytes | Phycocosm[8] | https://phycocosm.jgi.doe.gov/cgi-bin/dispGeneModel?db=Mesen1_1&id=7825 |
| ClACR3 | *Closterium sp.* NIES-68 | Charophytes | NCBI | GJP46584.1 |
| KnACR3;1 | *Klebsormidium nitens* NIES-2285 | Charophytes | NCBI | GAQ91548.1 |
| KnACR3;2 | *Klebsormidium nitens* NIES-2285 | Charophytes | NCBI | GAQ89538.1 |

| | | | | |
|---|---|---|---|---|
| CsACR3 | *Coccomyxa subellipsoidea* C-169 | Chlorophytes | Phycocosm | https://phycocosm.jgi.doe.gov/cgi-bin/dispGeneModel?db=Cosub3&id=1208183 |
| CnACR3;1 | *Coccomyxa elongate* SAG216_3B | Chlorophytes | NCBI | CAL8460937.1 |
| CnACR3;2 | *Coccomyxa elongate* SAG216_3B | Chlorophytes | NCBI | CAL8465694.1 |
| CnACR3;3 | *Coccomyxa elongate* SAG216_3B | Chlorophytes | NCBI | CAL8467475.1 |
| CvACR3;1 | *Coccomyxa viridis* | Chlorophytes | NCBI | CAK0787439.1 |
| CvACR3;2 | *Coccomyxa viridis* | Chlorophytes | NCBI | CAK0768907.1 |
| TsACR3 | *Trebouxia sp.* C0004 | Chlorophytes | NCBI | KAL0019817.1 |
| CoACR3 | *Chlorella ohadii* isolate 1 | Chlorophytes | NCBI | KAI7841530.1 |
| BbACR3 | *Botryococcus braunii* Showa | Chlorophytes | Phycocosm | https://phycocosm.jgi.doe.gov/cgi-bin/dispGeneModel?db=Botrbrau1&id=23661 |
| CeACR3 | *Chlamydomonas eustigma* NIES-2499 | Chlorophytes | NCBI | GAX84956.1 |
| CrACR3 | *Chloromonas remiasii* | Chlorophytes | Phycocosm | https://phycocosm.jgi.doe.gov/cgi-bin/dispGeneModel?db=Chlremi1&id=2447160 |
| LsACR3 | *Limnomonas spitsbergensis* CCryo_02-99_CH | Chlorophytes | Phycocosm | https://phycocosm.jgi.doe.gov/cgi-bin/dispGeneModel?db=Limspi1&id=589 dmel_g5956.t1 |
| SuACR3 | *Sanguina aurantia* | Chlorophytes | NCBI | MEW5305434.1 |
| CzACR3 | *Chromochloris zofingiensis* | Chlorophytes | Phycocosm | https://phycocosm.jgi.doe.gov/cgi-bin/dispGeneModel?db=Chrzof1&id=14366 |
| MmACR3 | *Monoraphidium minutum* isolate 26B-AM | Chlorophytes | NCBI | KAI8476375.1 |
| RsACR3 | *Raphidocelis subcapitata* NIES-35 | Chlorophytes | NCBI | GBF99582.1 |
| SoACR3 | *Scenedesmus obliquus* UTEX 3031 | Chlorophytes | NCBI | WIA08162.1 |
| EcACR3 | *Enallax costatus* CCAP 276/31 | Chlorophytes | Phycocosm | https://phycocosm.jgi.doe.gov/cgi-bin/dispGeneModel?db=Enacos1_1&id=6449963 |

| | | | | |
|---|---|---|---|---|
| FrACR3 | *Flechtneria rotunda* SEV3VF49 | Chlorophytes | Phycocosm | https://phycocosm.jgi.doe.gov/cgi-in/dispGeneModel?db=Flerot1_1&id=11394391 |

[1]www.ncbi.nlm.nih.gov
[2]phytozome-next.jgi.doe.gov
[3]gigadb.org/dataset/100613
[4]datadryad.org/stash/dataset/doi:10.5061/dryad.ht76hdrdr
[5]datadryad.org/stash/dataset/doi:10.5061/dryad.0vm37
[6]fernbase.org
[7]www.hornworts.uzh.ch/en/hornwort-genomes.html
[8]phycocosm.jgi.doe.gov

**Supplementary Table S2.** Oligonucleotides used in this work.

| Name | Sequence 5' to 3' |
|---|---|
| **Mutagenesis:** | |
| C29A-fw | GGTGAAGTTGTTGTGAAGGCTAAGCCCTTGTGGGACGA |
| C29A-rv | TCGTCCCACAAGGGCTTAGCCTTCACAACAACTTCACC |
| L45A-fw | GAGCGCGAAGGATGAGGCGAGGTGTCGGTTCTCC |
| L45A-rv | GGAGAACCGACACCTCGCCTCATCCTTCGCGCTC |
| L45F-fw | GAGCGCGAAGGATGAGTTCAGGTGTCGGTT |
| L45F-rv | AACCGACACCTGAACTCATCCTTCGCGCTC |
| L45V-fw | GCGCGAAGGATGAGGTGAGGTGTCGGTTC |
| L45V-rv | GAACCGACACCTCACCTCATCCTTCGCGC |
| R46A-fw | CGCGAAGGATGAGTTGGCGTGTCGGTTCTCCAAG |
| R46A-rv | CTTGGAGAACCGACACGCCAACTCATCCTTCGCG |
| R46N-fw | AGGGAGCGCGAAGGATGAGTTGAATTGTCGGTTCTCC |
| R46N-rv | GGAGAACCGACAATTCAACTCATCCTTCGCGCTCCCT |
| R46S-fw | CGAAGGATGAGTTGAGCTGTCGGTTCTCCAAGG |
| R46S-rv | CCTTGGAGAACCGACAGCTCAACTCATCCTTCG |
| R46T-fw | GCGAAGGATGAGTTGACGTGTCGGTTCTCCAAG |
| R46T-rv | CTTGGAGAACCGACACGTCAACTCATCCTTCGC |
| R46V-fw | GCGCGAAGGATGAGTTGGTGTGTCGGTTCTCCAAGG |
| R46V-rv | CCTTGGAGAACCGACACACCAACTCATCCTTCGCGC |
| C47A-fw | GCGAAGGATGAGTTGAGGGCTCGGTTCTCCAAGGAATC |
| C47A-rv | GATTCCTTGGAGAACCGAGCCCTCAACTCATCCTTCGC |
| R48A-fw | AAGGATGAGTTGAGGTGTGCGTTCTCCAAGGAATCGAG |
| R48A-rv | CTCGATTCCTTGGAGAACGCACACCTCAACTCATCCTT |
| R48D-fw | GCGAAGGATGAGTTGAGGTGTGATTTCTCCAAGGAATCGAGGAAT |
| R48D-rv | ATTCCTCGATTCCTTGGAGAAATCACACCTCAACTCATCCTTCGC |
| F49A-fw | GATGAGTTGAGGTGTCGGGCCTCCAAGGAATCGAGGAA |
| F49A-rv | TTCCTCGATTCCTTGGAGGCCCGACACCTCAACTCATC |
| F49I-fw | TGAGTTGAGGTGTCGGATCTCCAAGGAATCGAG |
| F49I-rv | CTCGATTCCTTGGAGATCCGACACCTCAACTCA |
| F49L-fw | ATGAGTTGAGGTGTCGGTTATCCAAGGAATCGAGG |
| F49L-rv | CCTCGATTCCTTGGATAACCGACACCTCAACTCAT |
| F49V-fw | TGAGTTGAGGTGTCGGGTCTCCAAGGAATCGAG |
| F49V-rv | CTCGATTCCTTGGAGACCCGACACCTCAACTCA |
| C71A-fw | CAATGCTGAAGGGTTTCTGGCTAAATGTAGTGCCGAGCTT |
| C71A-rv | AAGCTCGGCACTACATTTAGCCAGAAACCCTTCAGCATTG |
| C73A-fw | ACAATGCTGAAGGGTTTCTGTGTAAAGCTAGTGCCGAGCTTT |
| C73A-rv | AAAGCTCGGCACTAGCTTTACACAGAAACCCTTCAGCATTGT |
| C96A-fw | GCGATGAGCTCTCGGCTGAAGCCCTACCGG |
| C96A-rv | CCGGTAGGGCTTCAGCCGAGAGCTCATCGC |
| C236A-fw | GGTGGGAATCGCTCGCGCCATTGCCATGGTACTT |
| C236A-rv | AAGTACCATGGCAATGGCGCGAGCGATTCCCACC |
| E407A-fw | CAGCCAGTAACAACTTTGCGCTGGCCATAGCTG |
| E407A-rv | CAGCTATGGCCAGCGCAAAGTTGTTACTGGCTG |
| E434A-fw | CGGGCCTCTCATCGCGGTTCCCGTTCTAT |
| E434A-rv | ATAGAACGGGAACCGCGATGAGAGGCCCG |
| L45A-C47A-fw | GGGAGCGCGAAGGATGAGGCGAGGGCTCGGTTCTCCAAGGAATC |
| L45A-C47A-rv | GATTCCTTGGAGAACCGAGCCCTCGCCTCATCCTTCGCGCTCCC |

| | |
|---|---|
| R46A-C47A-fw | GAGCGCGAAGGATGAGTTGGCGGCTCGGTTCTCCAAGGAATCG |
| R46A-C47A-rv | CGATTCCTTGGAGAACCGAGCCGCCAACTCATCCTTCGCGCTC |
| C47A-R48A-fw | CGCGAAGGATGAGTTGAGGGCTGCGTTCTCCAAGGAATCGAGG |
| C47A-R48A-rv | CCTCGATTCCTTGGAGAACGCAGCCCTCAACTCATCCTTCGCG |
| L45A-F49A-fw | CGCGAAGGATGAGTTGAGGGCTGCGTTCTCCAAGGAATCGAGG |
| L45A-F49A-rv | CTCGATTCCTTGGAGGCCCGACACCTCGCCTCATCCTTCGCGCT |
| R46A-F49A-fw | AGCGCGAAGGATGAGGCGAGGTGTCGGGCCTCCAAGGAATCGAG |
| R46A-F49A-rv | CCTCGATTCCTTGGAGGCCCGACACGCCAACTCATCCTTCGCG |
| R48A-F49A-fw | CGAAGGATGAGTTGAGGTGTGCGGCCTCCAAGGAATCGAGGAATG |
| R48A-F49A-rv | CATTCCTCGATTCCTTGGAGGCCGCACACCTCAACTCATCCTTCG |
| **Gene cloning** | |
| BamHI-MpACR3-fw | TAGAACTAGTGGATCCATGAGGGGCTCGGAGATG |
| SalI-MpACR3-rv | ACATGTCGAGGTCGACAGCTTGTTCTTTTGAGAGCC |
| BamHI-PpACR3-fw | TAGAACTAGTGGATCCATGGCAACGCGTCACGAGA |
| SalI-PpACR3-rv | ACATGTCGAGGTCGACGAAGAATTTCTTCCTAAAAAAGAGA |
| BamHI-PvACR3-fw | TAGAACTAGTGGATCCATGGAAAATTCATCCGCTGAAAG |
| SalI-PvACR3-rv | ACATGTCGAGGTCGACAACAGATGGACCCTTTCTTTGG |
| Δ27-98-MpACR3-fw | TTGTTGTGGCCCTACCGGACTCGGAC |
| Δ27-98-MpACR3-rv | GTAGGGCCACAACAACTTCACCAGCTTC |
| N-tail-MpACR3-fw | GTCGACCTCGACATGTCT |
| N-tail-MpACR3-rv | ATCGGTGCGATTCACCAT |
| N-tail-Δ-ScACR3-fw | AAGGGCGTGGTTACAATACTGACTACGATCAAGTC |
| N-tail-Δ-ScACR3-rv | CATGTCGAGGTCGACATTGTTCCATATATAATATGGTT |
| N-tail-ScACR3-fw | GTCGACCTCGACATGTCTAAAGG |
| N-tail-ScACR3-rv | TGTAACCACGCCCTTGACCT |
| N-tail-Δ-MpACR3-fw | GTGAATCGCACCGATGCTGGTCAAATCTACAAACAGC |
| N-tail-Δ-MpACR3-rv | CATGTCGAGGTCGACAGCTTGTTCTTTTGAGAGCC |
| N-tail-MpACR3-pGREG-fw | GAATTCGATAATCAAGCTTATCGATACCGTCGACAATGAGAGGTTCTGAAATGACTGAAATGG |
| N-tail-MpACR3-pGREG-rv | GCGTGACATAACTAATTACATGACTCGAGGTCGACTTAGTCGACTCTGTCCAGCAAAGACAATTGT |
| MpACR3-GUSGFP-fw | GAGAACACGGGGGACTCTAGATGAGGGGCTCGGAGATGACTGAG |
| MpACR3-GUSGFP-rv | GATCGAATTGATCCTCTAGAGCTTGTTCTTTTGAGAGCCATTTCC |
| **qPCR:** | |
| MpACR3q-fw | GTGCCCTTGCTGCTTTACTT |
| MpACR3q-rv | GTTACTGGCTGCAGTGAAGG |
| MpAPT1q-fw | CGAAAGCCCAAGAAGCTACC |
| MpAPT1q-rv | GTACCCCCGGTTGCAATAAG |
| mVENUSq-fw | GATGAAGCAGCACGACTTC |
| mVENUSq-rv | CGTCCTTGAAGAAGATGGT |

**Supplementary Table S3.** Expression plasmids used in this work.

| **Name** | **Description** | **Source** |
|---|---|---|
| **Yeast expression:** | | |
| pUG35 | MET17 promoter, yeGFP, CEN, URA3, AmpR | Lab collection |
| pScACR3 | $_{pro}$*MET17:ScACR3-yeGFP* in pUG35 | Lab collection |
| pMpACR3 | $_{pro}$*MET17:*Mp*ACR3-yeGFP* in pUG35 | This study |
| pMpACR3-C29A | $_{pro}$*MET17:*Mp*ACR3-C29A-yeGFP* in pUG35 | This study |
| pMpACR3-L45A | $_{pro}$*MET17:*Mp*ACR3-L45A-yeGFP* in pUG35 | This study |
| pMpACR3-L45F | $_{pro}$*MET17:*Mp*ACR3-L45F-yeGFP* in pUG35 | This study |
| pMpACR3-L45V | $_{pro}$*MET17:*Mp*ACR3-L45V-yeGFP* in pUG35 | This study |
| pMpACR3-R46A | $_{pro}$*MET17:*Mp*ACR3-R46A-yeGFP* in pUG35 | This study |
| pMpACR3-R46N | $_{pro}$*MET17:*Mp*ACR3-R46N-yeGFP* in pUG35 | This study |
| pMpACR3-R46S | $_{pro}$*MET17:*Mp*ACR3-R46S-yeGFP* in pUG35 | This study |
| pMpACR3-R46T | $_{pro}$*MET17:*Mp*ACR3-R46T-yeGFP* in pUG35 | This study |
| pMpACR3-R46V | $_{pro}$*MET17:*Mp*ACR3-R46V-yeGFP* in pUG35 | This study |
| pMpACR3-C47A | $_{pro}$*MET17:*Mp*ACR3-C47A-yeGFP* in pUG35 | This study |
| pMpACR3-R48A | $_{pro}$*MET17:*Mp*ACR3-R48A-yeGFP* in pUG35 | This study |
| pMpACR3-R48D | $_{pro}$*MET17:*Mp*ACR3-R48D-yeGFP* in pUG35 | This study |
| pMpACR3-F49A | $_{pro}$*MET17:*Mp*ACR3-F49A-yeGFP* in pUG35 | This study |
| pMpACR3-F49I | $_{pro}$*MET17:*Mp*ACR3-F49I-yeGFP* in pUG35 | This study |
| pMpACR3-F49L | $_{pro}$*MET17:*Mp*ACR3-F49L-yeGFP* in pUG35 | This study |
| pMpACR3-F49V | $_{pro}$*MET17:*Mp*ACR3-F49V-yeGFP* in pUG35 | This study |
| pMpACR3-C71A | $_{pro}$*MET17:*Mp*ACR3-C71A-yeGFP* in pUG35 | This study |
| pMpACR3-C73A | $_{pro}$*MET17:*Mp*ACR3-C73A-yeGFP* in pUG35 | This study |
| pMpACR3-C96A | $_{pro}$*MET17:*Mp*ACR3-C96A-yeGFP* in pUG35 | This study |
| pMpACR3-C236A | $_{pro}$*MET17:*Mp*ACR3-C236A-yeGFP* in pUG35 | This study |
| pMpACR3-E407A | $_{pro}$*MET17:*Mp*ACR3-E407A-yeGFP* in pUG35 | This study |
| pMpACR3-E434A | $_{pro}$*MET17:*Mp*ACR3-E434A-yeGFP* in pUG35 | This study |
| pMpACR3-L45A,C47A | $_{pro}$*MET17:*Mp*ACR3-L45A,C47A-yeGFP* in pUG35 | This study |
| pMpACR3-L45A,F49A | $_{pro}$*MET17:*Mp*ACR3-L45A,F49A-yeGFP* in pUG35 | This study |
| pMpACR3-R46A,C47A | $_{pro}$*MET17:*Mp*ACR3-R46A,C47A-yeGFP* in pUG35 | This study |
| pMpACR3-R46A F49A | $_{pro}$*MET17:*Mp*ACR3-R46A,F49A-yeGFP* in pUG35 | This study |
| pMpACR3-C47A,R48A | $_{pro}$*MET17:*Mp*ACR3-C47A,R48A-yeGFP* in pUG35 | This study |
| pMpACR3-R48A,F49A | $_{pro}$*MET17:*Mp*ACR3-R48A,F49A-yeGFP* in pUG35 | This study |
| pMpACR3-AAA | $_{pro}$*MET17:*Mp*ACR3-C29A,C47A,C71A-yeGFP* in pUG35 | This study |
| pN$_{Sc}$-MpACR3 | $_{pro}$*MET17:ScACR3*$_{1\text{-}22aa}$*-MpACR3-*$_{111\text{-}457aa}$*-yeGFP* in pUG35 | This study |
| pN$_{Mp}$-ScACR3 | $_{pro}$*MET17:*Mp*ACR3*$_{1\text{-}110aa}$*-ScACR3*$_{23\text{-}406aa}$*-yeGFP* in pUG35 | This study |
| pMpACR3-Δ27-98 | $_{pro}$*MET17:*Mp*ACR3-Δ27-98aa-yeGFP* in pUG35 | This study |
| pPpACR3 | $_{pro}$*MET17:PpACR3-yeGFP* in pUG35 | This study |
| pPvACR3 | $_{pro}$*MET17:PvACR3-yeGFP* in pUG35 | This study |
| pGREG535 | *GAL1* promoter, 7×HA, *CEN*, *LEU2*, AmpR | EUROSCARF |

| | | |
|---|---|---|
| p7HA-MpACR3-N-tail | $_{pro}$*GAL1:7xHA*-Mp*ACR3*$_{1\text{-}123aa}$ in pGREG535 | This study |
| p7HA-MpACR3-AAA-N-tail | $_{pro}$*GAL1:7xHA* -Mp*ACR3*$_{1\text{-}123aa}$-*C29A,C47A,C71A* in pGREG535 | This study |
| p7HA-MpACR3-Cys-null-N-tail | $_{pro}$*GAL1:7xHA*-Mp*ACR3*$_{1\text{-}123aa}$-*C29A,C47A,C71A,C73A,C96A* in pGREG535 | This study |
| ***E. coli*** **expression:** | | |
| pGEX4T-1 | tac promoter, GST, lacI, AmpR | BioCat GmbH |
| pGST-MpACR3-N-tail | $_{pro}$*tac:GST*-Mp*ACR3*$_{1\text{-}123aa}$ in pGEX4T-1 | This study |
| pGST-MpACR3-AAA-N-tail | $_{pro}$*tac:GST*-Mp*ACR3*$_{1\text{-}123aa}$ -*C29A,C47A,C71A* in pGEX4T-1 | This study |
| pGST-MpACR3-Cys-null-N-tail | $_{pro}$*tac:GST*-Mp*ACR3*$_{1\text{-}123aa}$ -*C29A,C47A,C71A,C73A,C96A* in pGEX4T-1 | This study |
| **Plant expression:** | | |
| pBy10-MpACR3 | $_{pro}$*35Sx2:*Mp*ACR3-mVenus,* HygR, KanR | This study |
| pBy10-MpACR3-C47A | $_{pro}$*35Sx2:*Mp*ACR3-C47A-mVenus,* HygR, KanR | This study |
| pBy10-MpACR3-R46A,C47A | $_{pro}$*35Sx2:*Mp*ACR3-R46A,C47A-mVenus*, HygR, KanR | This study |
| pCsA-MpACR3-ST-mScarlet | $_{pro}$*35Sx2:*Mp*ACR3-mVenus,* $_{pro}$*UbE2:52aaST-mScarlet*, HygR, SpecR | This study |
| pGFPGUSPlus | CaMV 35S promoter, EGFP, GUSplus™, HygR, KanR | Addgene |
| pGFPGUSPlus-MpACR3 | $_{pro}$*35S:*Mp*ACR3-EGFP* in pGFPGUSPlus | This study |
